# Local Disordered Region Sampling (LDRS) for Ensemble Modeling of Proteins with Experimentally Undetermined or Low Confidence Prediction Segments

**DOI:** 10.1101/2023.07.25.550520

**Authors:** Zi Hao Liu, João M.C. Teixeira, Oufan Zhang, Thomas E. Tsangaris, Jie Li, Claudiu C. Gradinaru, Teresa Head-Gordon, Julie D. Forman-Kay

## Abstract

**SUMMARY:** The Local Disordered Region Sampling (LDRS, pronounced *loaders*) tool, developed for the IDPConformerGenerator platform (Teixeira *et al*. 2022), provides a method for generating all-atom conformations of intrinsically disordered regions (IDRs) at N- and C-termini of and in loops or linkers between folded regions of an existing protein structure. These disordered elements often lead to missing coordinates in experimental structures or low confidence in predicted structures. Requiring only a pre-existing PDB structure of the protein with missing coordinates or with predicted confidence scores and its full-length primary sequence, LDRS will automatically generate physically meaningful conformational ensembles of the missing flexible regions to complete the full-length protein. The capabilities of the LDRS tool of IDPConformerGenerator include modeling phosphorylation sites using enhanced Monte Carlo Side Chain Entropy (MC-SCE) (Bhowmick and Head-Gordon 2015), transmembrane proteins within an all-atom bilayer, and multi-chain complexes. The modeling capacity of LDRS capitalizes on the modularity, ability to be used as a library and via command-line, and computational speed of the IDPConformerGenerator platform.

**AVAILABILITY AND IMPLEMENTATION:** The LDRS module is part of the IDPConformerGenerator modeling suite, which can be downloaded from GitHub at https://github.com/julie-forman-kay-lab/IDPConformerGenerator. IDPConformerGenerator is written in Python and works on Linux, Microsoft Windows, and Mac OS versions that support DSSP. Users can utilize LDRS’s Python API for scripting the same way they can use any part of IDPConformerGenerator’s API, by importing functions from the ‘idpconfgen.ldrs_helper’ library. Otherwise, LDRS can be used as a command line interface application within IDPConformerGenerator. Full documentation is available within the command-line interface (CLI) as well as on IDPConformerGenerator’s official documentation pages (https://idpconformergenerator.readthedocs.io/en/latest/).

**CONTACT:** For support with LDRS please contact Zi Hao (Nemo) Liu via or submit an issue in the IDPConformerGenerator repository on GitHub (https://github.com/julie-forman-kay-lab/IDPConformerGenerator/issues).

**SUPPLEMENTARY INFORMATION:** The supplementary information document contains, or links to, all the conformer ensembles generated for this publication, the generalized Python scripts using the LDRS Python API, figures of detailed methods, fractional secondary structure information, torsion angle sampling, and the time required to generate the different protein cases.

## MAIN

Atomistic structures of proteins, which are computational models of real proteins, have been a central goal of the field of structural biology to generate hypotheses and deepen our understanding of the relationship between protein structure and function. Experimental structures determined by X-ray crystallography, NMR spectroscopy and cryo-electron microscopy (cryoEM) have provided incredible structural insights (Dokholyan 2020; Burley *et al*. 2022). Most recently, accurate protein structure predictions have become available using machine learning methods such as AlphaFold (Jumper *et al*. 2021), RoseTTAFold (Baek *et al*. 2021), and ESMFold (Lin *et al*. 2023).

Intrinsically disordered protein regions (IDRs) are not visible by X-ray crystallography and cryoEM because data averaging due to conformational heterogeneity leads to missing electron density and, hence, missing coordinates in the final models (Villarreal and Stewart 2014; Djinovic-Carugo and Carugo 2015; Nwanochie and Uversky 2019). Computational predicted structures contain coordinates for IDRs, but these coordinates generally have low confidence predictions and are not likely representative (Ruff and Pappu 2021). Since about 30% of residues within the human proteome are expected to be within IDRs (Tsang *et al*. 2020), obtaining structural insights from ensemble models of these flexible regions has become a focus.

A variety of computational sampling methods are available to model single chain intrinsically disordered protein (IDPs) ensembles, including IDPConformerGenerator (Teixeira *et al*. 2022), Flexible-meccano (Ozenne *et al*. 2012), TraDES (Feldman and Hogue 2000, 2002), and others (Shrestha, Smith and Petridis 2021; Karamanos, Kalverda and Radford 2022). FastFloppyTail (Ferrie and Petersson 2020) can be used to generate IDPs or IDRs at the N- and C-termini of a folded domain. Currently, though, there is no easy-to-use modular method of modeling conformational ensembles of (i) IDRs between two folded domains within an experimental structure, (ii) IDRs representing low confidence regions of predicted structures, (iii) IDRs within transmembrane proteins, or (iv) IDRs found in dynamic protein complexes. While cyclic coordinate descent (CCD) and kinematic closure methods (KIC) (Canutescu and Dunbrack 2003; Boomsma and Hamelryck 2005; Stein and Kortemme 2013; O’Donnell, Robert and Cazals 2022) are able to model all-atom missing protein regions (i.e., breaks in the protein chain), these approaches are limited to 12 residues. In addition, existing “loop” closure methods such as CCD do not rely on statistical relationships between backbone torsion angles and protein sequence, resulting in the lack of sampling of realistic fractional secondary structure.

Here, we present the Local Disordered Region Sampling (LDRS) tool, to model all-atom sidechain inclusive ensembles of N-terminal and C-terminal IDRs (N-IDRs and C-IDRs, respectively), as well as IDRs between folded elements (L-IDRs, for loops or linkers). Because the knowledge-based sampling of IDPConformerGenerator takes into consideration the statistical relationship between protein sequence and backbone torsion angles along with other chain geometry, the resulting IDR conformer ensembles are expected to be representative, based on observed agreement between IDPConformerGenerator ensemble structural properties and experiment (Teixeira *et al*. 2022). LDRS has been developed as a modular command-line interface and Python library within the IDPConformerGenerator platform (Teixeira *et al*. 2022), exploiting the flexibility and modularity of IDPConformerGenerator’s design.

To showcase the multiple applications of LDRS, we have modeled five protein systems having different combinations of N-IDR, L-IDR, and C-IDR cases. Supplemental Figure 1 shows the schematics for the folded and IDR elements for studied systems. Side chain atoms can be modeled within IDPConformerGenerator using Monte Carlo-Side Chain Entropy (MC-SCE) (Bhowmick and Head-Gordon 2015), and we made enhancements to MC-SCE that now enable modeling post-translational modifications (PTMs) such as phosphorylation, methylation, N6-carboxylysine, and hydroxylation (Kia-Ki and Martinage 1992; Sirota *et al*. 2015). We also demonstrate the capabilities to model IDRs in transmembrane proteins in the context of a phospholipid bilayer and within multi-chain dynamic complexes. We use structures from the RCSB PDB (Berman *et al*. 2000) having missing coordinates due to lack of data and from the AlphaFold structure prediction database (Jumper *et al*. 2021; Varadi *et al*. 2022), from which we removed the low confidence residues. We modeled missing residues as ensembles using LDRS as demonstrated below.

Figure 1A presents a combination of the N-IDR and C-IDR cases occurring in the 5-fold phosphorylated eukaryotic translation initiation factor 4E-binding protein 2 (5p 4E-BP2), based on the NMR structure (PDB ID 2MX4) (Bah *et al*. 2015). 4E-BP2 is largely disordered but has a conditionally folded ∼40-residue domain upon phosphorylation (Bah *et al*. 2015; Dawson *et al*. 2020), leading to an N-IDR length of 18 residues and a C-IDR length of 59 residues (including 3 phosphorylation sites) surrounding the folded domain (containing 2 phosphorylation sites). The all-atom coordinates generated for 5p 4E-BP2 include the five phosphate groups. For the single L-IDR case (Figure 1B), we have modeled the STAS (sulfate transporter anti-sigma) domain of SLC26A9 (solute carrier family 26 member 9) including 86 residues not found in the electron density of the X-ray structure (PDB ID 7CH1) (Chi *et al*. 2020).

**Figure 1.**
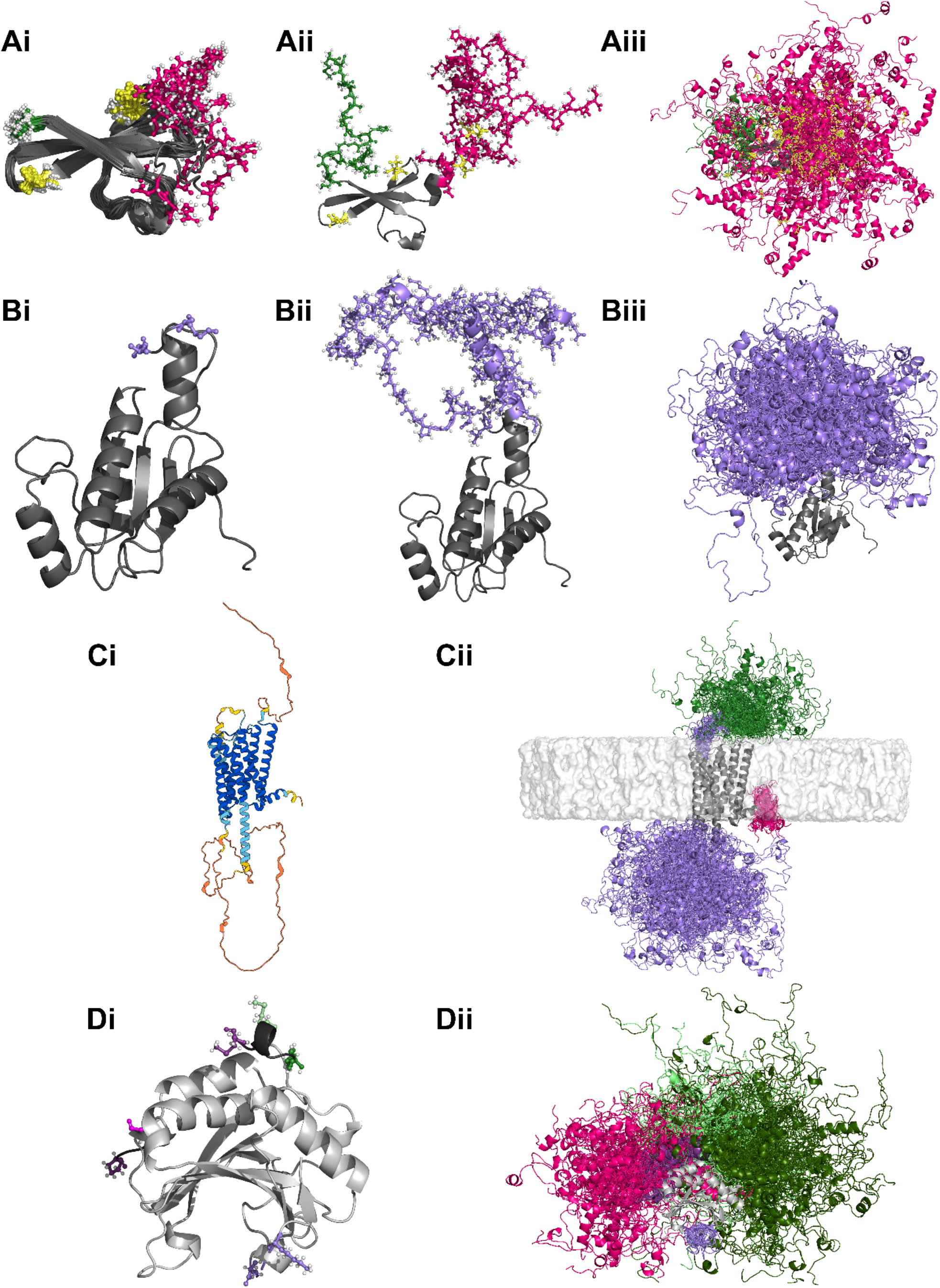
Diverse full-length all-atom and backbone-only ensembles generated using LDRS. Pre-existing solved or predicted structure is shown in grey, N-IDRs in green, L-IDRs in purple, and C-IDRs in magenta. Hydrogen atoms are white in the ball-and-stick models. **(A)** Five-fold phosphorylated 4E-BP2 (5p 4E-BP2): **(Ai)** NMR structure (20 poses from PDB ID 2MX4) with ball-and-stick representations of the terminal residues, N-terminal proline in green and C-terminal arginine in magenta. Phosphorylated residues within the folded domain (Thr35, Thr46) are shown in yellow. **(Aii)** A single full-length all-atom conformer with ball-and-stick representations of sidechains of the IDRs including phosphate residues (Ser65, Thr70, Ser83, in yellow) modeled using LDRS together with MC-SCE. **(Aiii)** Ensemble of 100 (of 1828 calculated) all-atom conformers of 5p 4E-BP2 calculated with LDRS. **(B)** SLC26A9 STAS domain: **(Bi)** X-ray structure (PDB ID 7CH1) with ball-and-stick representations of the residues immediately surrounding the missing loop colored for placement of the missing L-IDR. **(Bii)** A single all-atom model of the full-length domain with ball-and-stick representation of the LDRS-generated L-IDR missing in panel Bi. **(Biii)** Ensemble of 100 (of 692 calculated) all-atom conformers. **(C)** α_2A_ adrenergic receptor: **(Ci)** Structure predicted by AlphaFold (entry P08913), colored according to AlphaFold confidence, with lower confidence regions (pLDDT < 70) in orange and yellow (Jumper *et al*. 2021; Varadi *et al*. 2022). **(Cii)** Ensemble of 100 backbone structures (of 2000 calculated) with AlphaFold lower confidence regions (pLDDT < 70) modeled by LDRS in the context of a bilayer (in light grey). **(D)** 4E-BP2:eIF4E complex: **(Di)** Homology model based on the 4E-BP1:eIF4E X-ray structure (PDB ID 4UED) with 4E-BP2 elements to be modeled as fixed in dark grey and eIF4E in light grey (see Supplemental Information for more details). Ball-and-stick representations of the terminal residues of IDRs to be modeled for eIF4E in light colors and for 4E-BP2 in dark. **(Dii)** Ensemble of 100 (of 2000 calculated) full-length backbone conformers of the 4E-BP2:eIF4E complex with IDRs modeled by LDRS. eIF4E N-IDRs shown in light green and L-IDRs in light purple. 4E-BP2 N-IDRs shown in dark green, L-IDRs in dark purple, and C-IDRs in magenta.

Figure 1C presents the transmembrane α_2A_ adrenergic receptor predicted by AlphaFold (entry P08913), which has extensive regions with low confidence residues (pLDDT < 70) (Jumper *et al*. 2021; Varadi *et al*. 2022). We removed coordinates for these low confidence residues using a simple script that is easily modified to apply to other predicted structures (see Supplemental Information), leading to a 40-residue N-IDR, two L-IDRs of 18 residues and 125 residues, and a 9-residue C-IDR to be modeled by LDRS. Initially, the lipid bilayer was added using OPM (Orientations of Proteins in Membranes) (Lomize *et al*. 2012) and the CHARMM-GUI (Jo *et al*. 2008). The atomic coordinates of the lipid bilayer are then accounted for in LDRS for steric clash checking purposes. While only backbones were modeled for the IDRs, all atomic coordinates of the lipids are present in each of the LDRS models of the α_2A_adrenergic receptor.

Figure 1D shows an ensemble model of the dynamic interaction between the eukaryotic translation initiation factor 4E (eIF4E) and non-phosphorylated 4E-BP2 (Lukhele *et al*. 2013), with 4E-BP2 represented as having two fixed binding sites to eIF4E highlighted in dark grey. The backbone atoms for the N-IDR (54 residues), L-IDR (20 residues), and C-IDR (40 residues) of non-phosphorylated 4E-BP2 and for the N-IDR (35 residues) and L-IDR (9 residues) of eIF4E have been modeled using LDRS. The template for the complex is derived from an X-ray crystal structure of 4E-BP1 bound to eIF4E (PDB ID 4UED) (Peter *et al*. 2015), removing residues of 4E-BP2 that are extremely broadened in NMR spectra of the complex (Lukhele *et al*. 2013). Lastly, we have modeled the cellular tumor antigen p53 (p53) with all-atom conformations using the full-length predicted structure from AlphaFold (entry P04637) but removing low confidence regions (Supplemental Figure 2). Note that the α_2A_ adrenergic receptor transmembrane protein and the 4E-BP2:eIF4E complex are proof-of-principle examples as we only calculated backbone and not all-atom conformers, as additional work is required to automate sidechain generation with coordinate files having lipids and multiple chains.

The fraction secondary structure and Ramachandran plots for the calculated models of these proteins showcase the extent of sampling (Supplemental Figures 3-5). A strength of LDRS is its ability to sample transient helical secondary structure at the boundary of fragment attachment, based on the sequence-based torsion angle builder of IDPConformerGenerator. For example, we observe fractional α-helix secondary structures extending from the existing chains of 5p 4E-BP2 and the SLC26A9 STAS domain (Figures 1Aii and 1Bii). This extension of helical secondary structure is captured by DSSP (Kabsch and Sander 1983) as seen in Supplemental Figure 3. Relative speeds of conformer generation for each system demonstrate the efficiency of the modeling, but are dependent on IDR length and the restrictions from the folded domain and lipid bilayer (Supplemental Table 1).

For ensemble models of IDRs of AlphaFold-predicted structures (Jumper *et al*. 2021; Varadi *et al*. 2022), we find that the conformer ensembles exhibit a range of secondary structures (Supplemental Figures 3.1 and 3.2), as expected based on IDPConformerGenerator’s sampling of PDB-derived torsion angles, compared to the primarily coil or loop structures of the initial AlphaFold model (Supplemental Figure 3.3). In addition, Cα-Cα distances between modeled IDRs and the folded template sample a much broader range than the initial AlphaFold model, including in some cases instances with closer contacts reflective of potential interactions sampled by IDPConformerGenerator (Supplemental Figures 6.1 and 6.2).

A graphical summary of the LDRS method is shown in Supplemental Figures 7.1 and 7.2. The core of the LDRS approach is based on the Kabsch algorithm (Kabsch 1976), a mathematical protocol that ensures the alignment and connectivity of protein backbone fragments. First, LDRS identifies missing residues by comparing the given input PDB structure and full-length sequence. LDRS then builds the missing residues as isolated single chains using the IDPConformerGenerator (Teixeira *et al*. 2022) builder. Single chains are oriented to the sites of N-IDR, L-IDR, and C-IDR cases using the Kabsch algorithm, and a flexible van der Waals radii (Tsai *et al*. 1999) clash check is employed to remove chains that clash with the input structure. Although sidechains can be built during the initial modeling step, we recommend modeling the backbones first before the downstream sidechain and post-translational modification modeling with MC-SCE (Bhowmick and Head-Gordon 2015) to eliminate sidechain clashes.

To model IDRs between folded elements (L-IDR case), we have developed the *next seeker* algorithm in LDRS (see Supplemental Figure 7.1). After generating ensembles of full-length fragments of these IDRs representing missing residues from both sides of the gap, *next seeker* identifies fragment pairs that can close the chain. This is done by verifying that the Cα(i)-C(i)-N(i+1) bond angle, ω backbone torsion angle, and bond lengths (d_C-N_, dC_-Cα_) of residues i and i+1 from each of the respective pairs comply with average values observed in the IDPConformerGenerator database. This database was generated from non-redundant PDB IDs of X-ray crystal structures with resolutions better than or equal to 1.8 Å. After a match has been found, *next seeker* can remodel the carbonyl oxygen and the amino hydrogen at the point of closure.

As LDRS appends a new library of functions in the IDPConformerGenerator API, this generalized tool can be used to model intricate protein systems limited only by the imagination of the user. For example, using the scripts provided in the supplemental material archive as a starting point, LDRS could be used to model ensembles of transmembrane protein interactions comprising several different protein chains.

We envision that the LDRS module within IDPConformerGenerator will be a useful tool to generate ensemble representations of highly flexible loops and tails that are poorly represented by data or other prediction methods. The agreement of *ab initio* IDPConformerGenerator ensembles with experimental data (Teixeira *et al*. 2022) suggests that the IDRs built with LDRS should be representative. If experimental data on torsion angle preferences (i.e., NMR chemical shifts and J-coupling data) are available for the IDRs, the LDRS module in conjunction with the CSSS (custom secondary structure sampling) module (Teixeira *et al*. 2022) within IDPConformerGenerator can build IDRs with these preferences. Furthermore, all-atom conformer ensembles generated with LDRS can be input into sub-setting or reweighting protocols using experimental data, including nuclear magnetic resonance (NMR) spectroscopy, small-angle X-ray scattering (SAXS), and fluorescence resonance energy transfer (FRET) (Bottaro, Bengtsen and Lindorff-Larsen 2020; Gomes *et al*. 2020; Lincoff *et al*. 2020; Liu *et al*. 2023; Tsangaris *et al*. 2023) to generate more realistic ensemble representations that agree with these data. Future development of the LDRS toolkit within the IDPConformerGenerator platform will include streamlining generation of dynamic complexes involving IDPs and IDRs, improving the efficiency of sidechain packing by MC-SCE (Bhowmick and Head-Gordon 2015) and enhancing its capabilities to represent additional post-translational modification types. The expanding toolkit of IDPConformerGenerator will facilitate structural modeling of the many IDRs present in human and other proteomes, providing valuable insights into structure-dynamics-disorder-function relationships.

## Supporting information

supplemental_archive

## ACKNOWLEDGEMENTS

This work was supported by the National Institutes of Health [5R01GM127627-04 and 2R01GM127627-05] to T.H.-G. and J.D.F.-K.; the Natural Sciences and Engineering Research Council of Canada (NSERC) [2016-06718] to J.D.F.-K.; the Canada Research Chairs Program to J.D.F.-K.; and NSERC [RGPIN-2023-04864] to C.C.G. We acknowledge Philip Rößler for helping identify the α_2A_ adrenergic receptor system as a transmembrane protein case, and Zhi Wei Zeng for assistance with the protocol to model the bilayer seen in the α_2A_ adrenergic receptor system.

