## supplemental_information for "Local Disordered Region Sampling (LDRS) for Ensemble Modeling of Proteins with Experimentally Undetermined or Low Confidence Prediction Segments"

Summary. This section presents further details regarding the systems used as examples to demonstrate the capabilities of the Local Disordered Region Sampling (LDRS) module of IDPConformerGenerator (Supplemental Figure 1), models of p53 (entry P04637) (Jumper *et al.* 2021; Varadi *et al.* 2022) (Supplemental Figure 2), secondary structural and Ramachandran torsion angle sampling for the proteins modeled (Supplemental Figures 3, 4 and 5), comparison of Cα-Cα distances between IDRs and the closest residue in folded domains for the LDRS ensembles and AlphaFold structures (Supplemental Figures 6.1 and 6.2), methodological details employed in the LDRS module (Supplemental Text and Supplemental Figures 7.1 and 7.2), and speeds of conformer generation (Supplemental Table 1).

### SUPPLEMENTAL TEXT

Technical specifications of hardware. All computational calculations were performed on the Graham cluster hosted by the Digital Research Alliance of Canada. Each utilized node possesses 32 workers (CPUs) on an Intel E5-2683 v4 Broadwell CPU @ 2.1GHz with a maximum amount of requested RAM at 64 GB.

Modeling parameters. Conformer ensembles generated using IDPConformerGenerator used the same database presented in (Teixeira *et al.* 2022). The agnostic secondary structure sampling flag was used (--dany) along with a backbone energy threshold of 100 kJ (--etbb 100). (Note that other secondary structure sampling flags are possible to utilize, including CSSS to incorporate experimental NMR chemical shift and $^3$J coupling information on torsion angle preferences.) Recommended default settings were used otherwise. Models with sidechains were packed after backbone generation with MC-SCE (Bhowmick and Head-Gordon 2015).

Application of LDRS in Python scripting. Sample scripts in the supplemental materials archive have been given for each system outlining the process to use the modular LDRS libraries to generate conformer ensembles using parallelized computing to generate an ensemble in a shorter period. LDRS libraries could be used for all PDB files, not necessarily only ones generated from IDPConformerGenerator, to take advantage of the built-in clash-checking functions, stitching protocols, as well as the *next seeker* algorithm. As shown in Supplemental Figure 6.1, the alignment and *next seeker* algorithms require the modeled IDR conformer sequence to have a 2 residue overlap with the termini of the template PDB structure to ensure the desired alignment, clash-checking, and stitching results.

Removing low confidence regions of predicted structures. The Python script in the supplemental materials archive (remove_lowconfidence_residues.py) can be used to remove low confidence regions of AlphaFold-predicted structures (Jumper *et al.* 2021; Varadi *et al.* 2022) including p53 (entry P04637) and the $\alpha_{2A}$ adrenergic receptor (entry P08913). Using the recommended pLDDT confidence threshold of 70, we have removed all coordinates of residues with a pLDDT score less than 70 from the predicted structures to be modeled with LDRS. To use the script, change the `input_file` and `output_file` variables to paths pointing to the PDB files of the predicted structure as well as the path for the new processed structure. The threshold value can also be changed to facilitate different metrics used in AlphaFold (Jumper *et al.* 2021), ESMFold (Lin *et al.* 2023), and RoseTTAFold (Baek *et al.* 2021).

Methodology for transmembrane proteins. The upper and lower boundaries of the bilayer was determined using the Orientations of Proteins in Membranes (OPM) web-server (Lomize *et al.* 2012) and all-atom lipids were populated using the CHARMM-GUI web-server (Jo *et al.* 2008). Specifically, PPM 3.0 (Lomize, Todd and Pogozheva 2022) on the OPM web-server was used. After removing the low confidence residues from the AlphaFold structure of the $\alpha_{2A}$ adrenergic receptor transmembrane system (Jumper *et al.* 2021; Varadi *et al.* 2022), the processed PDB file was uploaded to PPM 3.0 where we defined the type of membrane as a mammalian plasma membrane. Curvature was turned off and the N-terminus of the chain was defined as extracellular (topology `out` option). Heteroatoms were not included for positioning of the transmembrane protein in the membrane.

The PPM 3.0 processed PDB structure was then used as an input in the Membrane Builder (Lee *et al.* 2019) input generator from CHARMM-GUI. Due to pre-processing using PPM, our structure is already oriented. The specific size of the system along the x- and y-axes was estimated by taking the length of the longest IDR chain (in this case, 123 residues), dividing it in half and multiplying it by an average length of 3.4 Å per residue (Ainavarapu *et al.* 2007), leading to ~200 Å. A hexagonal bilayer system was then selected and only POPC (phosphatidylcholine) lipids were added. We utilized the conformer from Step 4, foregoing the addition of water and ions, because the purpose of the bilayer here is to prevent IDR conformations that clash with the bilayer region. A single POPC lipid was removed near the first linker region (residue 181) and near the C-terminus (residue 456) to increase the number of conformers without steric clashes. The entire process for adding a bilayer should take no longer than a few minutes depending on the size of the membrane to build using the Membrane Builder function of the CHARMM-GUI.

The final structure of the $\alpha_{2A}$ adrenergic receptor transmembrane protein with the membrane bilayer was used in downstream clash-checking protocols to ensure no generated IDRs clash with the phospholipids in the bilayer. Example scripts have been provided in the A2ADR/scripts folder in the supplemental materials archive.

*Methodology for multi-chain complexes.*
In order for LDRS to recognize multiple protein chains in the same PDB file, we used the IDPConformerGenerator API to split the chains internally to obtain the correct backbone coordinates for alignment for each chain. Clash-checking was performed with the whole structure however, to ensure no backbone steric clashes. An example of the modeling process is presented in the BP24E/scripts folder in the supplemental materials archive.

*Packing sidechains onto select regions of a protein structure.* After modeling the backbone conformations of a protein system, MC-SCE (Bhowmick and Head-Gordon 2015) was used to pack steric-clash free sidechains onto user-specified backbone-only regions of a PDB file. New developments have been made in MC-SCE to enable adding post-translational modifications onto specified sidechains, depending on the residue name (which can be renamed using the `resre` module in IDPConformerGenerator). MC-SCE uses the rotamer library developed by SIDEpro (Nagata, Randall and Baldi 2012) for completing the residues with post-translational modifications dependent on the $\chi_1$ torsion angles that have been selected by the backbone dependent rotamer library (Shapovalov and Dunbrack 2011). Based on the probabilities of occurrence determined in the SIDEpro library, the remaining sidechain torsion angles that define the post-translational modifications are expanded around the standard deviations in a procedure similar to the original MC-SCE algorithm; this will be reported in full in a separate publication. In the 5p 4E-BP2 case, where there are already sidechains from the NMR structures between residues 18-62, MC-SCE accepts a user-specified parameter with the

`--fix` flag to specify the residues with sidechains to ignore (e.g. -f 18-62). For instances where an experimentally determined structure lacks certain sidechains (e.g., at residue 40 in an example protein), MC-SCE can attempt to find sidechain solutions for that residue along with other backbone only residues in the protein system.

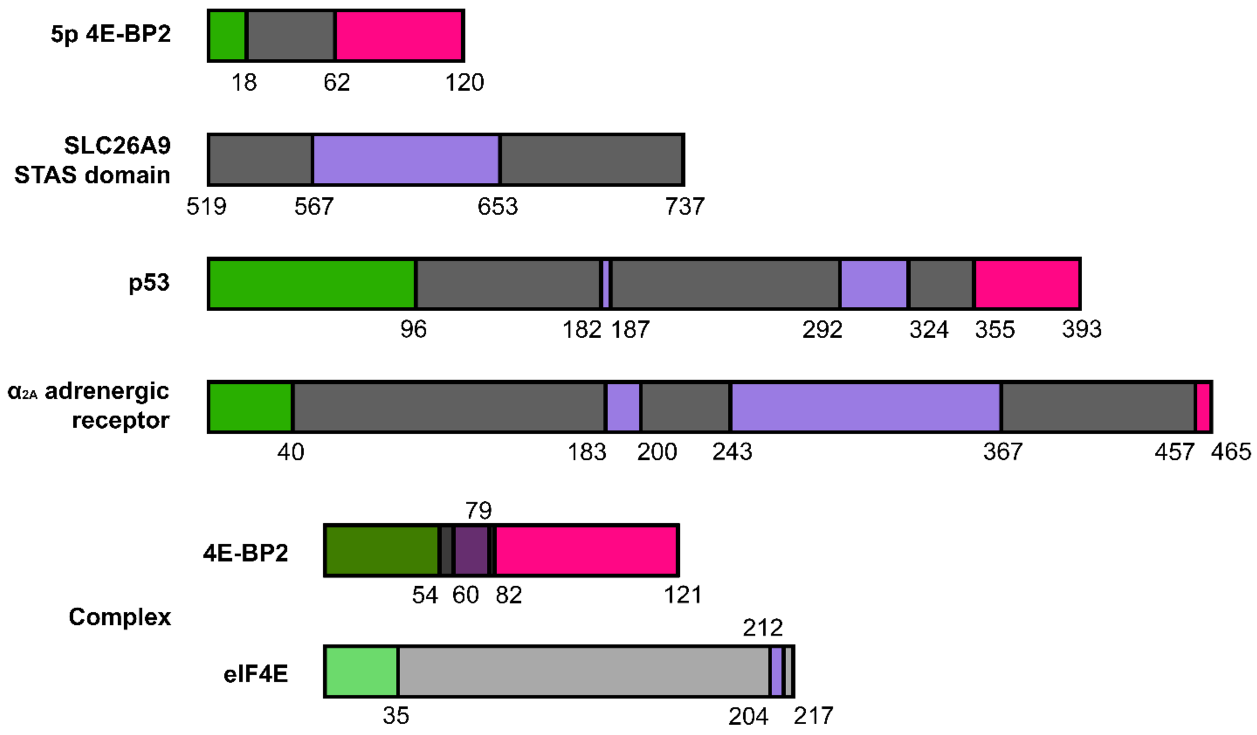


**Supplemental Figure 1. Schematic of folded domains and IDRs modeled by LDRS in each protein system.** Experimentally determined or confidently predicted folded domains are shown in grey rectangles, N-IDRs in green, L-IDRs in purple, and C-IDRs in magenta. Bordering residues of IDRs are included in the IDR to be modeled. The largest L-IDR modeled is 125 residues in the transmembrane $\alpha_{2A}$ adrenergic receptor.

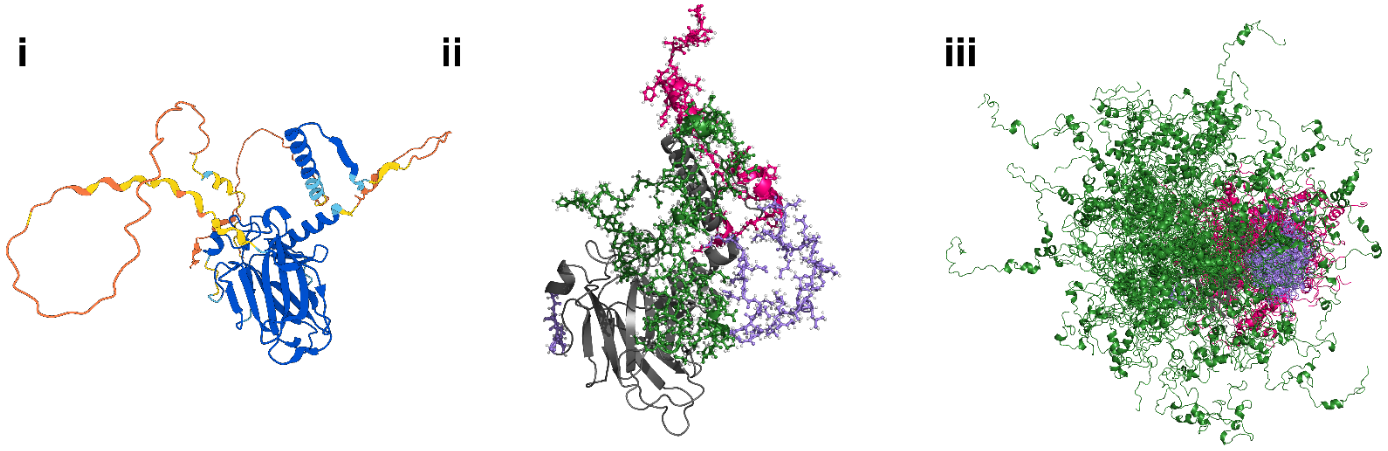


**Supplemental Figure 2. Models of all-atom p53 conformers generated using the LDRS method in IDPConformerGenerator. i)** AlphaFold2 prediction of p53 (entry P04637) (Jumper *et al.* 2021; Varadi *et al.* 2022) highlighting low confidence regions (yellow and orange). **ii)** A single all-atom model of IDRs within p53 generated using IDPConformerGenerator LDRS and MC-SCE to replace low confidence regions, represented in ball-and-stick. Green represents N-IDR, light purple represents L-IDRs, magenta represents C-IDRs, and hydrogens are colored white. **iii)** Ensemble of 100 all-atom conformers of p53 generated with the same coloring as in ii).

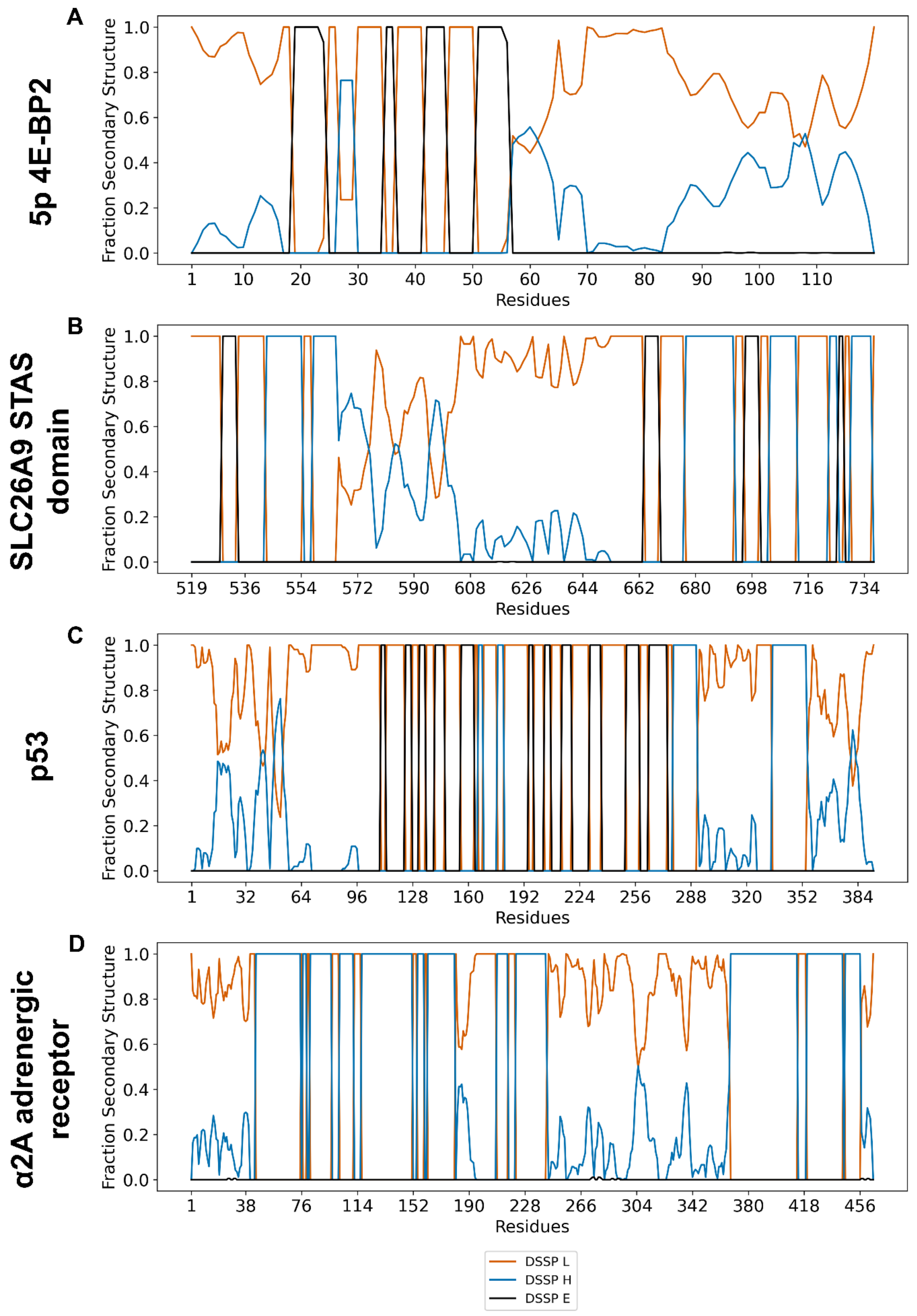
A
5p 4E-BP2
B
SLC26A9 STAS domain
C
p53
D
α2A adrenergic receptor
Fraction Secondary Structure
Residues
DSSP L
DSSP H
DSSP E

**Supplemental Figure 3.1. Fractional secondary structure plots for each single chain example system.** Fractional secondary structure calculated using DSSP (Kabsch and Sander 1983) through IDPConformerGenerator (Teixeira *et al.* 2022). **A)** 5p 4E-BP2, with ensemble of 1828 all-atom conformers used for calculations (folded region from PDB ID 2MX4) (Bah *et al.* 2015). **B)** SLC26A9 STAS domain (PDB ID 7CH1) (Chi *et al.* 2020), with ensemble of 692 all-atom conformers used for calculations. **C)** p53 (AlphaFold entry P04637), with ensemble of 101 all-atom conformers used for calculations. **D)** $\alpha_{2A}$ adrenergic receptor (AlphaFold entry P08913), with ensemble of 2000 backbone conformers used for calculations.

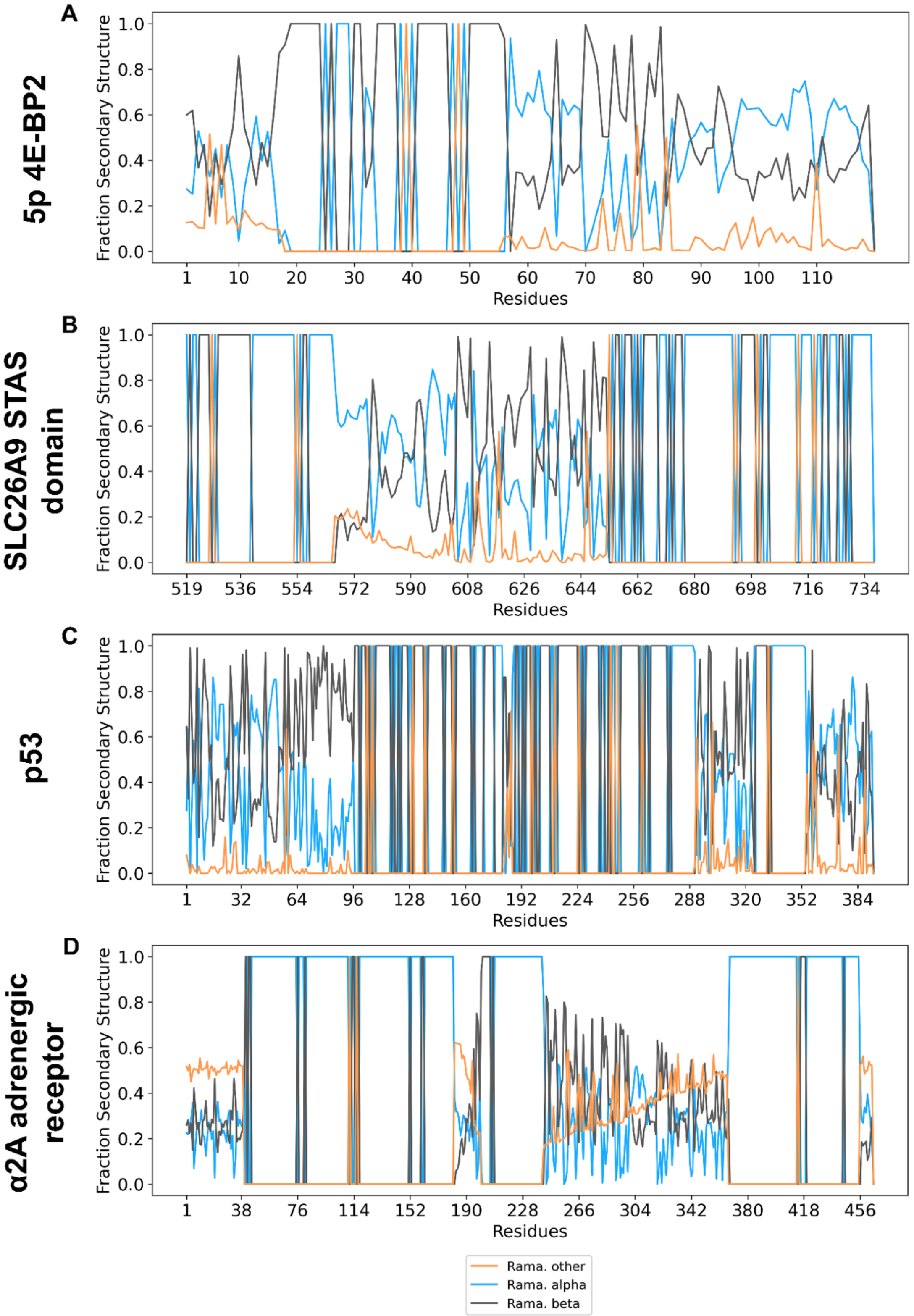


**Supplemental Figure 3.2. Fractional secondary structure plots for each single chain example system.** Fractional secondary structure calculated using alpha, beta, and other regions on the Ramachandran diagram. Descriptions for panels A, B, C, D, are the same as in Supplemental Figure 3.1.

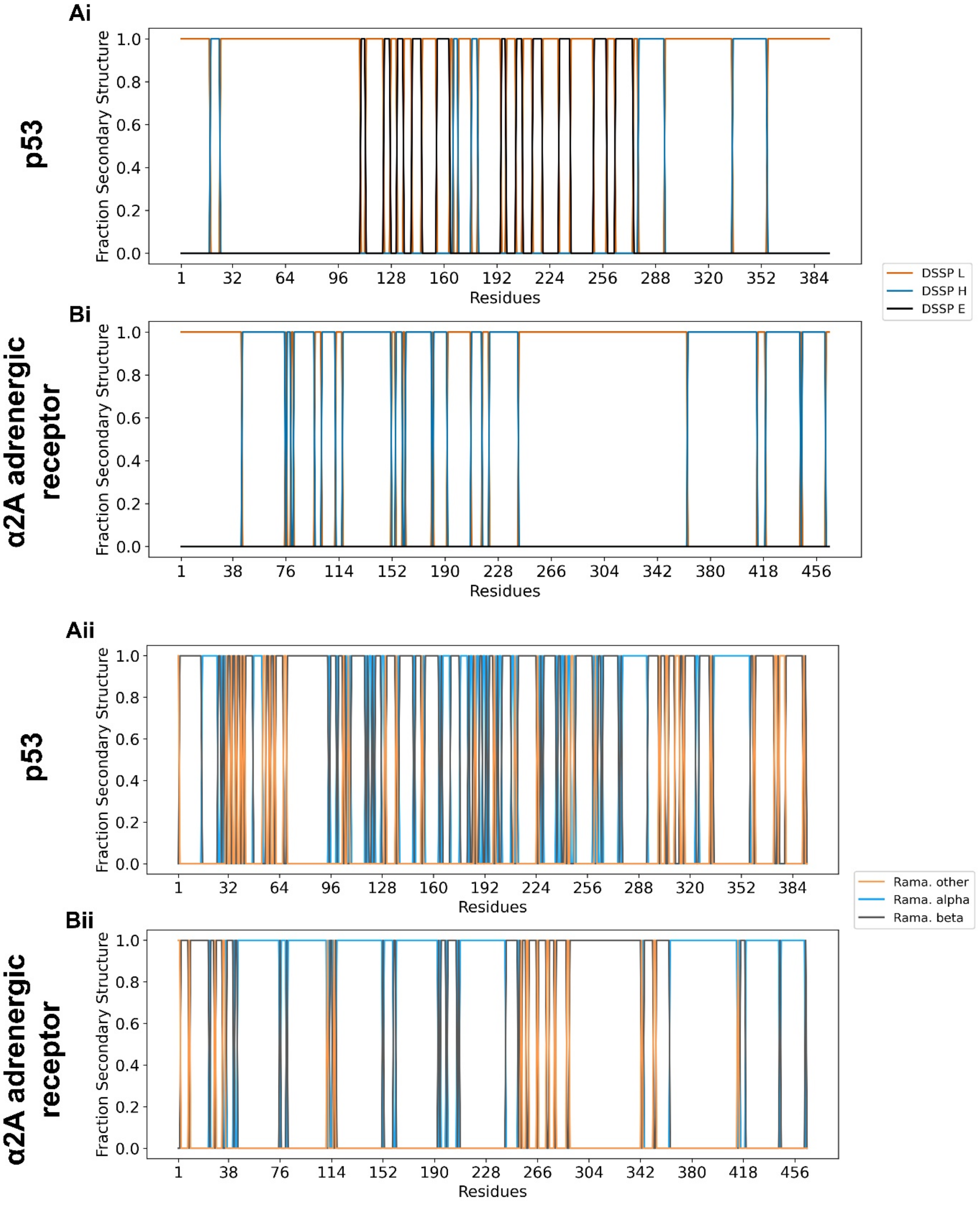


**Supplemental Figure 3.3. Secondary structure plots for AlphaFold predicted proteins. i)** Secondary structure calculated using DSSP (Kabsch and Sander 1983) through IDPConformerGenerator (Teixeira *et al.* 2022). **ii)** Secondary structure calculated using alpha, beta, and other regions on the Ramachandran diagram. **A)** p53 (AlphFold entry P04637). **B)** $\alpha_{2A}$ adrenergic receptor (AlphaFold entry P08913).

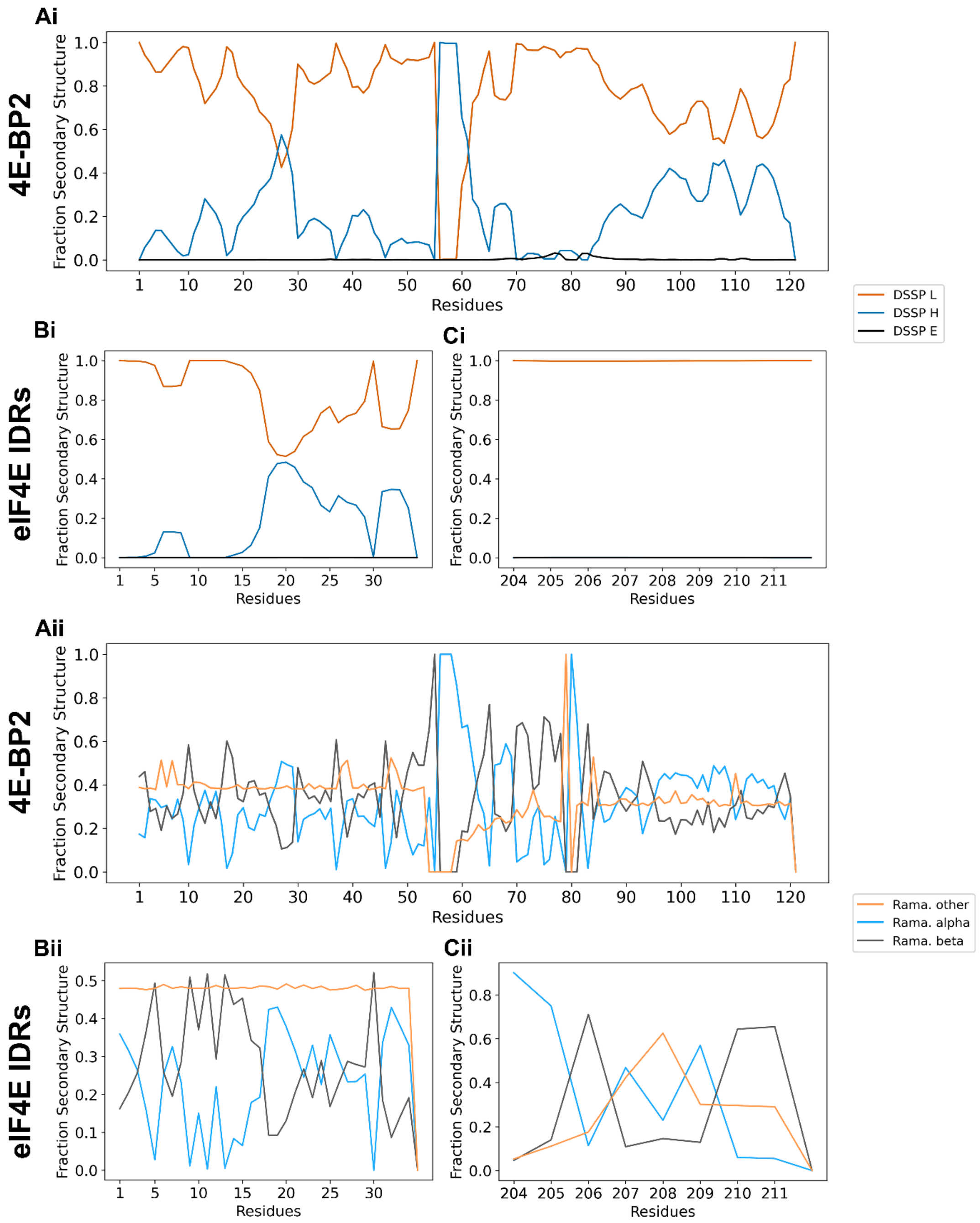


**Supplemental Figure 4. Fractional secondary structure plots for the 4E-BP2:eIF4E complex.** Section **i)** Fractional secondary structure calculated using DSSP (Kabsch and Sander 1983) through IDPConformerGenerator (Teixeira *et al.* 2022). Section **ii)** Fractional secondary structure calculated using alpha, beta, and other regions on the Ramachandran diagram. Ensemble of 2000 backbone chains were used for all calculations. **A)** Models of full length non-phosphorylated 4E-BP2 (with fixed elements from chain B of PDB ID 4UED) (Peter *et al.* 2015). **B)** Models of the N-terminal IDR of eIF4E from chain A of PDB ID 4UED. **C)** Models of the L-IDR from residues 204-212 of eIF4E.

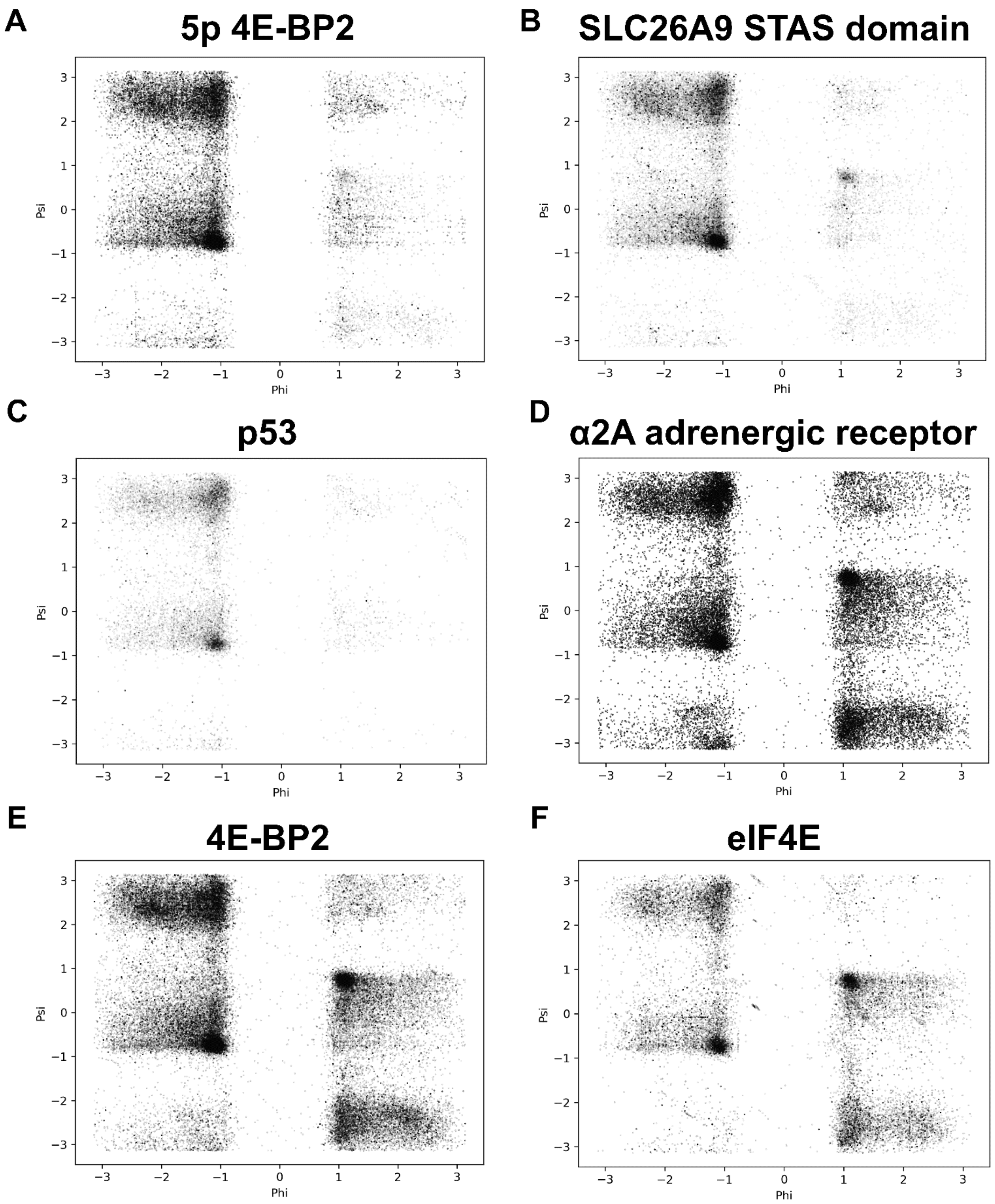
A
5p 4E-BP2
B
SLC26A9 STAS domain
C
p53
D
α2A adrenergic receptor
E
4E-BP2
F
eIF4E
Psi
Phi

**Supplemental Figure 5. Ramachandran diagrams for all modeled IDRs in the studied protein systems.** Only IDRs modeled from each system are presented, demonstrating the sampling using the IDPConformerGenerator database. Units for Phi and Psi are given in radians. **A)** 5p 4E-BP2, with ensemble of 1828 all-atom conformers used for calculations (LDRS-modeled "missing" residues 1-18 and 62-120 from PDB ID 2MX4) (Bah *et al.* 2015). **B)** SLC26A9 STAS domain (LDRS-modeled "missing residues 567-653 from PDB ID 7CH1 (Chi *et al.* 2020), with ensemble of 692 all-atom conformers used for calculations. **C)** p53 (AlphaFold entry P04637), with modeled N-IDR (1-96), L-IDRs (182-187 and 292-324), and C-IDR (355-393) from an ensemble of 101 all-atom conformers. **D)** LDRS-derived models of N-IDR (1-40), L-IDRs (183-200 and 243-367), and C-IDR (457-465) on the $\alpha_{2A}$ adrenergic receptor (AlphaFold entry P08913). **E)** LDRS-derived models of the N-IDR (1-54), L-IDR (60-79), and C-IDR (82-121) of non-phosphorylated 4E-BP2 from chain B of PDB ID 4UED (Peter *et al.* 2015). **F)** LDRS-derived models of the N-IDR (1-35) and L-IDR (204-212) of eIF4E from chain A of PDB ID 4UED. Ensemble of 2000 backbone conformers used for calculations for D, E, F.

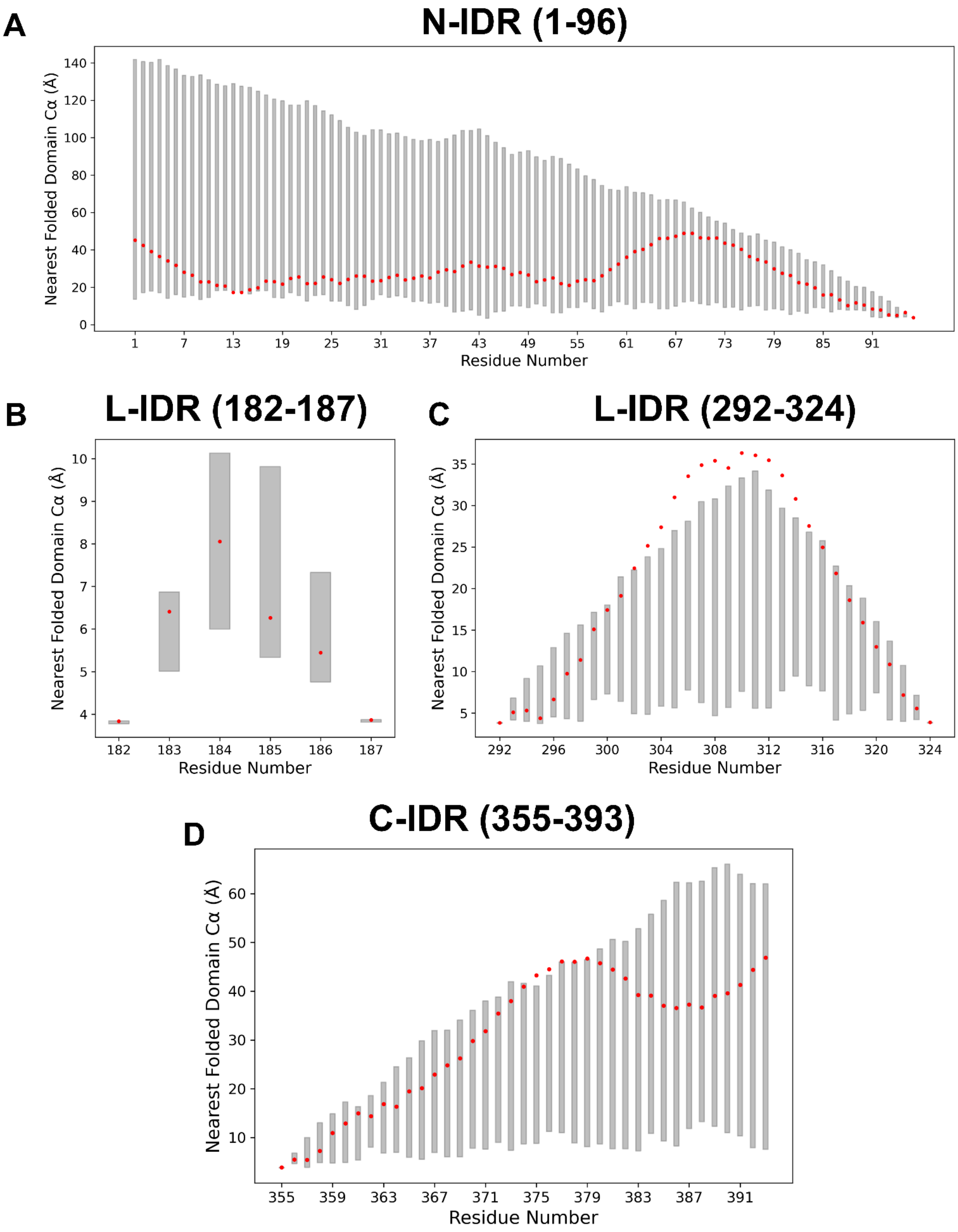


**Supplemental Figure 6.1. Cα-Cα distances between IDRs and the closest residue in folded domains for p53.** AlphaFold (entry P04637) (Jumper *et al.* 2021; Varadi *et al.* 2022) predicted residues are plotted in red. Maximum and minimum ranges from IDRs modeled using LDRS (N = 101) are plotted as grey boxes. **A)** N-terminal IDR (N-IDR), residues 1-96. **B)** First L-IDR, residues 182-187. **C)** Second L-IDR, residues 292-324. **D)** C-terminal IDR (C-IDR), residues 355-393.

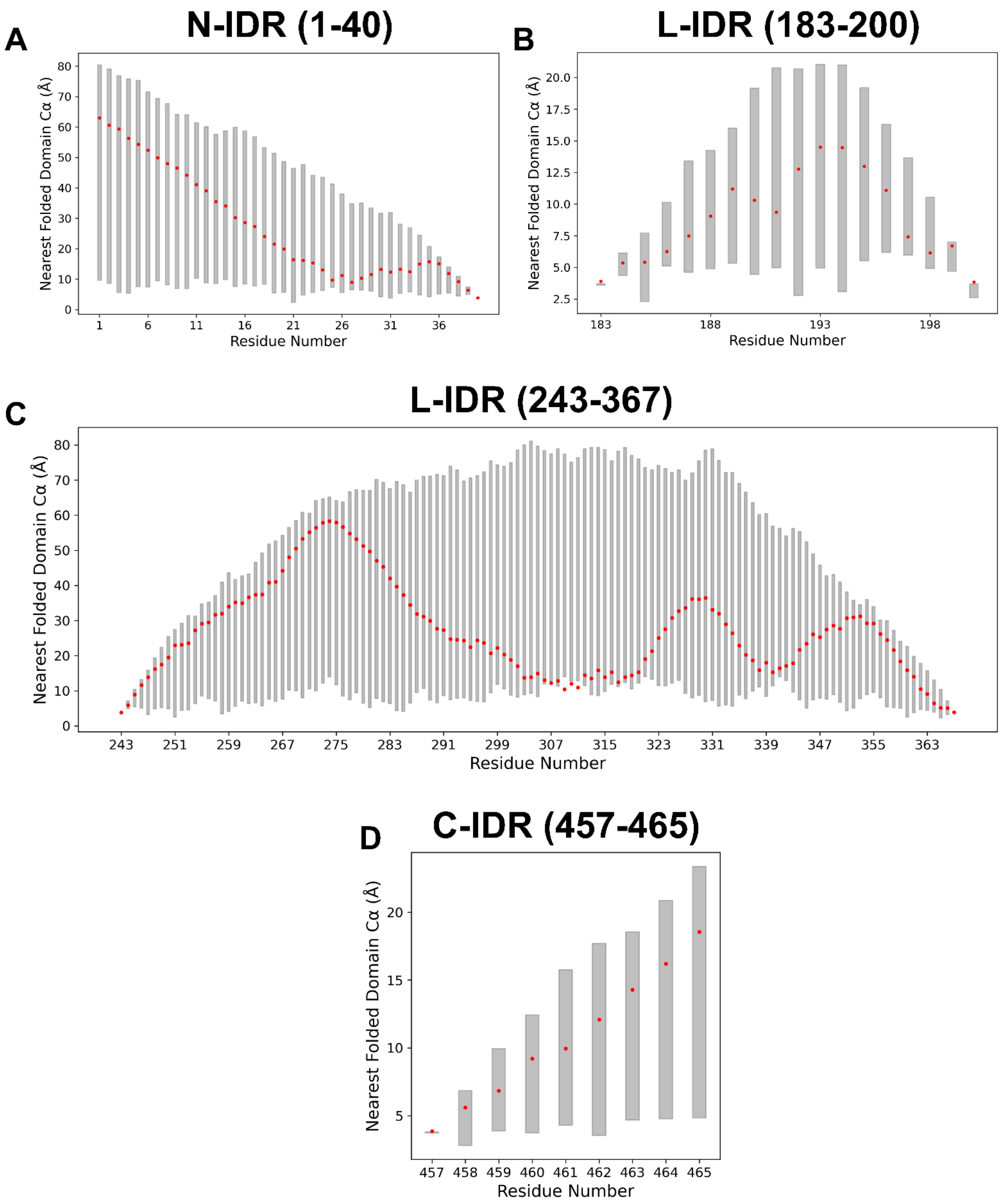


**Supplemental Figure 6.2. Cα-Cα distances between IDRs and the closest residue in folded domains for the $\alpha_{2A}$ adrenergic receptor.** AlphaFold (entry P08913) (Jumper *et al.* 2021; Varadi *et al.* 2022) predicted residues are plotted in red. Maximum and minimum ranges from IDRs modeled using LDRS (N = 2000) are plotted as grey boxes. **A)** N-terminal IDR (N-IDR), residues 1-40. **B)** First L-IDR, residues 183-200. **C)** Second L-IDR, residues 243-367. **D)** C-terminal IDR (C-IDR), residues 457-465.

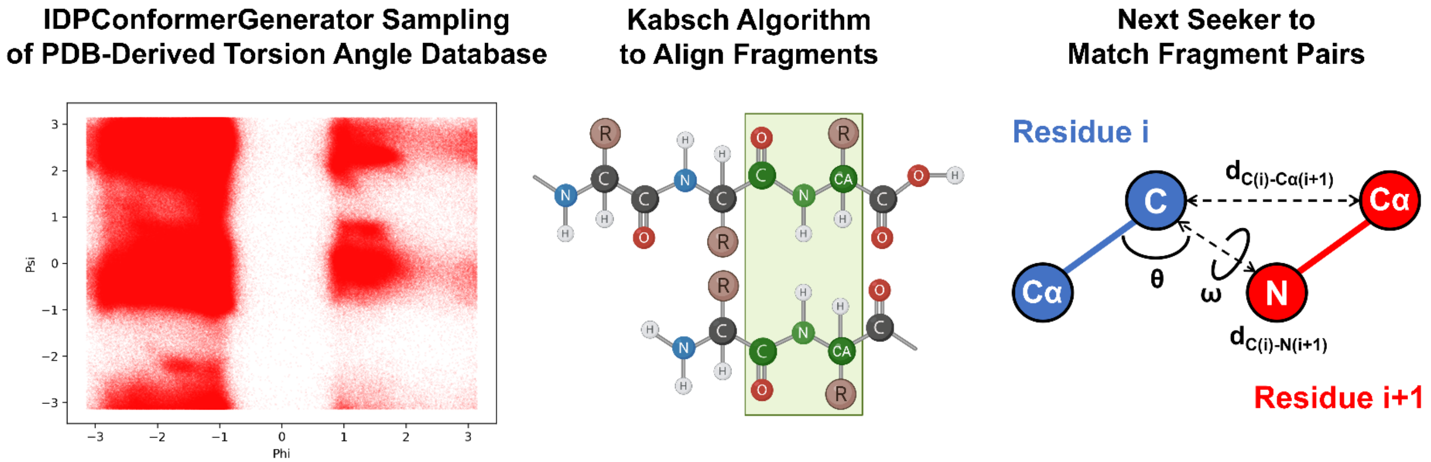


**Supplemental Figure 7.1. Overview of sampling, alignment, and chain break closure methods in LDRS.** IDR fragments are built using IDPConformerGenerator and are aligned to termini fragments using the Kabsch algorithm (Kabsch 1976) and the geometry of Cα(i), C(i), N(i+1), and Cα(i+1) coordinates of overlapping residue regions to maintain backbone continuity. The novel *next seeker* algorithm identifies fragments that can close a broken chain by verifying that the Cα(i)-C(i)-N(i+1) bond angle (θ), ω torsion angle, and bond lengths ($d_{C(i)-N(i+1)}$, $d_{C(i)-C\alpha(i+1)}$) comply with average values observed in the IDPConformerGenerator database of non-redundant PDB IDs of X-ray crystal structures with resolutions better or equal to 1.8 Å (Teixeira *et al.* 2022).

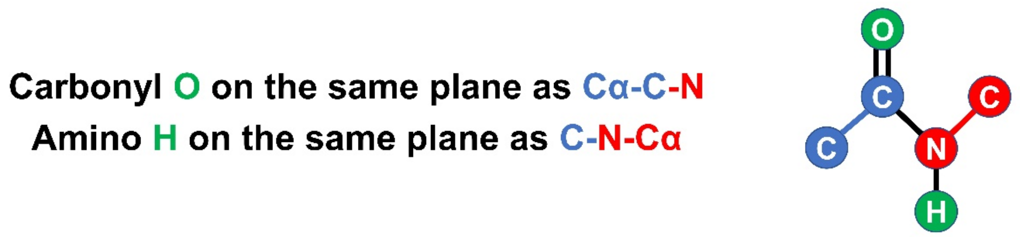


$$\theta_{OXY} = \cos^{-1}\left(\frac{\overrightarrow{C_\alpha C} \cdot \overrightarrow{CN}}{\widehat{C_\alpha C} \cdot \widehat{CN}}\right) \qquad \theta_H = \cos^{-1}\left(\frac{\overrightarrow{CN} \cdot \overrightarrow{NC_\alpha}}{\widehat{CN} \cdot \widehat{NC_\alpha}}\right)$$

$$\overrightarrow{CO} = \vec{C} - |CO|\sin\left(\frac{\theta_{OXY}}{2}\right)\left(\frac{\overrightarrow{C_\alpha C}}{\widehat{C_\alpha C}} + \frac{\overrightarrow{CN}}{\widehat{CN}}\right) \qquad \overrightarrow{NH} = \vec{N} - |NH|\sin\left(\frac{\theta_H}{2}\right)\left(\frac{\overrightarrow{CN}}{\widehat{CN}} + \frac{\overrightarrow{NC_\alpha}}{\widehat{NC_\alpha}}\right)$$

**Supplemental Figure 7.2.** Vector equations used to remodel the carbonyl oxygen and amino hydrogen atoms (green). If residue i+1 is a proline, the amino hydrogen does not need to be remodeled. Blue color corresponds to residue i and red to i+1 in the same manner as Supplemental Figure 6.1.

| Protein System | IDR modeled (number of residues) | Number of backbones calculated | Time for backbones (hr) | Time for alignment (hr) | Number of successful backbones | Time for sidechains (hr) | Number of structures | Time per conformer (min) |
|---|---|---|---|---|---|---|---|---|
| 5p 4E-BP2 | N-IDR (18)<br>C-IDR (59) | 8,320<br>8,320 | 0.036<br>0.067 | 0.106<br>0.229 | 2741<br>1352 | 2.612 | 1,832 | 0.100 |
| SLC26A9 STAS domain | L-IDR (87) | 100,000 | 0.796 | 10.658 | 8,643 | 9.000 | 692 | 1.773 |
| p53 | N-IDR (96)<br>L-IDR (6)<br>L-IDR (33)<br>C-IDR (39) | 100,000<br>100,000<br>100,000<br>100,000 | 1.138<br>0.098<br>0.202<br>0.304 | 2.631<br>1.605<br>6.000<br>0.389 | 56,324<br>58,833<br>3,306<br>62,828 | 5.578 | 101 | 10.660 |
| $\alpha_{2A}$ adrenergic receptor | N-IDR (40)<br>L-IDR (18)<br>L-IDR (125)<br>C-IDR (9) | 100,000<br>100,000<br>100,000<br>100,000 | 0.337<br>0.401<br>2.787<br>0.122 | 0.934<br>0.577<br>22.911<br>0.174 | 7,428<br>188<br>248<br>10,638 | - | 2,000 | 0.847 |
| 4E-BP2: eIF4E complex (BP2:4E) | BP2 N-IDR (54)<br>BP2 L-IDR (20)<br>BP2 C-IDR (40)<br>4E N-IDR (35)<br>4E L-IDR (9) | 60,000<br>100,000<br>60,000<br>60,000<br>100,000 | 0.643<br>0.548<br>0.241<br>0.147<br>0.195 | 2.101<br>5.006<br>1.846<br>0.854<br>1.472 | 11,986<br>811<br>23,991<br>42,829<br>37,543 | - | 2,000 | 0.392 |

**Supplemental Table 1. Time to generate conformers at each step for each system.** Backbones were generated using IDPConformerGenerator (Teixeira *et al.* 2022). Alignment and clash checking were performed using the LDRS functions in the IDPConformerGenerator Python library. All-atom sidechains were attached using MC-SCE (Bhowmick and Head-Gordon 2015). The final number of successfully generated structures were used to calculate the total time per conformer. All alignment and sidechain jobs were run in parallel using 4 compute nodes on the Graham cluster, while backbones were generated on a single node. Note that speed not only depends on the sequence length but also on factors such as the location of the missing chain to be modeled and the occupancy of the folded or existing protein domain. These factors act to reduce the occurrence of clashes between backbone and sidechain atoms during modeling. Sidechains were not modeled for the α2A adrenergic receptor and 4E-BP2:eIF4E complex, denoted with a dash (-) in the column for sidechain time.
