## supplemental_archive for "Local Disordered Region Sampling (LDRS) for Ensemble Modeling of Proteins with Experimentally Undetermined or Low Confidence Prediction Segments": A2ADR_bilayer_cc.py

```
'''
Clash check generated conformers of the
missing linker to the entire protein including
the bilayer of A2ADR.
Author: Zi Hao Liu (@menoliu)
Date: July 17, 2023
'''
import re
import glob
import numpy as np

from functools import partial
from idpconfgen.libs.libmulticore import pool_function
from idpconfgen.libs.libstructure import (
    Structure,
    col_name,
    col_resSeq,
    cols_coords,
    col_segid,
    parse_pdb_to_array,
    write_PDB,
    structure_to_pdb,
    )
from idpconfgen.ldrs_helper import (
    align_coords,
    count_clashes,
    )

from idpconfgen import Path


def term_idr_attach(idr, fld_struc, case, output):
    # Input folded structure should already be built
    fld_name = fld_struc.data_array[:, col_name]
    fld_seq = fld_struc.data_array[:, col_resSeq].astype(int)
    fld_segid = fld_struc.data_array[:, col_segid]
    
    # Hard-coded here for the residue numbers around the chain break
    # because of nature of PDB file with bilayer and protein
    first_seq = 200
    last_seq = 183
    protein_chain = "PROA"
    
    fld_term_idx = {}
    if case == "NIDR":
        case = "N-IDR"
        lu = "U"
        for j, atom in enumerate(fld_name):
            if fld_segid[j] == protein_chain:
                curr_seq = fld_seq[j]
                if curr_seq == first_seq + 1 and atom == "N":
                    fld_term_idx["N"] = j
                elif curr_seq == first_seq + 1 and atom == "CA":
                    fld_term_idx["CA"] = j
                elif curr_seq == first_seq and atom == "C":
                    fld_term_idx["C"] = j
                elif curr_seq == first_seq + 2:
                    break
    elif case == "CIDR":
        case = "C-IDR"
        lu = "L"
        for j, _atom in enumerate(fld_name):
            k = len(fld_name) - 1 - j
            if fld_segid[k] == protein_chain:
                curr_seq = fld_seq[k]
                if curr_seq == last_seq and fld_name[k] == "N":
                    fld_term_idx["N"] = k
                elif curr_seq == last_seq and fld_name[k] == "CA":
                    fld_term_idx["CA"] = k
                elif curr_seq == last_seq - 1 and fld_name[k] == "C":
                    fld_term_idx["C"] = k
                elif curr_seq == last_seq - 2:
                    break
    
    fld_Cxyz = fld_struc.data_array[fld_term_idx["C"]][cols_coords].astype(float)  # noqa: E501
    fld_Nxyz = fld_struc.data_array[fld_term_idx["N"]][cols_coords].astype(float)  # noqa: E501
    fld_CAxyz = fld_struc.data_array[fld_term_idx["CA"]][cols_coords].astype(float)  # noqa: E501
    # Coordinates of boundary to stitch to later on
    fld_coords = np.array([fld_Cxyz, fld_Nxyz, fld_CAxyz])
    
    with open(idr) as f:
        idr_data = f.read()
    try:
        idr_arr = parse_pdb_to_array(idr_data)
    except AssertionError:
        return

    aligned_idr = align_coords(idr_arr, fld_coords, case)
    clashes, fragment = count_clashes(
        aligned_idr,
        fld_struc,
        case,
        max_clash=80,
        tolerance=0.8
        )
    if type(clashes) is int:
        aligned_struc = structure_to_pdb(fragment)
        idr_split = idr.split("/")
        idr_int = int(re.search(r'\d+', idr_split[-1]).group())
        # Change PDB name here if desired
        write_PDB(aligned_struc, output + f"aligned_1{lu}_{idr_int}.pdb")


# Change variables here
parent_folder = "/PATH/TO/PARENT/FOLDER"
ncores = 60

# The first chain break (LIDR1)
# "L" signifies lower, or N-terminal part of the linker
case = "LIDR1_L"
idr_lst = glob.glob(f"/{parent_folder}/{case}/*.pdb")
fld = f"{parent_folder}/A2A_bilayer_hex_clean.pdb"
fld_struc = Structure(Path(fld))
fld_struc.build()
output = f"{parent_folder}/{case}_filtered/"

execute = partial(
    term_idr_attach,
    fld_struc=fld_struc,
    case="CIDR",
    output=output,
    )
execute_pool = pool_function(execute, idr_lst, ncores=ncores)
for _ in execute_pool:
    pass

# "U" signifies upper, or C-terminal part of the linker
case = "LIDR1_U"
idr_lst = glob.glob(f"/{parent_folder}/{case}/*.pdb")
fld = f"{parent_folder}/A2A_bilayer_hex_clean.pdb"
fld_struc = Structure(Path(fld))
fld_struc.build()
output = f"{parent_folder}/{case}_filtered/"

execute = partial(
    term_idr_attach,
    fld_struc=fld_struc,
    case="NIDR",
    output=output,
    )
execute_pool = pool_function(execute, idr_lst, ncores=ncores)
for _ in execute_pool:
    pass
```
