## supplemental_archive for "Local Disordered Region Sampling (LDRS) for Ensemble Modeling of Proteins with Experimentally Undetermined or Low Confidence Prediction Segments": A2ADR_plot_idr_fld_dist.py

```
'''
Plot CA distance distribution of IDRs to folded domain for A2ADR.
Author: Zi Hao Liu (@menoliu)
Date: July 20, 2023
'''
import glob
import numpy as np
import matplotlib.pyplot as plt

from functools import partial
from idpconfgen import Path
from idpconfgen.libs.libmulticore import pool_function
from idpconfgen.libs.libstructure import (
    Structure,
    col_name,
    col_resSeq,
    cols_coords,
    )
from idpconfgen.ldrs_helper import calculate_distance


def get_fld_ca_coords(path, ranges):
    struc = Structure(Path(path))
    struc.build()
    atom_names = struc.data_array[:, col_name]
    res_seqs = struc.data_array[:, col_resSeq].astype(int)
    coords = struc.data_array[:, cols_coords].astype(float)
    
    fld_ca_coords = []
    
    for r in ranges:
        lower = r[0]
        upper = r[1]
        
        for i, res in enumerate(res_seqs):
            if lower <= res <= upper:
                name = atom_names[i]
                if name == "CA":
                    fld_ca_coords.append(coords[i])
            elif res > upper:
                break
    
    return fld_ca_coords


def calc_dist_distributions(path, fld_ca_coords, ranges):
    struc = Structure(Path(path))
    struc.build()
    atom_names = struc.data_array[:, col_name]
    res_seqs = struc.data_array[:, col_resSeq].astype(int)
    coords = struc.data_array[:, cols_coords].astype(float)
    
    total_distributions = []
    
    for r in ranges:
        lower = r[0]
        upper = r[1]
        
        idr_coords = []
        distances = []
        for j, seq in enumerate(res_seqs):
            if lower <= seq <= upper:
                name = atom_names[j]
                if name == "CA":
                    idr_coords.append(coords[j])
            elif seq > upper:
                break
        
        for ic in idr_coords:
            d = []
            for fc in fld_ca_coords:
                d.append(calculate_distance(ic, fc))
            distances.append(min(d))
        
        total_distributions.append(distances)
    
    return total_distributions


def plot_distance_scatter(fld_values, idr_ranges, res, output):
    lower = res[0] - res[0] + 1
    upper = res[1] - res[0] + 1
    # Change width here to fit more points in figure
    width = 5
    fig, ax = plt.subplots()
    total_res = upper - lower + 1
    increment = int(total_res / width)
    
    xvals = np.arange(lower, upper + 1, 1)
    xticks = np.arange(lower, upper + 1, increment)
    xticks_labels = list(xticks + res[0] - 1)
    
    for i, r in enumerate(idr_ranges):
        ax.fill_between([i + 0.8, i + 1.2], r[0], r[1], color='gray', alpha=0.5)
        
    ax.scatter(
        xvals,
        fld_values,
        color='r',
        zorder=2,
        alpha=1.0,
        s=6
        )
    
    ax.set_xticks(xticks)
    ax.set_xticklabels(labels=xticks_labels)
    ax.tick_params(axis='x', labelsize=12)
    ax.tick_params(axis='y', labelsize=12)
    plt.xlabel("Residue Number", fontsize=15)
    plt.ylabel("Nearest Folded Domain Cα (Å)", fontsize=15)
    fig.set_size_inches(width, 5)
    fig.savefig(output, dpi=300)
    

# Change index and path of plot_output for different results across the ranges
alphafold_structure = "./A2ADR_P08913_aligned.pdb"
final_conformers = glob.glob("./A2ADR_conformers/*.pdb")
plot_output = "./figs/A2ADR_CIDR_distances.png"
ncores = 5
index = 3
idr_res = ((1, 40), (183, 200), (243, 367), (457, 465))
fld_res = ((41, 182), (201, 242), (368, 456))

af2_ca_coords = get_fld_ca_coords(alphafold_structure, fld_res)
af2_idr_dists = calc_dist_distributions(alphafold_structure, af2_ca_coords, idr_res)

collection = []

execute = partial(
    calc_dist_distributions,
    fld_ca_coords=af2_ca_coords,
    ranges=idr_res,
    )
execute_pool = pool_function(execute, final_conformers, ncores=ncores)
for dist in execute_pool:
    collection.append(dist[index])

transposed_data = list(zip(*collection))
idr_ranges = [(min(column), max(column)) for column in transposed_data]

plot_distance_scatter(af2_idr_dists[index], idr_ranges, idr_res[index], plot_output)
```
