## supplemental_archive for "Local Disordered Region Sampling (LDRS) for Ensemble Modeling of Proteins with Experimentally Undetermined or Low Confidence Prediction Segments": A2ADR_stitching.py

```
'''
Clash-checking modeled IDRs with each other
and attach them onto the A2ADR transmembrane
protein.
Author: Zi Hao Liu (@menoliu)
Date: July 17, 2023
'''
import os
import hashlib
import glob
import random
import numpy as np

from functools import partial
from multiprocessing import Queue
from idpconfgen.libs.libmulticore import pool_function
from idpconfgen.libs.libstructure import (
    Structure,
    col_chainID,
    col_segid,
    col_serial,
    col_resSeq,
    parse_pdb_to_array,
    structure_to_pdb,
    )
from idpconfgen.ldrs_helper import count_clashes

from idpconfgen import Path


CONF_NUMBER = Queue()


def remove_duplicates(folder_path):
    # Get a list of all files in the folder
    file_list = os.listdir(folder_path)
    
    # Create a dictionary to store file content hashes
    content_hashes = {}
    
    # Iterate over each file in the folder
    for file_name in file_list:
        file_path = os.path.join(folder_path, file_name)
        
        # Calculate the content hash of the file
        with open(file_path, 'rb') as file:
            file_content = file.read()
            content_hash = hashlib.md5(file_content).hexdigest()
        
        # Check if a file with the same content hash already exists
        if content_hash in content_hashes:
            # Delete the current file
            os.remove(file_path)
        else:
            # Store the content hash and file path in the dictionary
            content_hashes[content_hash] = file_path


def check_clash_quattro(double_quattro):
    # `double_quattro` is a set of (quattro_arr, quatrro_struc)
    # order is (nidr, lidr1, lidr2, cidr)
    nidr_arr = double_quattro[0][0]
    lidr1_arr = double_quattro[0][1]
    cidr_arr = double_quattro[0][3]
    lidr1_struc = double_quattro[1][1]
    lidr2_struc = double_quattro[1][2]
    cidr_struc = double_quattro[1][3]
    
    nb1, _ = count_clashes(nidr_arr, lidr1_struc)
    nb2, _ = count_clashes(nidr_arr, lidr2_struc)
    nc, _ = count_clashes(nidr_arr, cidr_struc)
    b1b2, _ = count_clashes(lidr1_arr, lidr2_struc)
    b1c, _ = count_clashes(lidr1_arr, cidr_struc)
    b2c, _ = count_clashes(cidr_arr, lidr2_struc)

    if type(nb1) is bool or \
        type(nb2) is bool or \
        type(nc) is bool or \
        type(b1b2) is bool or \
        type(b1c) is bool or \
        type(b2c) is bool:
            return None
    
    return double_quattro[0]


# Customized psurgeon function. Original can be found in ldrs_helper.py
# Stitching should be done with a "prepared" PDB file with the
# first residues of termini and IDRs removed
def psurgeon_quattro(quattro_arr, fld_arr, output, rs):
    nidr = quattro_arr[0]
    seq_nidr = nidr[:, col_resSeq]
    
    lidr1 = quattro_arr[1]
    seq_lidr1 = lidr1[:, col_resSeq]
    
    lidr2 = quattro_arr[2]
    seq_lidr2 = lidr2[:, col_resSeq]
    
    cidr = quattro_arr[3]
    seq_cidr = cidr[:, col_resSeq]
    
    # Remove last residue on NIDR
    for i, _ in enumerate(seq_nidr):
        j = len(seq_nidr) - 1 - i
        curr = seq_nidr[j]
        prev = seq_nidr[j - 1]
        nidr = nidr[:-1]
        if prev != curr:
            break
    
    # Remove first and last residue on lidr1 and lidr2
    for i, seq in enumerate(seq_lidr1):
            next = seq_lidr1[i + 1]
            lidr1 = lidr1[1:]
            if next != seq:
                break
    for i, _ in enumerate(seq_lidr1):
        j = len(seq_lidr1) - 1 - i
        curr = seq_lidr1[j]
        prev = seq_lidr1[j - 1]
        lidr1 = lidr1[:-1]
        if prev != curr:
            break
    
    for i, seq in enumerate(seq_lidr2):
            next = seq_lidr2[i + 1]
            lidr2 = lidr2[1:]
            if next != seq:
                break
    for i, _ in enumerate(seq_lidr2):
        j = len(seq_lidr2) - 1 - i
        curr = seq_lidr2[j]
        prev = seq_lidr2[j - 1]
        lidr2 = lidr2[:-1]
        if prev != curr:
            break
    
    # Remove first residue on CIDR
    for i, seq in enumerate(seq_cidr):
            next = seq_cidr[i + 1]
            cidr = cidr[1:]
            if next != seq:
                break
    
    nidr_lst = nidr.tolist()
    fld_lst = fld_arr.tolist()
    
    # Correct residue numbering for lidr1 (+181)
    for i, line in enumerate(lidr1):
        resnum = int(line[col_resSeq])
        lidr1[i][col_resSeq] = str(resnum + 181)
    lidr1_lst = lidr1.tolist()
    # Insert lidr1
    fld_seq = fld_arr[:, col_resSeq]
    for i, seq in enumerate(fld_seq):
        next_seq = fld_seq[i + 1]
        if int(next_seq) - int(seq) > 1:
            lidr1_idx = i + 1
            break
    for i, row in enumerate(lidr1_lst):
        fld_lst.insert(lidr1_idx + i, row)
    
    # Correct residue numbering for lidr2 (+241)
    for i, line in enumerate(lidr2):
        resnum = int(line[col_resSeq])
        lidr2[i][col_resSeq] = str(resnum + 241)
    lidr2_lst = lidr2.tolist()
    # Insert lidr2
    fld_arr = np.array(fld_lst)
    fld_seq = fld_arr[:, col_resSeq]
    for i, seq in enumerate(fld_seq):
        next_seq = fld_seq[i + 1]
        if int(next_seq) - int(seq) > 1:
            lidr2_idx = i + 1
            break
    for i, row in enumerate(lidr2_lst):
        fld_lst.insert(lidr2_idx + i, row)
    
    # Correct residue numbering for CIDR (+455)
    for i, line in enumerate(cidr):
        resnum = int(line[col_resSeq])
        cidr[i][col_resSeq] = str(resnum + 455)
    cidr_lst = cidr.tolist()
    
    new_struc_lst = nidr_lst + fld_lst + cidr_lst
    new_struc_arr = np.array(new_struc_lst)
    new_serial = [str(i) for i in range(1, len(new_struc_arr) + 1)]
    new_struc_arr[:, col_serial] = new_serial
    new_struc_arr[:, col_chainID] = "A"
    new_struc_arr[:, col_segid] = "A"
    new_struc = structure_to_pdb(new_struc_arr)

    with open(output + f"conformer_rs{rs}_{CONF_NUMBER.get()}.pdb", 'w') as fout:
        for line in new_struc:
            fout.write(line + "\n")
    
        
# Change variables here
parent_path = "/home/nemoliu/Documents/modelling/A2A/results"
nconfs = 180
ncores = 60

output_path = f"{parent_path}/A2A_bb_confs/"
nidr_paths = glob.glob(f"{parent_path}/NIDR_results/*.pdb")
lidr1_paths = glob.glob(f"{parent_path}/LIDR1_results/*.pdb")
lidr2_paths = glob.glob(f"{parent_path}/LIDR2_results/*.pdb")
cidr_paths = glob.glob(f"{parent_path}/CIDR_results/*.pdb")
fld_path = f"{parent_path}/A2A_prepped.pdb"
fld_struc = Structure(Path(fld_path))
fld_struc.build()
fld_arr = fld_struc.data_array

for rs in range(17):
    random.seed(rs)
    
    for i in range(1, nconfs + 1):
        CONF_NUMBER.put(i)

    nidr_arr = []
    nidr_struc = []
    lidr1_arr = []
    lidr1_struc = []
    lidr2_arr = []
    lidr2_struc = []
    cidr_arr = []
    cidr_struc = []

    quattro_check = []
    final_quattro_arr = []

    random.shuffle(nidr_paths)
    random.shuffle(lidr1_paths)
    random.shuffle(lidr2_paths)
    random.shuffle(cidr_paths)
    nidr_paths = nidr_paths[:nconfs]
    lidr1_paths = lidr1_paths[:nconfs]
    lidr2_paths = lidr2_paths[:nconfs]
    cidr_paths = cidr_paths[:nconfs]

    quattro_paths = []

    for i, nidr in enumerate(nidr_paths):
        quattro_paths.append([nidr, lidr1_paths[i], lidr2_paths[i], cidr_paths[i]])

    for p in quattro_paths:
        p_nidr = p[0]
        p_lidr1 = p[1]
        p_lidr2 = p[2]
        p_cidr = p[3]

        with open(p_nidr) as nidr:
            data_nidr = nidr.read()
        with open(p_lidr1) as lidr1:
            data_lidr1 = lidr1.read()
        with open(p_lidr2) as lidr2:
            data_lidr2 = lidr2.read()
        with open(p_cidr) as cidr:
            data_cidr = cidr.read()

        try:
            arr = parse_pdb_to_array(data_nidr)
            struc = Structure(Path(p_nidr))
            struc.build()

            nidr_arr.append(arr)
            nidr_struc.append(struc)

            arr = parse_pdb_to_array(data_lidr1)
            struc = Structure(Path(p_lidr1))
            struc.build()

            lidr1_arr.append(arr)
            lidr1_struc.append(struc)

            arr = parse_pdb_to_array(data_lidr2)
            struc = Structure(Path(p_lidr2))
            struc.build()

            lidr2_arr.append(arr)
            lidr2_struc.append(struc)

            arr = parse_pdb_to_array(data_cidr)
            struc = Structure(Path(p_cidr))
            struc.build()

            cidr_arr.append(arr)
            cidr_struc.append(struc)
        except AssertionError:
            continue

    quattro_arr = []

    for i, _ in enumerate(quattro_paths):
        trio_arr = ([nidr_arr[i], lidr1_arr[i], lidr2_arr[i], cidr_arr[i]])
        trio_struc = [nidr_struc[i], lidr1_struc[i], lidr2_struc[i], cidr_struc[i]]
        quattro_check.append((trio_arr, trio_struc))


    execute_clash = pool_function(check_clash_quattro, quattro_check, ncores=ncores)
    for result in execute_clash:
        if result is not None:
            final_quattro_arr.append(result)

    execute = partial(
        psurgeon_quattro,
        fld_arr=fld_arr,
        output=output_path,
        rs=rs
        )
    execute_pool = pool_function(execute, final_quattro_arr, ncores=ncores)
    for _ in execute_pool:
        pass

remove_duplicates(output_path)
```
