## supplemental_archive for "Local Disordered Region Sampling (LDRS) for Ensemble Modeling of Proteins with Experimentally Undetermined or Low Confidence Prediction Segments": BP24E_LIDR.py

```
'''
Align and clash-check possible linker IDRs for
both eIF4E and 4E-BP2 in the BP2:4E complex.
Author: Zi Hao Liu (@menoliu)
Date: July 17, 2023
'''
import glob
import re

import numpy as np

from functools import partial
from idpconfgen.libs.libmulticore import pool_function
from idpconfgen.libs.libstructure import (
    Structure,
    col_name,
    col_resSeq,
    col_chainID,
    cols_coords,
    parse_pdb_to_array,
    write_PDB,
    structure_to_pdb,
    )
from idpconfgen.ldrs_helper import (
    align_coords,
    count_clashes,
    )

from idpconfgen import Path


def break_idr_attach(idr, fld, upper, lower, chain, output_upper=False, output_lower=False):
    fld_struc = Structure(Path(fld))
    fld_struc.build()
    atom_names = fld_struc.data_array[:, col_name]
    fld_seq = fld_struc.data_array[:, col_resSeq].astype(int)
    chain_ids = fld_struc.data_array[:, col_chainID]
    first_seq = fld_seq[0]
    if first_seq > 1:
        fld_seq -= first_seq - 1
        first_seq = fld_seq[0]
    
    fld_term_idx = {}
    for s, aa in enumerate(fld_seq):
        c = chain_ids[s]
        if c == chain:
            if aa == lower - 1 and atom_names[s] == 'C':
                fld_term_idx["CL"] = s
            elif aa == lower:
                if atom_names[s] == 'N':
                    fld_term_idx["NL"] = s
                elif atom_names[s] == 'CA':
                    fld_term_idx["CAL"] = s
            elif aa == upper and atom_names[s] == 'C':
                fld_term_idx['CU'] = s
            elif aa == upper + 1:
                if atom_names[s] == 'N':
                    fld_term_idx['NU'] = s
                elif atom_names[s] == 'CA':
                    fld_term_idx['CAU'] = s
            if aa > upper + 1:
                break
    
    fld_CLxyz = fld_struc.data_array[fld_term_idx["CL"]][cols_coords].astype(float)
    fld_NLxyz = fld_struc.data_array[fld_term_idx["NL"]][cols_coords].astype(float)
    fld_CALxyz = fld_struc.data_array[fld_term_idx["CAL"]][cols_coords].astype(float)
    fld_coords_L = np.array([fld_CLxyz, fld_NLxyz, fld_CALxyz])
    
    fld_CUxyz = fld_struc.data_array[fld_term_idx["CU"]][cols_coords].astype(float)
    fld_NUxyz = fld_struc.data_array[fld_term_idx["NU"]][cols_coords].astype(float)
    fld_CAUxyz = fld_struc.data_array[fld_term_idx["CAU"]][cols_coords].astype(float)
    fld_coords_U = np.array([fld_CUxyz, fld_NUxyz, fld_CAUxyz])
    
    with open(idr) as f:
        idr_data = f.read()
    try:
        idr_arr = parse_pdb_to_array(idr_data)
    except AssertionError:
        return
    
    if output_lower:
        aligned_break_L = align_coords(idr_arr, fld_coords_L, "C-IDR")
        clashesL, fragmentL = count_clashes(
            aligned_break_L,
            fld_struc,
            "C-IDR",
            max_clash=40,
            tolerance=0.4
            )
        if type(clashesL) is int:
            aligned_struc = structure_to_pdb(fragmentL)
            idr_split = idr.split("/")
            idr_int = int(re.search(r'\d+', idr_split[-1]).group())
            write_PDB(aligned_struc, output_lower + f"aligned_L_{idr_int}.pdb")
    
    elif output_upper:
        aligned_break_U = align_coords(idr_arr, fld_coords_U, "N-IDR")
        clashesU, fragmentU = count_clashes(
            aligned_break_U,
            fld_struc,
            "N-IDR",
            max_clash=40,
            tolerance=0.4
            )
        if type(clashesU) is int:
            aligned_struc = structure_to_pdb(fragmentU)
            idr_split = idr.split("/")
            idr_int = int(re.search(r'\d+', idr_split[-1]).group())
            write_PDB(aligned_struc, output_upper + f"aligned_U_{idr_int}.pdb")


# Change variables here
parent = "/PATH/TO/PARENT/FOLDER"
pdb_fld = f"{parent}/4UED_cleaned_both.pdb"
ncores = 32

# For the L-IDR on eIF4E
chain_target = "A"
protein = "4E"
lower = 170
upper = 178
idps = glob.glob(f"{parent}/LIDR_{protein}_ANY_bb/*.pdb")
output_upper = f"{parent}/LIDR_{protein}_U/"
output_lower = f"{parent}/LIDR_{protein}_L/"

execute = partial(
    break_idr_attach,
    fld=pdb_fld,
    upper=upper,
    lower=lower,
    chain=chain_target,
    output_upper=output_upper,
    output_lower=False,
    )
execute_pool = pool_function(execute, idps, ncores=ncores)
for _ in execute_pool:
    pass

execute = partial(
    break_idr_attach,
    fld=pdb_fld,
    upper=upper,
    lower=lower,
    chain=chain_target,
    output_upper=False,
    output_lower=output_lower,
    )
execute_pool = pool_function(execute, idps, ncores=ncores)
for _ in execute_pool:
    pass

# For the L-IDR on 4E-BP2
chain_target = "B"
protein = "BP2"
lower = 26
upper = 45
idps = glob.glob(f"{parent}/LIDR_{protein}_ANY_bb/*.pdb")
output_upper = f"{parent}/LIDR_{protein}_U/"
output_lower = f"{parent}/LIDR_{protein}_L/"

execute = partial(
    break_idr_attach,
    fld=pdb_fld,
    upper=upper,
    lower=lower,
    chain=chain_target,
    output_upper=output_upper,
    output_lower=False,
    )
execute_pool = pool_function(execute, idps, ncores=ncores)
for _ in execute_pool:
    pass

execute = partial(
    break_idr_attach,
    fld=pdb_fld,
    upper=upper,
    lower=lower,
    chain=chain_target,
    output_upper=False,
    output_lower=output_lower,
    )
execute_pool = pool_function(execute, idps, ncores=ncores)
for _ in execute_pool:
    pass
```
