## supplemental_archive for "Local Disordered Region Sampling (LDRS) for Ensemble Modeling of Proteins with Experimentally Undetermined or Low Confidence Prediction Segments": BP24E_NIDR_CIDR_search.py

```
'''
Align and clash-check possible N- and C-IDRs
on the BP24E complex.
Author: Zi Hao Liu (@menoliu)
Date: July 17, 2023
'''
import glob
import re

import numpy as np

from functools import partial
from idpconfgen.libs.libmulticore import pool_function
from idpconfgen.libs.libstructure import (
    Structure,
    col_name,
    col_resSeq,
    col_chainID,
    cols_coords,
    parse_pdb_to_array,
    write_PDB,
    structure_to_pdb,
    )
from idpconfgen.ldrs_helper import (
    align_coords,
    count_clashes,
    )

from idpconfgen import Path


def linker_to_chain(idp, case, fld, chain, output, **kwargs):
    folded_domain = Path(fld)
    fld_struc = Structure(folded_domain)
    fld_struc.build()
    atom_names = fld_struc.data_array[:, col_name]
    chain_ids = fld_struc.data_array[:, col_chainID]
    fld_seq = fld_struc.data_array[:, col_resSeq].astype(int)
    chain_found = False
    
    fld_term_idx = {}
    for j, atom in enumerate(atom_names):
        if case == "C-IDR":
            k = len(atom_names) - 1 - j
            if chain_ids[k] == chain and chain_found == False:
                last_seq = fld_seq[k]
                chain_found = True
            if chain_found == True:
                seq = fld_seq[k]

                if seq == last_seq and atom_names[k] == "N":
                    fld_term_idx["N"] = k
                elif seq == last_seq and atom_names[k] == "CA":
                    fld_term_idx["CA"] = k
                elif seq == last_seq - 1 and atom_names[k] == "C":
                    fld_term_idx["C"] = k
                elif seq == last_seq - 2:
                    break
        elif case == "N-IDR":
            if chain_ids[j] == chain and chain_found == False:
                first_seq = fld_seq[j]
                chain_found = True
            if chain_found == True:
                seq = fld_seq[j]
                
                if seq == first_seq + 1 and atom == "N":
                    fld_term_idx["N"] = j
                elif seq == first_seq + 1 and atom == "CA":
                    fld_term_idx["CA"] = j
                elif seq == first_seq and atom == "C":
                    fld_term_idx["C"] = j
                elif seq == first_seq + 2:
                    break
                
    fld_Cxyz = fld_struc.data_array[fld_term_idx["C"]][cols_coords].astype(float).tolist()  # noqa: E501
    fld_Nxyz = fld_struc.data_array[fld_term_idx["N"]][cols_coords].astype(float).tolist()  # noqa: E501
    fld_CAxyz = fld_struc.data_array[fld_term_idx["CA"]][cols_coords].astype(float).tolist()  # noqa: E501
    # Coordinates of boundary to stitch to later on
    fld_coords = np.array([fld_Cxyz, fld_Nxyz, fld_CAxyz])
    
    with open(idp) as f:
        idp_data = f.read()
    try:
        idp_arr = parse_pdb_to_array(idp_data)
    except AssertionError:
        return
    aligned = align_coords(idp_arr, fld_coords, case)
    clash, frag = count_clashes(
        fragment=aligned,
        parent=fld_struc,
        case=case,
        max_clash=40,
        tolerance=0.4,
        )
    if type(clash) is bool:
        return

    struc = structure_to_pdb(frag)
    split = idp.split("/")
    number = int(re.search(r'\d+', split[-1]).group())
    
    write_PDB(struc, output + f"passed_{number}.pdb")


# Change custom variables here
ncores = 32
parent = "/PATH/TO/PARENT/FOLDER"
pdb_fld = f"{parent}/4UED_cleaned_both.pdb"


chain_target = "A"
protein = "4E"
case = "N-IDR"
idps = glob.glob(f"{parent}/{case}_{protein}_ANY_bb/*.pdb")
output_folder = f"{parent}/{case}_{protein}_passed/"

execute = partial(
    linker_to_chain,
    case=case,
    fld=pdb_fld,
    chain=chain_target,
    output=output_folder,
    )
execute_pool = pool_function(execute, idps, ncores=ncores)
for _ in execute_pool:
    pass

chain_target = "B"
protein = "BP2_canonical"
case = "N-IDR"

pdb_fld = f"{parent}/4UED_cleaned_both.pdb"
idps = glob.glob(f"{parent}/{case}_{protein}_ANY_bb/*.pdb")
output_folder = f"{parent}/{case}_{protein}_passed/"

execute = partial(
    linker_to_chain,
    case=case,
    fld=pdb_fld,
    chain=chain_target,
    output=output_folder,
    )
execute_pool = pool_function(execute, idps, ncores=ncores)
for _ in execute_pool:
    pass

chain_target = "B"
protein = "BP2_secondary"
case = "C-IDR"

pdb_fld = f"{parent}/4UED_cleaned_both.pdb"
idps = glob.glob(f"{parent}/{case}_{protein}_ANY_bb/*.pdb")
output_folder = f"{parent}/{case}_{protein}_passed/"

execute = partial(
    linker_to_chain,
    case=case,
    fld=pdb_fld,
    chain=chain_target,
    output=output_folder,
    )
execute_pool = pool_function(execute, idps, ncores=ncores)
for _ in execute_pool:
    pass
```
