## supplemental_archive for "Local Disordered Region Sampling (LDRS) for Ensemble Modeling of Proteins with Experimentally Undetermined or Low Confidence Prediction Segments": BP24E_stitching_both.py

```
'''
Clash-check and stitch all IDR fragments onto BP24E template.
Author: Zi Hao Liu (@menoliu)
Date: July 17, 2023
'''
import glob
import random
import numpy as np

from functools import partial
from multiprocessing import Queue
from idpconfgen.libs.libmulticore import pool_function
from idpconfgen.libs.libstructure import (
    Structure,
    col_chainID,
    col_serial,
    col_segid,
    col_resSeq,
    parse_pdb_to_array,
    write_PDB,
    structure_to_pdb,
    )
from idpconfgen.ldrs_helper import count_clashes

from idpconfgen import Path


CONF_NUMBER = Queue()


def check_clash_cinco(double_cinco):
    # `double_cinco` is a set of (cinco_arr, cinco_struc)
    # order is (n4e, b4e, nbp2, bbp2, cbp2)
    n4e_arr = double_cinco[0][0]
    b4e_arr = double_cinco[0][1]
    nbp2_arr = double_cinco[0][2]
    bbp2_arr = double_cinco[0][3]
    b4e_struc = double_cinco[1][1]
    nbp2_struc = double_cinco[1][2]
    bbp2_struc = double_cinco[1][3]
    cbp2_struc = double_cinco[1][4]
    
    n4e_b4e, _ = count_clashes(n4e_arr, b4e_struc)
    n4e_nbp2, _ = count_clashes(n4e_arr, nbp2_struc)
    n4e_bbp2, _ = count_clashes(n4e_arr, bbp2_struc)
    n4e_cbp2, _ = count_clashes(n4e_arr, cbp2_struc)
    b4e_nbp2, _ = count_clashes(b4e_arr, nbp2_struc)
    b4e_bbp2, _ = count_clashes(b4e_arr, bbp2_struc)
    b4e_cbp2, _ = count_clashes(b4e_arr, cbp2_struc)
    nbp2_bbp2, _ = count_clashes(nbp2_arr, bbp2_struc)
    nbp2_cbp2, _ = count_clashes(nbp2_arr, cbp2_struc)
    bbp2_cbp2, _ = count_clashes(bbp2_arr, cbp2_struc)

    if type(n4e_b4e) is bool or \
        type(n4e_nbp2) is bool or \
        type(n4e_bbp2) is bool or \
        type(n4e_cbp2) is bool or \
        type(b4e_nbp2) is bool or \
        type(b4e_bbp2) is bool or \
        type(b4e_cbp2) is bool or \
        type(nbp2_bbp2) is bool or \
        type(nbp2_cbp2) is bool or \
        type(bbp2_cbp2) is bool:
            return None
    
    return double_cinco[0]


def psurgeon_cinco(cinco_arr, fld_arr, output, rs):
    n4e = cinco_arr[0]
    seq_n4e = n4e[:, col_resSeq]
    
    b4e = cinco_arr[1]
    seq_b4e = b4e[:, col_resSeq]
    
    nbp2 = cinco_arr[2]
    seq_nbp2 = nbp2[:, col_resSeq]
    
    bbp2 = cinco_arr[3]
    seq_bbp2 = bbp2[:, col_resSeq]
    
    cbp2 = cinco_arr[4]
    seq_cbp2 = cbp2[:, col_resSeq]
    
    for i, _ in enumerate(n4e):
        n4e[i][col_chainID] = 'A'
        n4e[i][col_segid] = 'A'
    
    for i, _ in enumerate(b4e):
        b4e[i][col_chainID] = 'A'
        b4e[i][col_segid] = 'A'

    # Remove last residue on NIDR_4E
    for i, _ in enumerate(seq_n4e):
        j = len(seq_n4e) - 1 - i
        curr = seq_n4e[j]
        prev = seq_n4e[j - 1]
        n4e = n4e[:-1]
        if prev != curr:
            break
    
    # Remove first and last residue on LIDR_4E
    for i, seq in enumerate(seq_b4e):
            next = seq_b4e[i + 1]
            b4e = b4e[1:]
            if next != seq:
                break
    for i, _ in enumerate(seq_b4e):
        j = len(seq_b4e) - 1 - i
        curr = seq_b4e[j]
        prev = seq_b4e[j - 1]
        b4e = b4e[:-1]
        if prev != curr:
            break
   
    # Remove last residue on NIDR_BP2
    for i, _ in enumerate(seq_nbp2):
        j = len(seq_nbp2) - 1 - i
        curr = seq_nbp2[j]
        prev = seq_nbp2[j - 1]
        nbp2 = nbp2[:-1]
        if prev != curr:
            break
    
    # Remove first and last residue on LIDR_BP2
    for i, seq in enumerate(seq_bbp2):
            next = seq_bbp2[i + 1]
            bbp2 = bbp2[1:]
            if next != seq:
                break
    for i, _ in enumerate(seq_bbp2):
        j = len(seq_bbp2) - 1 - i
        curr = seq_bbp2[j]
        prev = seq_bbp2[j - 1]
        bbp2 = bbp2[:-1]
        if prev != curr:
            break
    
    # Remove first residue on CIDR_BP2
    for i, seq in enumerate(seq_cbp2):
            next = seq_cbp2[i + 1]
            cbp2 = cbp2[1:]
            if next != seq:
                break
    
    n4e_lst = n4e.tolist()
    fld_lst = fld_arr.tolist()
    
    # Correct residue numbering for LIDR_4E (+202)
    for i, line in enumerate(b4e):
        resnum = int(line[col_resSeq])
        b4e[i][col_resSeq] = str(resnum + 202)
    b4e_lst = b4e.tolist()
    
    # Correct residue numbering for LIDR_BP2 (+58)
    for i, line in enumerate(bbp2):
        resnum = int(line[col_resSeq])
        bbp2[i][col_resSeq] = str(resnum + 58)
        bbp2[i][col_chainID] = 'B'
        bbp2[i][col_segid] = 'B'
    bbp2_lst = bbp2.tolist()
    
    # Correct residue numbering for CIDR on BP2 (+80)
    for i, line in enumerate(cbp2):
        resnum = int(line[col_resSeq])
        cbp2[i][col_resSeq] = str(resnum + 80)
        cbp2[i][col_chainID] = 'B'
        cbp2[i][col_segid] = 'B'
    
    # Change chain ID to "B" for BP2 fragments
    for i, _ in enumerate(nbp2):
        nbp2[i][col_chainID] = 'B'
        nbp2[i][col_segid] = 'B'
    nbp2_lst = nbp2.tolist()
    cbp2_lst = cbp2.tolist()
    
    # Insert LIDR_4E, NIDR_BP2, and LIDR_BP2
    fld_seq = fld_arr[:, col_resSeq]
    for i, seq in enumerate(fld_seq):
        next_seq = fld_seq[i + 1]
        if int(next_seq) - int(seq) > 1:
            b4e_idx = i + 1
            break
    for i, row in enumerate(b4e_lst):
        fld_lst.insert(b4e_idx + i, row)

    fld_arr = np.array(fld_lst)
    fld_chain = fld_arr[:, col_chainID]
    for i, _ in enumerate(fld_chain):
        next_chain = fld_chain[i + 1]
        if next_chain == "B":
            nbp2_idx = i + 1
            break
    for i, row in enumerate(nbp2_lst):
        fld_lst.insert(nbp2_idx + i, row)
    
    fld_arr = np.array(fld_lst)
    fld_seq = fld_arr[:, col_resSeq]
    fld_chain = fld_arr[:, col_chainID]
    for i, seq in enumerate(fld_seq):
        next_seq = fld_seq[i + 1]
        if int(next_seq) - int(seq) > 1 and fld_chain[i + 1] == "B":
            bbp2_idx = i + 1
            break
    for i, row in enumerate(bbp2_lst):
        fld_lst.insert(bbp2_idx + i, row)

    new_struc_lst = n4e_lst + fld_lst + cbp2_lst
    new_struc_arr = np.array(new_struc_lst)
    new_serial = [str(i) for i in range(1, len(new_struc_arr) + 1)]
    new_struc_arr[:, col_serial] = new_serial
    
    new_struc = structure_to_pdb(new_struc_arr)
    
    write_PDB(new_struc, output + f"conformer_rs{rs}_{CONF_NUMBER.get()}.pdb")
        
# Change variables here
parent_path = "/PATH/TO/PARENT/FOLDER"
nconfs = 419
rs = 0
ncores = 32
random.seed(rs)

output_path = f"{parent_path}/BP24E_4UED_both/"
n4e_paths = glob.glob(f"{parent_path}/NIDR_4E_mcsce/*.pdb")
nbp2_canon_paths = glob.glob(f"{parent_path}/NIDR_BP2_canon_mcsce/*.pdb")
nbp2_second_paths = glob.glob(f"{parent_path}/NIDR_BP2_second_mcsce/*.pdb")
b4e_paths = glob.glob(f"{parent_path}/LIDR_4E_mcsce/*.pdb")
bbp2_paths = glob.glob(f"{parent_path}/LIDR_BP2_mcsce/*.pdb")
ccanon_paths = glob.glob(f"{parent_path}/CIDR_BP2_canon_mcsce/*.pdb")
csecond_paths = glob.glob(f"{parent_path}/CIDR_BP2_second_mcsce/*.pdb")
both_fld = f"{parent_path}/PDB_prepped/4UED_both_prepped.pdb"
canon_fld = f"{parent_path}/PDB_prepped/4UED_canon_prepped.pdb"
second_fld = f"{parent_path}/PDB_prepped/4UED_secondary_prepped.pdb"

for i in range(1, nconfs + 1):
    CONF_NUMBER.put(i)

both_fld_struc = Structure(Path(both_fld))
both_fld_struc.build()
both_arr = both_fld_struc.data_array

canon_fld_struc = Structure(Path(canon_fld))
canon_fld_struc.build()
canon_arr = canon_fld_struc.data_array

second_fld_struc = Structure(Path(second_fld))
second_fld_struc.build()
second_arr = second_fld_struc.data_array

n4e_arr = []
n4e_struc = []
nbp2_arr = []
nbp2_struc = []
b4e_arr = []
b4e_struc = []
bbp2_arr = []
bbp2_struc = []
cbp2_arr = []
cbp2_struc = []

cinco_check = []
final_cinco_arr = []

random.shuffle(n4e_paths)
random.shuffle(nbp2_canon_paths)
random.shuffle(nbp2_second_paths)
random.shuffle(b4e_paths)
random.shuffle(bbp2_paths)
random.shuffle(ccanon_paths)
random.shuffle(csecond_paths)

n4e_paths = n4e_paths[:nconfs]
nbp2_canon_paths = nbp2_canon_paths[:nconfs]
nbp2_second_paths = nbp2_second_paths[:nconfs]
b4e_paths = b4e_paths[:nconfs]
bbp2_paths = bbp2_paths[:nconfs]
ccanon_paths = ccanon_paths[:nconfs]
csecond_paths = csecond_paths[:nconfs]

cinco_paths = []

# Both case
for i, nidr in enumerate(n4e_paths):
    cinco_paths.append([nidr, b4e_paths[i], nbp2_canon_paths[i], bbp2_paths[i], csecond_paths[i]])

for p in cinco_paths:
    p_n4e = p[0]
    p_b4e = p[1]
    p_nbp2 = p[2]
    p_bbp2 = p[3]
    p_cbp2 = p[4]
    
    with open(p_n4e) as n4e:
        data_n4e = n4e.read()
    with open(p_b4e) as b4e:
        data_b4e = b4e.read()
    with open(p_nbp2) as nbp2:
        data_nbp2 = nbp2.read()
    with open(p_bbp2) as bbp2:
        data_bbp2 = bbp2.read()
    with open(p_cbp2) as cbp2:
        data_cbp2 = cbp2.read()
    
    try:
        arr = parse_pdb_to_array(data_n4e)
        struc = Structure(Path(p_n4e))
        struc.build()
        
        n4e_arr.append(arr)
        n4e_struc.append(struc)
        
        arr = parse_pdb_to_array(data_b4e)
        struc = Structure(Path(p_b4e))
        struc.build()
        
        b4e_arr.append(arr)
        b4e_struc.append(struc)
        
        arr = parse_pdb_to_array(data_nbp2)
        struc = Structure(Path(p_nbp2))
        struc.build()
        
        nbp2_arr.append(arr)
        nbp2_struc.append(struc)
        
        arr = parse_pdb_to_array(data_bbp2)
        struc = Structure(Path(p_bbp2))
        struc.build()
        
        bbp2_arr.append(arr)
        bbp2_struc.append(struc)
        
        arr = parse_pdb_to_array(data_cbp2)
        struc = Structure(Path(p_cbp2))
        struc.build()
        
        cbp2_arr.append(arr)
        cbp2_struc.append(struc)
    except AssertionError:
        continue

for i, _ in enumerate(cinco_paths):
    cinco_arr = ([n4e_arr[i], b4e_arr[i], nbp2_arr[i], bbp2_arr[i], cbp2_arr[i]])
    cinco_struc = [n4e_struc[i], b4e_struc[i], nbp2_struc[i], bbp2_struc[i], cbp2_struc[i]]
    cinco_check.append((cinco_arr, cinco_struc))


execute_clash = pool_function(check_clash_cinco, cinco_check, ncores=ncores)
for result in execute_clash:
    if result is not None:
        final_cinco_arr.append(result)

execute = partial(
    psurgeon_cinco,
    fld_arr=both_arr,
    output=output_path,
    rs=rs
    )
execute_pool = pool_function(execute, final_cinco_arr, ncores=ncores)
for _ in execute_pool:
    pass
```
