## supplemental_archive for "Local Disordered Region Sampling (LDRS) for Ensemble Modeling of Proteins with Experimentally Undetermined or Low Confidence Prediction Segments": move_and_rename.py

```
'''
Move or copy "X" number of files to another
folder while renaming them.
Author: Zi Hao Liu (@menoliu)
Date: July 6, 2023
'''
import os
import shutil


def move_and_rename_files(source_folder, destination_folder, num_files, option="copy"):
    file_list = os.listdir(source_folder)
    num_files = min(num_files, len(file_list))
    
    if option not in ("move", "copy"):
        print(f"Invalide mode {option}. Please change to either move or copy.")
        return

    for i in range(num_files):
        file_name = file_list[i]
        source_path = os.path.join(source_folder, file_name)

        # Generate the new file name
        new_file_name = f"conformer_{i+1}.pdb"  # Replace with your desired naming convention

        # Generate the new file path in the destination folder
        destination_path = os.path.join(destination_folder, new_file_name)
        
        if option == "move":
            # Move the file and rename it
            shutil.move(source_path, destination_path)

            # Uncomment the below line to print every move to console
            #print(f"Moved '{file_name}' to '{new_file_name}'.")
        else:
            # Move the file and rename it
            shutil.copy(source_path, destination_path)
        
            # Uncomment the below line to print every copy to console
            #print(f"Copied '{file_name}' to '{new_file_name}'.")  


# Change the following variables to your requirements
source_folder = "/PATH/TO/SOURCE"
destination_folder = "/PATH/TO/DESTINATION"
num_files_to_move = 1500

move_and_rename_files(source_folder, destination_folder, num_files_to_move)
```
