## supplemental_archive for "Local Disordered Region Sampling (LDRS) for Ensemble Modeling of Proteins with Experimentally Undetermined or Low Confidence Prediction Segments": plot_ramachandran.py

```
'''
Plot Ramachandran plot from output of IDPConformerGenerator torsions sublcient.
Author: Zi Hao Liu (@menoliu)
Date: July 10, 2023
'''
import matplotlib.pyplot as plt
import json
import math


angles = {"omega": [], "phi": [], "psi": []}


def plot_ramachandran(angles, output):
    phi = angles["phi"]
    psi = angles["psi"]

    # Feel free to change plotting parameters here
    plt.scatter(phi, psi, s=0.1, c='black', marker='o', alpha=0.1)
    plt.xlabel("Phi")
    plt.ylabel("Psi")
    plt.savefig(output, dpi=300, transparent=True, bbox_inches='tight')
    plt.show()


# Change path to torsions.json file generated by `idpconfgen torsions` subclient
# Degrees vs radians are changed within the torsions subclient
path_to_torsions = '/PATH/TO/TORSIONS.json'
output_png = '/PATH/TO/IMAGE.png'

with open(path_to_torsions) as j:
    data = json.load(j)
    for conformer in data:
        angles["phi"].extend(data[conformer]["phi"])
        angles["psi"].extend(data[conformer]["psi"])
    
plot_ramachandran(angles, output_png)
```
