## supplemental_archive for "Local Disordered Region Sampling (LDRS) for Ensemble Modeling of Proteins with Experimentally Undetermined or Low Confidence Prediction Segments": remove_duplicates.py

```
'''
Faster way to remove duplicate files in a folder.
Author: Zi Hao Liu (@menoliu)
Date: July 6, 2023
'''
import os
import hashlib


def remove_duplicates(folder_path):
    # Get a list of all files in the folder
    file_list = os.listdir(folder_path)
    
    # Create a dictionary to store file content hashes
    content_hashes = {}
    
    # Iterate over each file in the folder
    for file_name in file_list:
        file_path = os.path.join(folder_path, file_name)
        
        # Calculate the content hash of the file
        with open(file_path, 'rb') as file:
            file_content = file.read()
            content_hash = hashlib.md5(file_content).hexdigest()
        
        # Check if a file with the same content hash already exists
        if content_hash in content_hashes:
            # Delete the current file
            os.remove(file_path)
        else:
            # Store the content hash and file path in the dictionary
            content_hashes[content_hash] = file_path


# Change variables here
path_to_folder = "/CHANGE/PATH/HERE"
remove_duplicates(path_to_folder)
```
