## supplemental_archive for "Local Disordered Region Sampling (LDRS) for Ensemble Modeling of Proteins with Experimentally Undetermined or Low Confidence Prediction Segments": remove_lowconfidence_residues.py

```
'''
Remove residues in a PDB file based on values in the B-factor
column. User-editable confidence score thresholds.
Author: Zi Hao Liu (@menoliu)
Date: July 13, 2023
'''
import numpy as np

from idpconfgen.libs.libstructure import (
    Structure,
    col_serial,
    col_temp,
    structure_to_pdb,
    )
from idpconfgen import Path


def remove_residues(pdb_path, threshold, output):
    pdb_struc = Structure(Path(pdb_path))
    pdb_struc.build()
    
    pdb_arr = pdb_struc.data_array
    pdb_lst = pdb_arr.tolist()
    scores = pdb_arr[:, col_temp].astype(float)
    
    new_structure_lst = []

    for i, s in enumerate(scores):
        # Change the comparison operator here if needed
        # need to be changed to '<' for RoseTTAFold structures
        if s > threshold:
            new_structure_lst.append(pdb_lst[i])

    new_struc_arr = np.array(new_structure_lst)
    # Fix serial numbers
    new_serial = [str(i) for i in range(1, len(new_struc_arr) + 1)]
    new_struc_arr[:, col_serial] = new_serial
    
    new_struc = structure_to_pdb(new_struc_arr)
    with open(output, 'w') as f:
        for line in new_struc:
            f.write(line + "\n")


# Example usage shown below
input_file = "/PATH/TO/PREDICTED.pdb"
output_file = "/PATH/TO/PROCESSED.pdb"
# Recommended 70 for AlphaFold structures (pLDDT)
# Recommended 0.7 for ESMFold structures (pLDDT)
# Recommended 5.0 for RoseTTAFold full-model structures (Angstroms)
threshold = 70
remove_residues(input_file, threshold, output_file)
```
