## supplemental_archive for "Local Disordered Region Sampling (LDRS) for Ensemble Modeling of Proteins with Experimentally Undetermined or Low Confidence Prediction Segments": split_chains.py

```
'''
Split protein complex into distinct chains.
Currently for chain A and B.
Author: Zi Hao Liu (@menoliu)
Date: July 11, 2023
'''
import glob
import numpy as np

from idpconfgen.libs.libstructure import (
    Structure,
    col_chainID,
    structure_to_pdb,
    )
from idpconfgen import Path


def split_chains(pdb_path, output):
    pdb_struc = Structure(Path(pdb_path))
    pdb_struc.build()
    
    pdb_arr = pdb_struc.data_array
    pdb_lst = pdb_arr.tolist()
    pdb_chain = pdb_arr[:, col_chainID]
    
    chain_A = []
    chain_B = []
    
    # Split the chains into separate lists
    # Can redefine chain ID here for your purposes
    for i, chain in enumerate(pdb_chain):
        if chain == "A":
            chain_A.append(pdb_lst[i])
        elif chain == "B":
            chain_B.append(pdb_lst[i])
    
    chain_A_arr = np.array(chain_A)
    chain_B_arr = np.array(chain_B)
    
    pdb_path_split = pdb_path.split("/")
    pdb_name = pdb_path_split[-1]
    
    chain_A_struc = structure_to_pdb(chain_A_arr)
    chain_B_struc = structure_to_pdb(chain_B_arr)
    # Please note you need to have a folder called "chainA" and "chainB"
    # Can also edit these folders to whatever you like
    output_A = output + f"/chainA/A_{pdb_name}"
    output_B = output + f"/chainB/B_{pdb_name}"
    with open(output_A, 'w') as f:
        for line in chain_A_struc:
            f.write(line + "\n")
    with open(output_B, 'w') as f:
        for line in chain_B_struc:
            f.write(line + "\n")


# Example usage shown below, will iterate through each PDB file
original_pdb_paths = glob.glob("/PATH/TO/COMPLEXES/*.pdb")
output_folder = "/PATH/TO/OUTPUT/FOLDER"
for pdb in original_pdb_paths:
    split_chains(pdb, output_folder)
```
