## supplemental_archive for "Local Disordered Region Sampling (LDRS) for Ensemble Modeling of Proteins with Experimentally Undetermined or Low Confidence Prediction Segments": split_conformers.py

```
'''
Split a large number of conformers to managable
chunks to set up parallel processing.
Author: Zi Hao Liu (@menoliu)
Date: July 6, 2023
'''
import glob
import shutil


# Number to split by (e.g. if we have 6000 total conformers, this will split them
# into two sets of 3000 conformers
n_splitted = 3000

# Path to conformers
conformers = glob.glob("/PATH/TO/CONFORMERS/*.pdb")

# Please note you must create the destination folders ahead of time
# destination_0, destination_1, etc.
destination = "/PATH/TO/FOLDER"

splitted = [conformers[x:x+n_splitted] for x in range (0, len(conformers), n_splitted)]

for c, chunk in enumerate(splitted):
    for file in chunk:
        shutil.move(file, f"{destination}_{c}")
```
