## supplemental_archive for "Local Disordered Region Sampling (LDRS) for Ensemble Modeling of Proteins with Experimentally Undetermined or Low Confidence Prediction Segments": p53_LIDR_processing.py

```
'''
Processes the first linker IDR in p53.
Author: Zi Hao Liu (@menoliu)
Date: July 17, 2023
'''
import re
import glob
import numpy as np

from functools import partial
from idpconfgen.libs.libmulticore import pool_function
from idpconfgen.libs.libstructure import (
    Structure,
    col_name,
    col_resSeq,
    cols_coords,
    parse_pdb_to_array,
    write_PDB,
    structure_to_pdb,
    )
from idpconfgen.ldrs_helper import (
    align_coords,
    count_clashes,
    next_seeker,
    )

from idpconfgen import Path


def break_idr_attach(idr, fld, upper, lower, output_upper=False, output_lower=False):
    fld_struc = Structure(Path(fld))
    fld_struc.build()
    atom_names = fld_struc.data_array[:, col_name]
    
    fld_seq = fld_struc.data_array[:, col_resSeq].astype(int)
    first_seq = fld_seq[0]
    

    if first_seq > 1:
        fld_seq -= first_seq - 1
        first_seq = fld_seq[0]
    
    fld_term_idx = {}
    for s, aa in enumerate(fld_seq):
        if aa == lower - 1 and atom_names[s] == 'C':
            fld_term_idx["CL"] = s
        elif aa == lower:
            if atom_names[s] == 'N':
                fld_term_idx["NL"] = s
            elif atom_names[s] == 'CA':
                fld_term_idx["CAL"] = s
        elif aa == upper and atom_names[s] == 'C':
            fld_term_idx['CU'] = s
        elif aa == upper + 1:
            if atom_names[s] == 'N':
                fld_term_idx['NU'] = s
            elif atom_names[s] == 'CA':
                fld_term_idx['CAU'] = s
        if aa > upper + 1:
            break
    
    fld_CLxyz = fld_struc.data_array[fld_term_idx["CL"]][cols_coords].astype(float)
    fld_NLxyz = fld_struc.data_array[fld_term_idx["NL"]][cols_coords].astype(float)
    fld_CALxyz = fld_struc.data_array[fld_term_idx["CAL"]][cols_coords].astype(float)
    fld_coords_L = np.array([fld_CLxyz, fld_NLxyz, fld_CALxyz])
    
    fld_CUxyz = fld_struc.data_array[fld_term_idx["CU"]][cols_coords].astype(float)
    fld_NUxyz = fld_struc.data_array[fld_term_idx["NU"]][cols_coords].astype(float)
    fld_CAUxyz = fld_struc.data_array[fld_term_idx["CAU"]][cols_coords].astype(float)
    fld_coords_U = np.array([fld_CUxyz, fld_NUxyz, fld_CAUxyz])
    
    with open(idr) as f:
        idr_data = f.read()
    try:
        idr_arr = parse_pdb_to_array(idr_data)
    except AssertionError:
        return
    
    if output_lower:
        aligned_break_L = align_coords(idr_arr, fld_coords_L, "C-IDR")
        clashesL, fragmentL = count_clashes(
            aligned_break_L,
            fld_struc,
            "C-IDR",
            max_clash=80,
            tolerance=0.4
            )
        if type(clashesL) is int:
            aligned_struc = structure_to_pdb(fragmentL)
            idr_split = idr.split("/")
            idr_int = int(re.search(r'\d+', idr_split[-1]).group())
            write_PDB(aligned_struc, output_lower + f"aligned_conf_L_{idr_int}.pdb")
    
    elif output_upper:
        aligned_break_U = align_coords(idr_arr, fld_coords_U, "N-IDR")
        clashesU, fragmentU = count_clashes(
            aligned_break_U,
            fld_struc,
            "N-IDR",
            max_clash=80,
            tolerance=0.4
            )
        if type(clashesU) is int:
            aligned_struc = structure_to_pdb(fragmentU)
            idr_split = idr.split("/")
            idr_int = int(re.search(r'\d+', idr_split[-1]).group())
            write_PDB(aligned_struc, output_upper + f"aligned_conf_U_{idr_int}.pdb")


# Change variables here
# LIDR1 is the first linker IDR while LIDR2 is the second.
# Upper and lower and the residue number ranges.
parent_folder = "/PATH/TO/PARENT/FOLDER"
case = "LIDR1"
ncores = 32
lower = 87
upper = 92

''' For the second linker IDR
case = "LIDR2"
lower = 197
upper = 229
'''

idr_lst = glob.glob(f"{parent_folder}/p53_{case}_ANY_bb/*.pdb")
idr_upper = glob.glob(f"{parent_folder}/{case}_U/*.pdb")
idr_lower = glob.glob(f"{parent_folder}/{case}_L/*.pdb")
fld = f"{parent_folder}/p53_cleaned.pdb"
output_upper = f"{parent_folder}/{case}_U/"
output_lower = f"{parent_folder}/{case}_L/"
output_combos = f"{parent_folder}/{case}_results/"

# Align and clach-check the upper, or C-term IDR
execute = partial(
    break_idr_attach,
    fld=fld,
    upper=upper,
    lower=lower,
    output_upper=output_upper,
    output_lower=False,
    )
execute_pool = pool_function(execute, idr_lst, ncores=ncores)
for _ in execute_pool:
    pass

# Align and clash-check the lower, or N-term IDR
execute = partial(
    break_idr_attach,
    fld=fld,
    upper=upper,
    lower=lower,
    output_upper=False,
    output_lower=output_lower,
    )
execute_pool = pool_function(execute, idr_lst, ncores=ncores)
for _ in execute_pool:
    pass

# Perform next-seeker on upper and lower IDR fragments
execute = partial(
    next_seeker,
    nterm_idr_lib=idr_upper,
    max_clash=80,
    tolerance=0.8,
    output_folder=output_combos,
    )
execute_pool = pool_function(execute, idr_lower, ncores=ncores)
for _ in execute_pool:
    pass
```
