## supplemental_archive for "Local Disordered Region Sampling (LDRS) for Ensemble Modeling of Proteins with Experimentally Undetermined or Low Confidence Prediction Segments": p53_NIDR+CIDR_processing.py

```
'''
Processes the N- and C-IDRs in p53.
Author: Zi Hao Liu (@menoliu)
Date: July 17, 2023
'''
import re
import glob
import numpy as np

from functools import partial
from idpconfgen.libs.libmulticore import pool_function
from idpconfgen.libs.libstructure import (
    Structure,
    col_name,
    col_resSeq,
    cols_coords,
    parse_pdb_to_array,
    write_PDB,
    structure_to_pdb,
    )
from idpconfgen.ldrs_helper import (
    align_coords,
    count_clashes,
    )

from idpconfgen import Path


def term_idr_attach(idr, fld_struc, case, output):
    # Folded structure should already be built
    fld_name = fld_struc.data_array[:, col_name]
    fld_seq = fld_struc.data_array[:, col_resSeq].astype(int)
    
    first_seq = fld_seq[0]
    last_seq = fld_seq[-1]
    
    fld_term_idx = {}
    if case == "NIDR":
        case = "N-IDR"
        for j, atom in enumerate(fld_name):
            curr_seq = fld_seq[j]
            if curr_seq == first_seq + 1 and atom == "N":
                fld_term_idx["N"] = j
            elif curr_seq == first_seq + 1 and atom == "CA":
                fld_term_idx["CA"] = j
            elif curr_seq == first_seq and atom == "C":
                fld_term_idx["C"] = j
            elif curr_seq == first_seq + 2:
                break
    elif case == "CIDR":
        case = "C-IDR"
        for j, _atom in enumerate(fld_name):
            k = len(fld_name) - 1 - j
            curr_seq = fld_seq[k]
            if curr_seq == last_seq and fld_name[k] == "N":
                fld_term_idx["N"] = k
            elif curr_seq == last_seq and fld_name[k] == "CA":
                fld_term_idx["CA"] = k
            elif curr_seq == last_seq - 1 and fld_name[k] == "C":
                fld_term_idx["C"] = k
            elif curr_seq == last_seq - 2:
                break
    
    fld_Cxyz = fld_struc.data_array[fld_term_idx["C"]][cols_coords].astype(float)  # noqa: E501
    fld_Nxyz = fld_struc.data_array[fld_term_idx["N"]][cols_coords].astype(float)  # noqa: E501
    fld_CAxyz = fld_struc.data_array[fld_term_idx["CA"]][cols_coords].astype(float)  # noqa: E501
    # Coordinates of boundary to stitch to later on
    fld_coords = np.array([fld_Cxyz, fld_Nxyz, fld_CAxyz])
    
    with open(idr) as f:
        idr_data = f.read()
    try:
        idr_arr = parse_pdb_to_array(idr_data)
    except AssertionError:
        return
    
    aligned_idr = align_coords(idr_arr, fld_coords, case)
    clashes, fragment = count_clashes(
        aligned_idr,
        fld_struc,
        case,
        max_clash=80,
        tolerance=0.4
        )
    if type(clashes) is int:
        aligned_struc = structure_to_pdb(fragment)
        idr_split = idr.split("/")
        idr_int = int(re.search(r'\d+', idr_split[-1]).group())
        write_PDB(aligned_struc, output + f"aligned_{case}_{idr_int}.pdb")


# Change parent folder here.
# Make sure the folders below already exist
parent_folder = "/PATH/TO/PARENT/FOLDER"
ncores = 32

case = "NIDR"
idr_lst = glob.glob(f"/{parent_folder}/p53_{case}_ANY_bb/*.pdb")
fld = f"{parent_folder}/p53_cleaned.pdb"
fld_struc = Structure(Path(fld))
fld_struc.build()
output = f"{parent_folder}/{case}_results/"

execute = partial(
    term_idr_attach,
    fld_struc=fld_struc,
    case=case,
    output=output,
    )
execute_pool = pool_function(execute, idr_lst, ncores=ncores)
for _ in execute_pool:
    pass

case = "CIDR"
idr_lst = glob.glob(f"/{parent_folder}/p53_{case}_ANY_bb/*.pdb")
fld = f"{parent_folder}/p53_cleaned.pdb"
fld_struc = Structure(Path(fld))
fld_struc.build()
output = f"{parent_folder}/{case}_results/"

execute = partial(
    term_idr_attach,
    fld_struc=fld_struc,
    case=case,
    output=output,
    )
execute_pool = pool_function(execute, idr_lst, ncores=ncores)
for _ in execute_pool:
    pass
```
