## supplemental_archive for "Local Disordered Region Sampling (LDRS) for Ensemble Modeling of Proteins with Experimentally Undetermined or Low Confidence Prediction Segments": p53_stitching_bb.py

```
'''
Stitch all fragments onto p53 template.
Author: Zi Hao Liu (@menoliu)
Date: July 17, 2023
'''
import glob
import random
import numpy as np

from functools import partial
from multiprocessing import Queue
from idpconfgen.libs.libmulticore import pool_function
from idpconfgen.libs.libstructure import (
    Structure,
    col_serial,
    col_resSeq,
    parse_pdb_to_array,
    write_PDB,
    structure_to_pdb,
    )
from idpconfgen.ldrs_helper import count_clashes

from idpconfgen import Path


CONF_NUMBER = Queue()


def check_clash_quattro(double_quattro):
    # `double_quattro` is a set of (quattro_arr, quattro_struc)
    # order is (nidr, bidr1, bidr2, cidr)
    nidr_arr = double_quattro[0][0]
    bidr1_arr = double_quattro[0][1]
    cidr_arr = double_quattro[0][3]
    bidr1_struc = double_quattro[1][1]
    bidr2_struc = double_quattro[1][2]
    cidr_struc = double_quattro[1][3]
    
    nb1, _ = count_clashes(nidr_arr, bidr1_struc)
    nb2, _ = count_clashes(nidr_arr, bidr2_struc)
    nc, _ = count_clashes(nidr_arr, cidr_struc)
    b1b2, _ = count_clashes(bidr1_arr, bidr2_struc)
    b1c, _ = count_clashes(bidr1_arr, cidr_struc)
    b2c, _ = count_clashes(cidr_arr, bidr2_struc)

    if type(nb1) is bool or \
        type(nb2) is bool or \
        type(nc) is bool or \
        type(b1b2) is bool or \
        type(b1c) is bool or \
        type(b2c) is bool:
            return None
    
    return double_quattro[0]


def psurgeon_quattro(quattro_arr, fld_arr, output, rs):
    nidr = quattro_arr[0]
    seq_nidr = nidr[:, col_resSeq]
    
    bidr1 = quattro_arr[1]
    seq_bidr1 = bidr1[:, col_resSeq]
    
    bidr2 = quattro_arr[2]
    seq_bidr2 = bidr2[:, col_resSeq]
    
    cidr = quattro_arr[3]
    seq_cidr = cidr[:, col_resSeq]
    
    # Remove last residue on NIDR
    for i, _ in enumerate(seq_nidr):
        j = len(seq_nidr) - 1 - i
        curr = seq_nidr[j]
        prev = seq_nidr[j - 1]
        nidr = nidr[:-1]
        if prev != curr:
            break
    
    # Remove first and last residue on BIDR1 and BIDR2
    for i, seq in enumerate(seq_bidr1):
            next = seq_bidr1[i + 1]
            bidr1 = bidr1[1:]
            if next != seq:
                break
    for i, _ in enumerate(seq_bidr1):
        j = len(seq_bidr1) - 1 - i
        curr = seq_bidr1[j]
        prev = seq_bidr1[j - 1]
        bidr1 = bidr1[:-1]
        if prev != curr:
            break
    
    for i, seq in enumerate(seq_bidr2):
            next = seq_bidr2[i + 1]
            bidr2 = bidr2[1:]
            if next != seq:
                break
    for i, _ in enumerate(seq_bidr2):
        j = len(seq_bidr2) - 1 - i
        curr = seq_bidr2[j]
        prev = seq_bidr2[j - 1]
        bidr2 = bidr2[:-1]
        if prev != curr:
            break
    
    # Remove first residue on CIDR
    for i, seq in enumerate(seq_cidr):
            next = seq_cidr[i + 1]
            cidr = cidr[1:]
            if next != seq:
                break
    
    nidr_lst = nidr.tolist()
    fld_lst = fld_arr.tolist()
    
    # Correct residue numbering for BIDR1 (+180)
    for i, line in enumerate(bidr1):
        resnum = int(line[col_resSeq])
        bidr1[i][col_resSeq] = str(resnum + 180)
    bidr1_lst = bidr1.tolist()
    # Insert BIDR1
    fld_seq = fld_arr[:, col_resSeq]
    for i, seq in enumerate(fld_seq):
        next_seq = fld_seq[i + 1]
        if int(next_seq) - int(seq) > 1:
            bidr1_idx = i + 1
            break
    for i, row in enumerate(bidr1_lst):
        fld_lst.insert(bidr1_idx + i, row)
    
    # Correct residue numbering for BIDR2 (+290)
    for i, line in enumerate(bidr2):
        resnum = int(line[col_resSeq])
        bidr2[i][col_resSeq] = str(resnum + 290)
    bidr2_lst = bidr2.tolist()
    # Insert BIDR2
    fld_arr = np.array(fld_lst)
    fld_seq = fld_arr[:, col_resSeq]
    for i, seq in enumerate(fld_seq):
        next_seq = fld_seq[i + 1]
        if int(next_seq) - int(seq) > 1:
            bidr2_idx = i + 1
            break
    for i, row in enumerate(bidr2_lst):
        fld_lst.insert(bidr2_idx + i, row)
    
    # Correct residue numbering for CIDR (+353)
    for i, line in enumerate(cidr):
        resnum = int(line[col_resSeq])
        cidr[i][col_resSeq] = str(resnum + 353)
    cidr_lst = cidr.tolist()
    
    new_struc_lst = nidr_lst + fld_lst + cidr_lst
    new_struc_arr = np.array(new_struc_lst)
    new_serial = [str(i) for i in range(1, len(new_struc_arr) + 1)]
    new_struc_arr[:, col_serial] = new_serial
    
    new_struc = structure_to_pdb(new_struc_arr)
    
    write_PDB(new_struc, output + f"conformer_rs{rs}_{CONF_NUMBER.get()}.pdb")
        
# Change variables here
parent_path = "/PATH/TO/PARENT/FOLDER"
ncores = 32
nconfs = 3300
rs = 0
random.seed(rs)

output_path = f"{parent_path}/p53_conformers_bb/"
nidr_paths = glob.glob(f"{parent_path}/NIDR_results/*.pdb")
bidr1_paths = glob.glob(f"{parent_path}/BIDR1_results/*.pdb")
bidr2_paths = glob.glob(f"{parent_path}/BIDR2_results/*.pdb")
cidr_paths = glob.glob(f"{parent_path}/CIDR_results/*.pdb")
fld_path = f"{parent_path}/p53_prepped.pdb"

for i in range(1, nconfs + 1):
    CONF_NUMBER.put(i)

fld_struc = Structure(Path(fld_path))
fld_struc.build()
fld_arr = fld_struc.data_array

nidr_arr = []
nidr_struc = []
bidr1_arr = []
bidr1_struc = []
bidr2_arr = []
bidr2_struc = []
cidr_arr = []
cidr_struc = []

quattro_check = []
final_quattro_arr = []

random.shuffle(nidr_paths)
random.shuffle(bidr1_paths)
random.shuffle(bidr2_paths)
random.shuffle(cidr_paths)
nidr_paths = nidr_paths[:nconfs]
bidr1_paths = bidr1_paths[:nconfs]
bidr2_paths = bidr2_paths[:nconfs]
cidr_paths = cidr_paths[:nconfs]

quattro_paths = []

for i, nidr in enumerate(nidr_paths):
    quattro_paths.append([nidr, bidr1_paths[i], bidr2_paths[i], cidr_paths[i]])

for p in quattro_paths:
    p_nidr = p[0]
    p_bidr1 = p[1]
    p_bidr2 = p[2]
    p_cidr = p[3]
    
    with open(p_nidr) as nidr:
        data_nidr = nidr.read()
    with open(p_bidr1) as bidr1:
        data_bidr1 = bidr1.read()
    with open(p_bidr2) as bidr2:
        data_bidr2 = bidr2.read()
    with open(p_cidr) as cidr:
        data_cidr = cidr.read()
    
    try:
        arr = parse_pdb_to_array(data_nidr)
        struc = Structure(Path(p_nidr))
        struc.build()
        
        nidr_arr.append(arr)
        nidr_struc.append(struc)
        
        arr = parse_pdb_to_array(data_bidr1)
        struc = Structure(Path(p_bidr1))
        struc.build()
        
        bidr1_arr.append(arr)
        bidr1_struc.append(struc)
        
        arr = parse_pdb_to_array(data_bidr2)
        struc = Structure(Path(p_bidr2))
        struc.build()
        
        bidr2_arr.append(arr)
        bidr2_struc.append(struc)
        
        arr = parse_pdb_to_array(data_cidr)
        struc = Structure(Path(p_cidr))
        struc.build()
        
        cidr_arr.append(arr)
        cidr_struc.append(struc)
    except AssertionError:
        continue

quattro_arr = []

for i, _ in enumerate(quattro_paths):
    trio_arr = ([nidr_arr[i], bidr1_arr[i], bidr2_arr[i], cidr_arr[i]])
    trio_struc = [nidr_struc[i], bidr1_struc[i], bidr2_struc[i], cidr_struc[i]]
    quattro_check.append((trio_arr, trio_struc))


execute_clash = pool_function(check_clash_quattro, quattro_check, ncores=ncores)
for result in execute_clash:
    if result is not None:
        final_quattro_arr.append(result)

execute = partial(
    psurgeon_quattro,
    fld_arr=fld_arr,
    output=output_path,
    rs=rs
    )
execute_pool = pool_function(execute, final_quattro_arr, ncores=ncores)
for _ in execute_pool:
    pass
```
