## supplemental_archive for "Local Disordered Region Sampling (LDRS) for Ensemble Modeling of Proteins with Experimentally Undetermined or Low Confidence Prediction Segments": A9_stitching.py

```
'''
Sitches LIDR backbone fragments for SLC26A9 STAS domain.
Author: Zi Hao Liu (@menoliu)
Date: July 17, 2023
'''
import glob
import random

from functools import partial
from multiprocessing import Queue
from idpconfgen.libs.libmulticore import pool_function
from idpconfgen.libs.libstructure import (
    Structure,
    col_resSeq,
    col_name,
    cols_coords,
    structure_to_pdb,
    )
from idpconfgen.core.definitions import vdW_radii_tsai_1999
from idpconfgen.ldrs_helper import psurgeon, calculate_distance

from idpconfgen import Path


CONF_NUMBER = Queue()

# Checks for clashes then uses default psurgeon function
# Custom clash check modified from count_clashes() in ldrs_helper.py
def check_and_stitch(idr_path, fld_path, output):
    # Custom ranges here can be changed
    lowup = [(49, 134)]
    idr_path = Path(idr_path)
    fragment_struc = Structure(idr_path)
    fragment_struc.build()
    fragment = fragment_struc.data_array
    
    fld_struc = Structure(Path(fld_path))
    fld_struc.build()
    
    fragment_atoms = fragment[:, col_name]
    fragment_seq = fragment[:, col_resSeq].astype(int)
    fragment_coords = fragment[:, cols_coords].astype(float)
    
    parent_coords = fld_struc.data_array[:, cols_coords].astype(float)
    parent_atoms = fld_struc.data_array[:, col_name]
    
    first_r = fragment_seq[0]
    last_r = fragment_seq[-1]
    
    for i, _ in enumerate(fragment_seq):
        try:  # In case the user wants to build less than 3 residues
            next = fragment_seq[i + 1]
        except IndexError:
            continue
        fragment_atoms = fragment_atoms[1:]
        fragment_coords = fragment_coords[1:]
        if next - first_r == 3:
            break
    
    for i, _ in enumerate(fragment_seq):
        j = len(fragment_seq) - 1 - i
        try:  # In case the user wants to build less than 3 residues
            prev = fragment_seq[j - 1]
        except IndexError:
            continue
        fragment_atoms = fragment_atoms[:-1]
        fragment_coords = fragment_coords[:-1]
        if last_r - prev == 3:
            break
    
    num_clashes = 0
    for i, atom1 in enumerate(parent_atoms):
        a1 = atom1[0]
        for j, atom2 in enumerate(fragment_atoms):
            a2 = atom2[0]
            # calculate distance between atoms
            distance = calculate_distance(parent_coords[i], fragment_coords[j])
            
            # get vdW radii for each atom
            vdw_radius1 = vdW_radii_tsai_1999[a1]
            vdw_radius2 = vdW_radii_tsai_1999[a2]
            
            # Check if a steric clash is detected by comparing
            # distance between atoms to the sum of their vdW radii
            if num_clashes > 10:
                return
            if distance < vdw_radius1 + vdw_radius2 + 0.4:
                num_clashes += 1

    new_struc_arr = psurgeon([[idr_path]], fld_struc, ["Linker-IDR"], lowup)
    
    new_struc = structure_to_pdb(new_struc_arr)
    output_p = output + f"conformer_{CONF_NUMBER.get()}.pdb"
    with open(output_p, 'w') as f:
        for line in new_struc:
            f.write(line + "\n")

        
# Change variables here
parent_path = "/PATH/TO/PARENT/FOLDER"
ncores = 32
nconfs = 10000
rs = 0
random.seed(rs)

output_path = f"{parent_path}/A9_conformers_bb/"
idr_paths = glob.glob(f"{parent_path}/A9_results_bb/*.pdb")
fld_path = f"{parent_path}/7CH1_STAS.pdb"

for i in range(1, nconfs + 1):
    CONF_NUMBER.put(i)

execute = partial(
    check_and_stitch,
    fld_path=fld_path,
    output=output_path,
    )
execute_pool = pool_function(execute, idr_paths, ncores=ncores)
for _ in execute_pool:
    pass
```
