## Supplementary figures and images for "Local Disordered Region Sampling (LDRS) for Ensemble Modeling of Proteins with Experimentally Undetermined or Low Confidence Prediction Segments"

### 2MX4_template.png

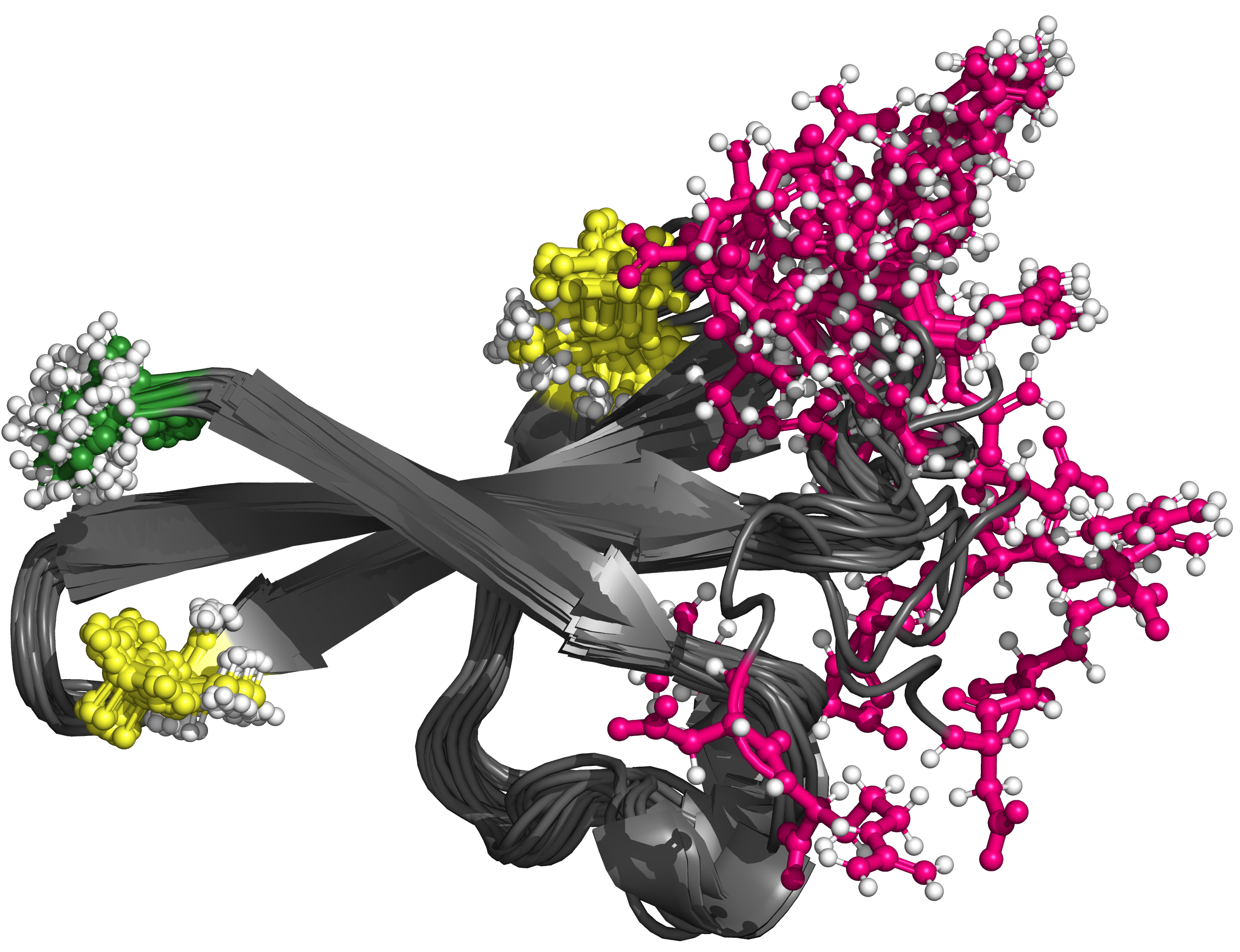

### 4E_BIDR_fracSS.png

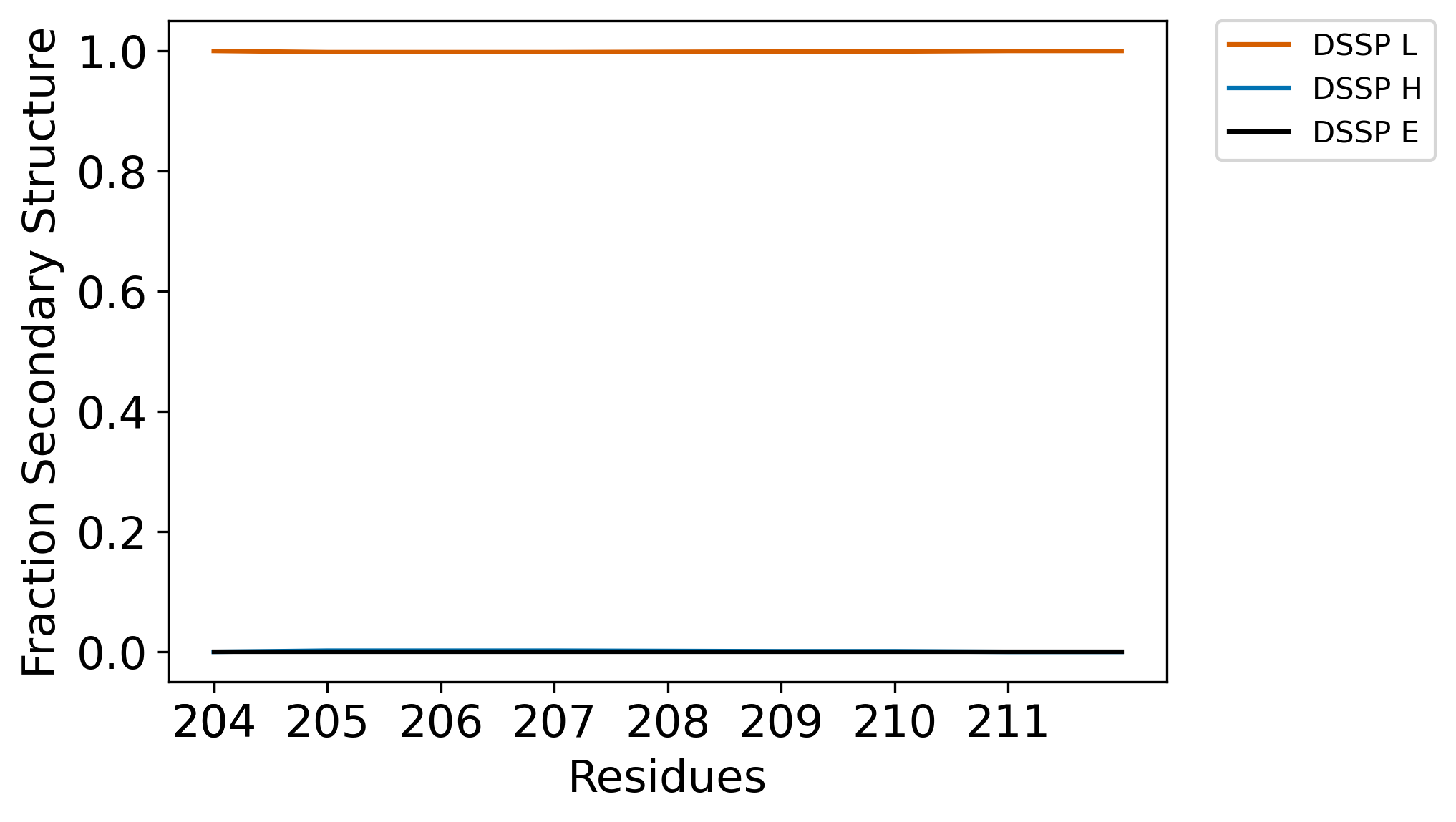

### 4E_BIDR_ramaSS.png

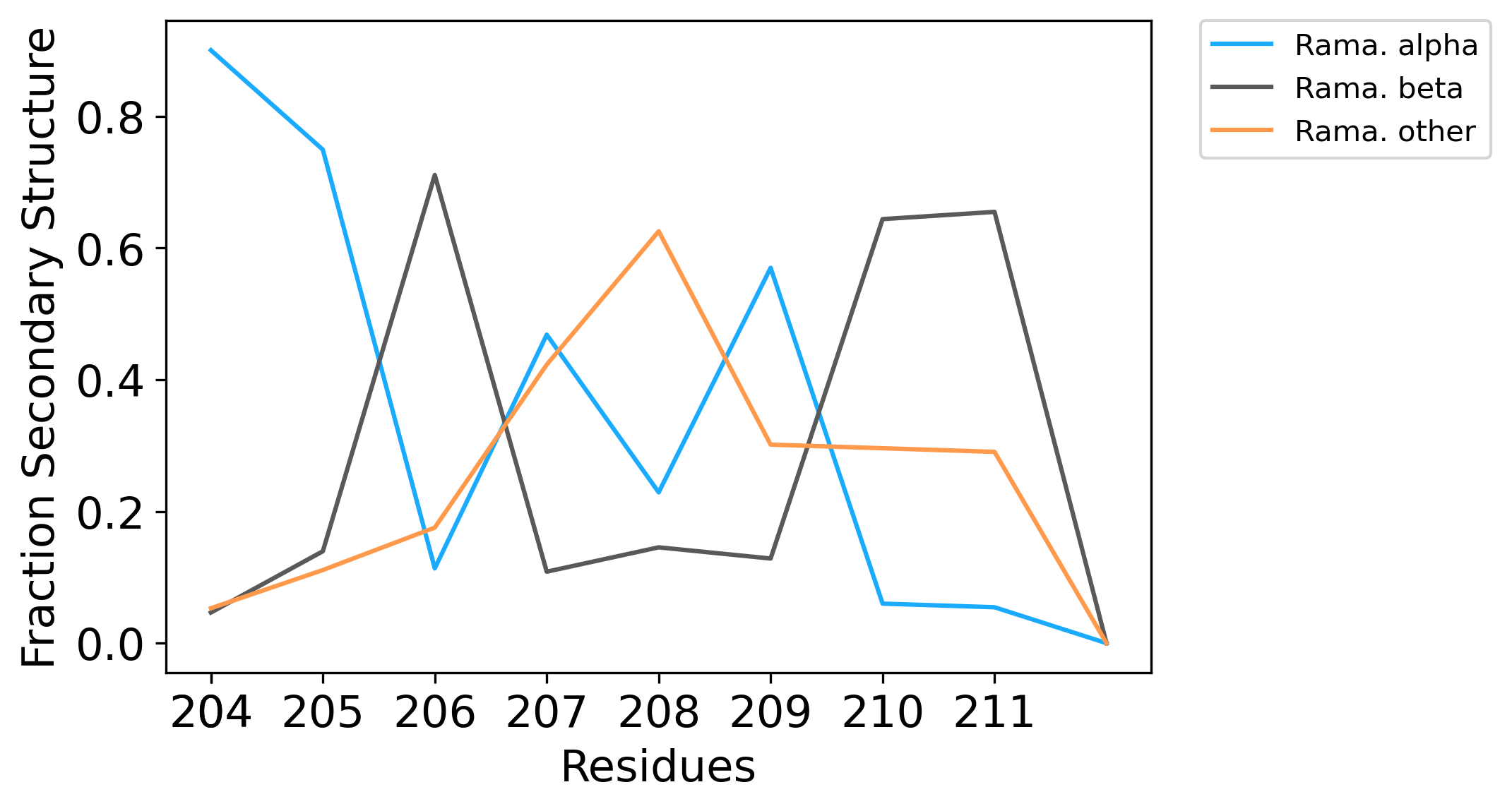

### 4E_NIDR_fracSS.png

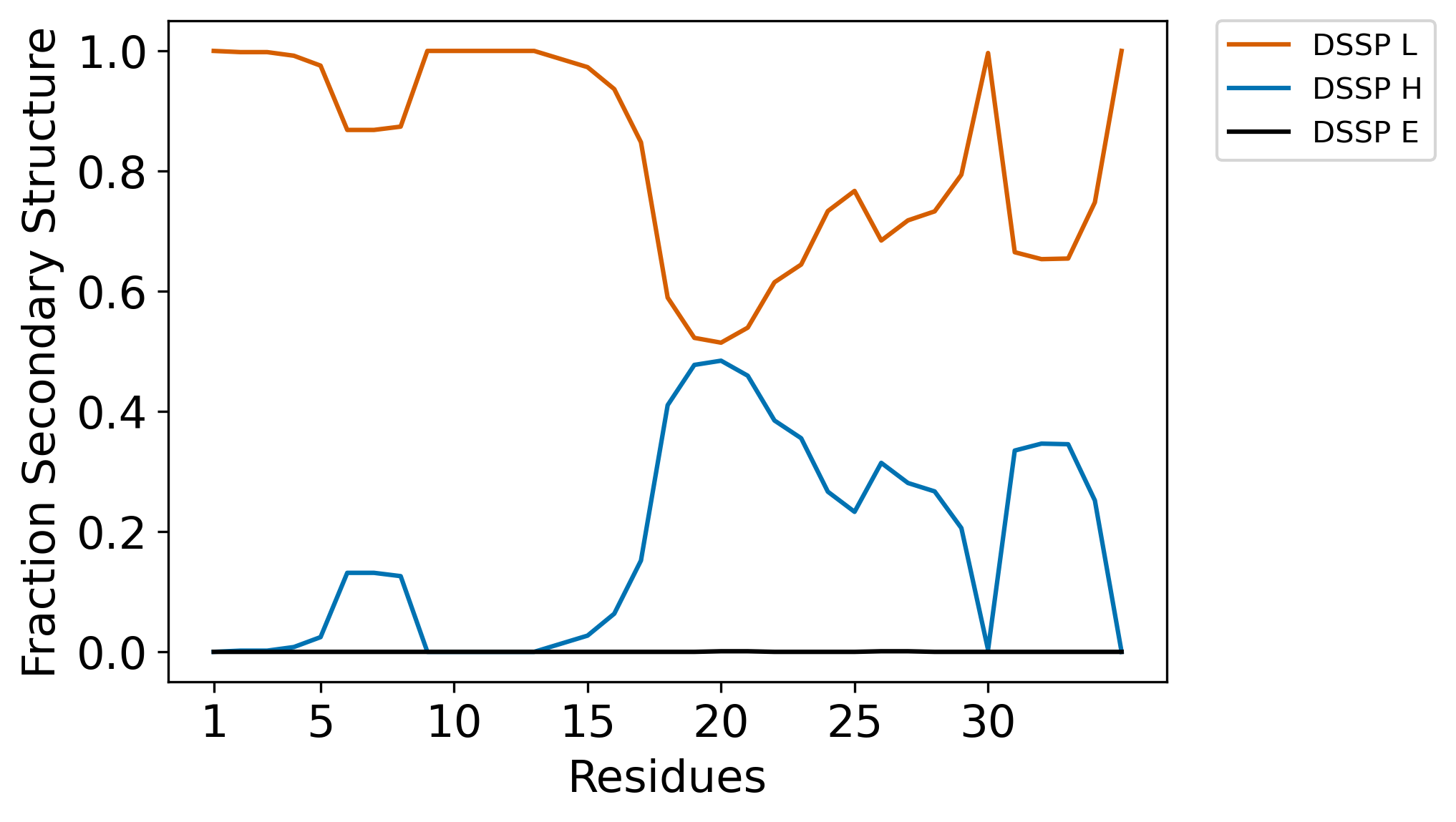

### 4E_NIDR_ramaSS.png

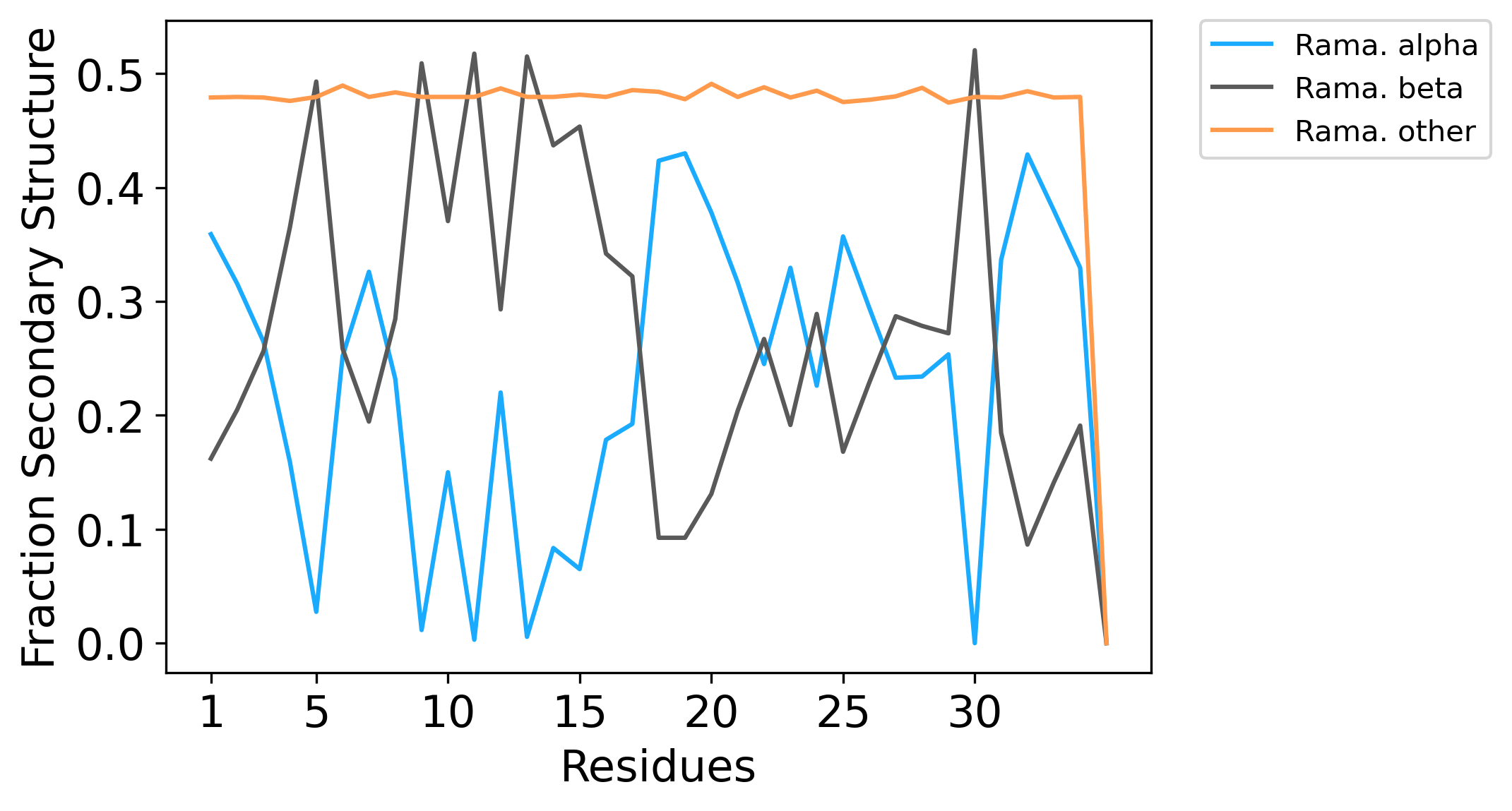

### 4E_rama_plot.png

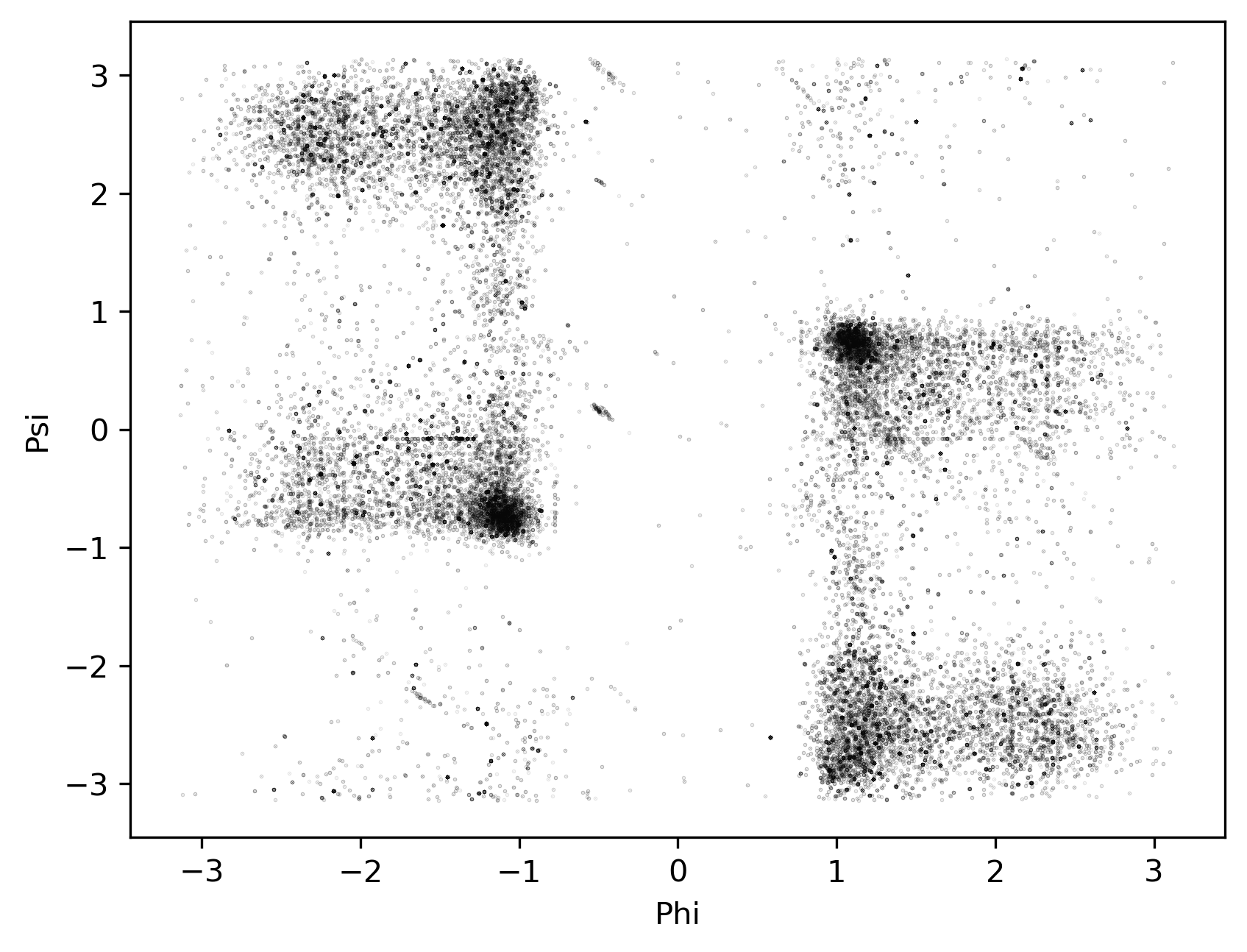

### 5p4EBP2_fracSS.png

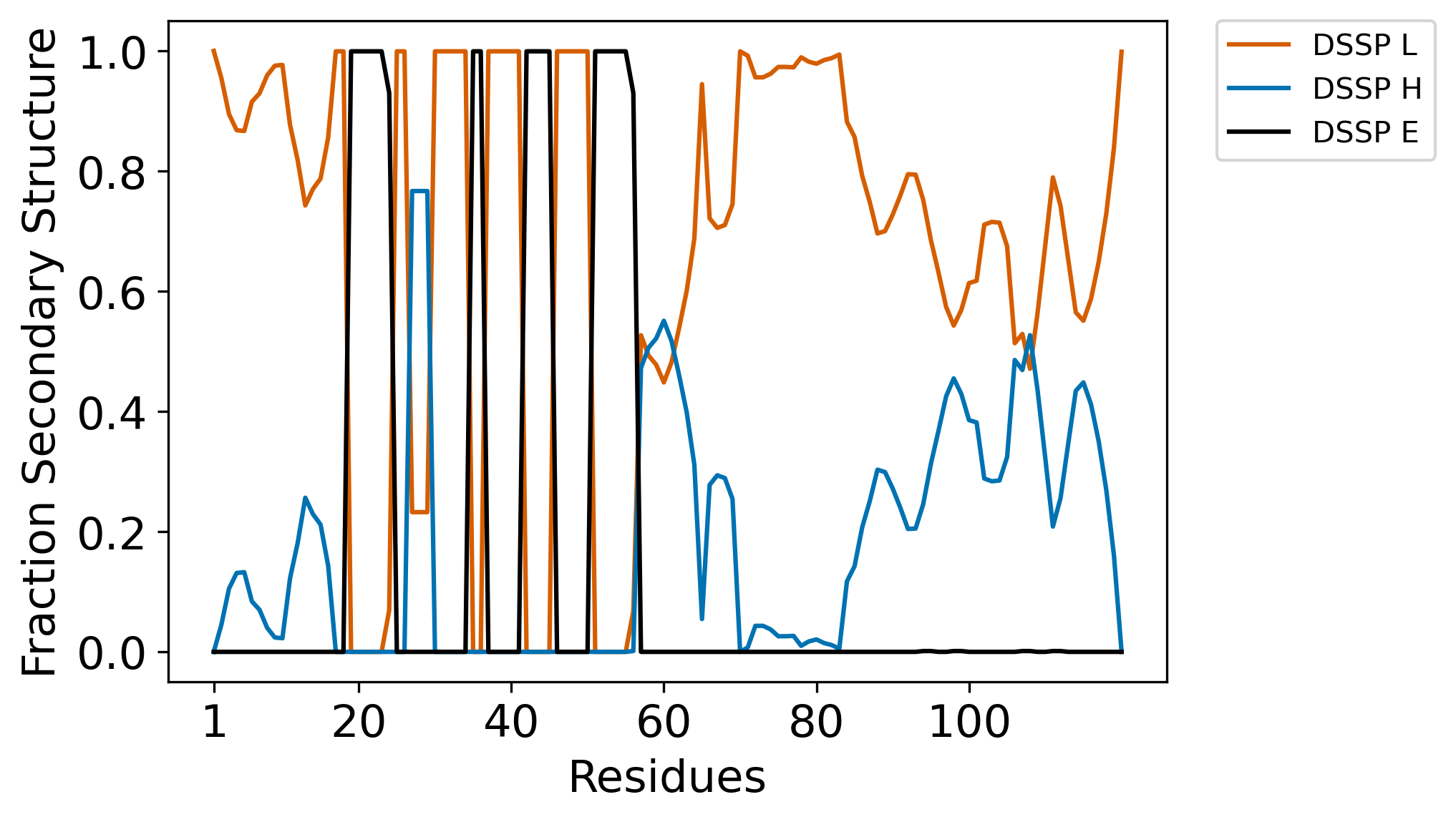

### 5p4EBP2_fracSS_long.png

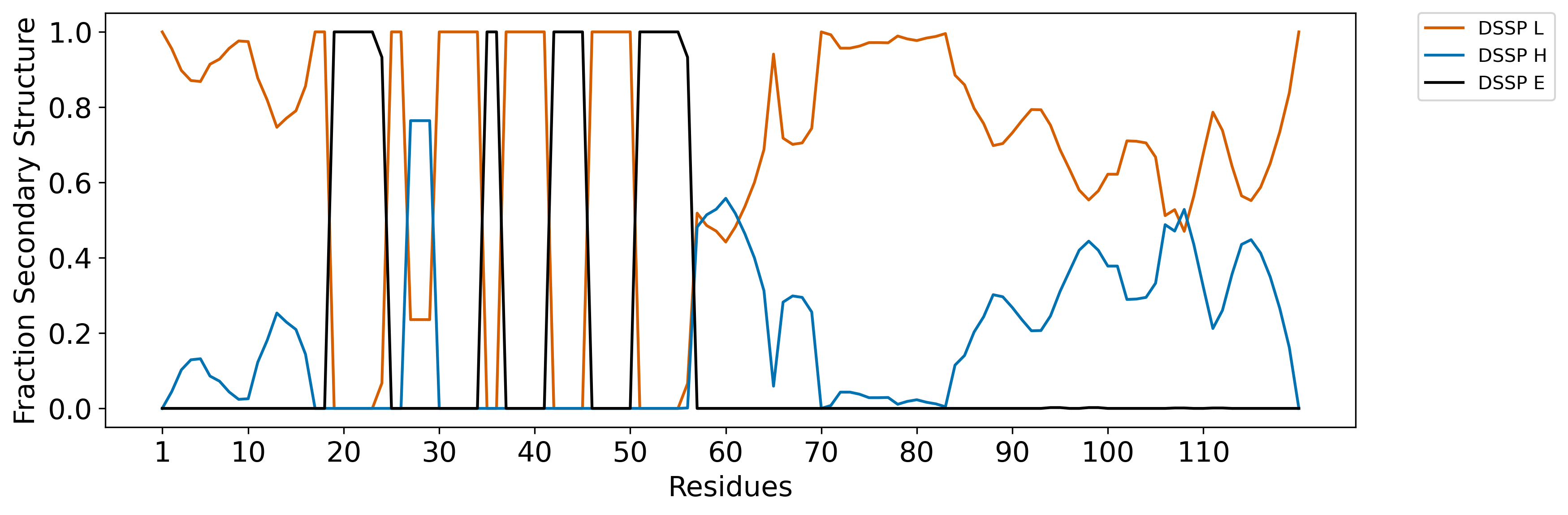

### 5p4EBP2_n1.png

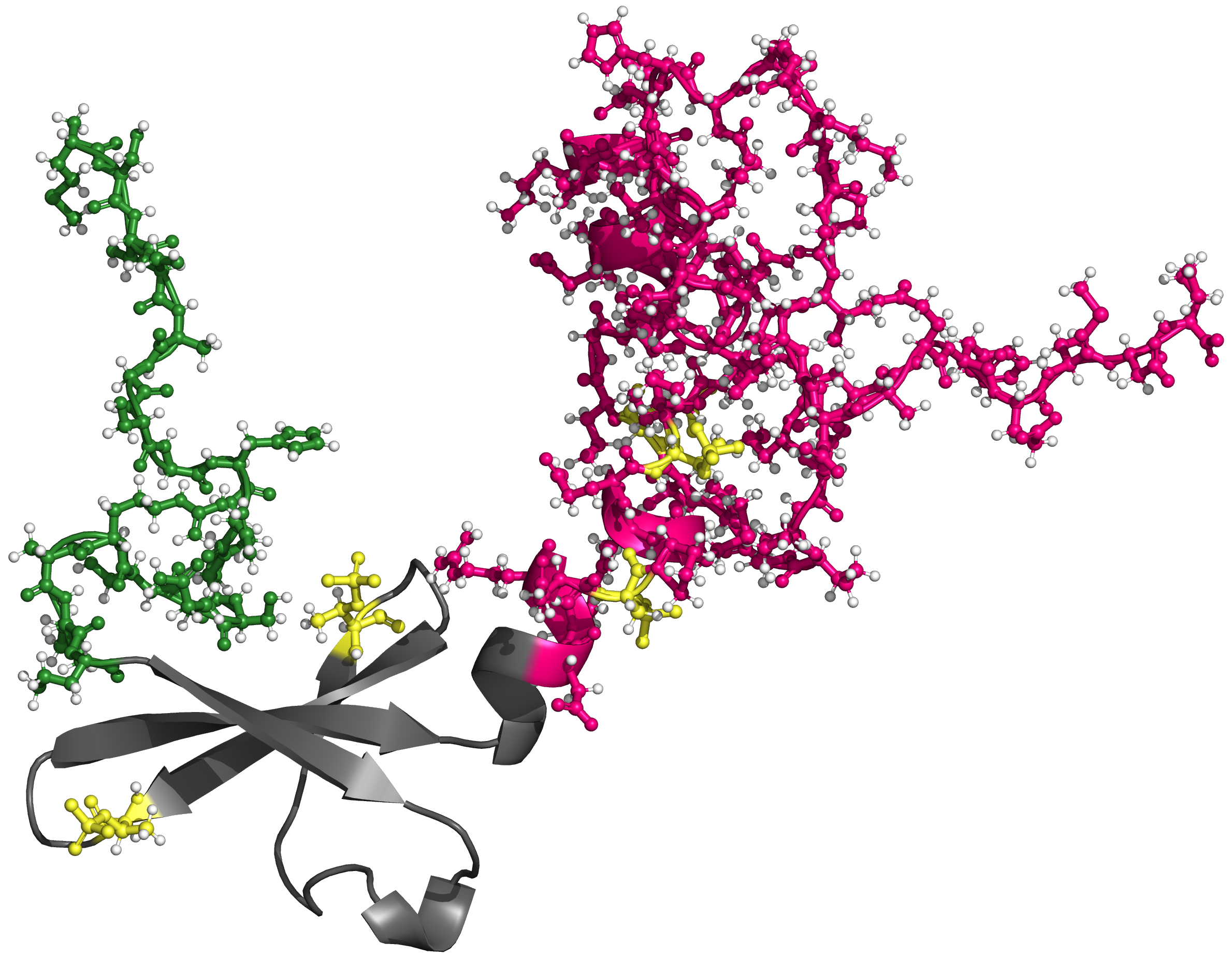

### 5p4EBP2_n100.png

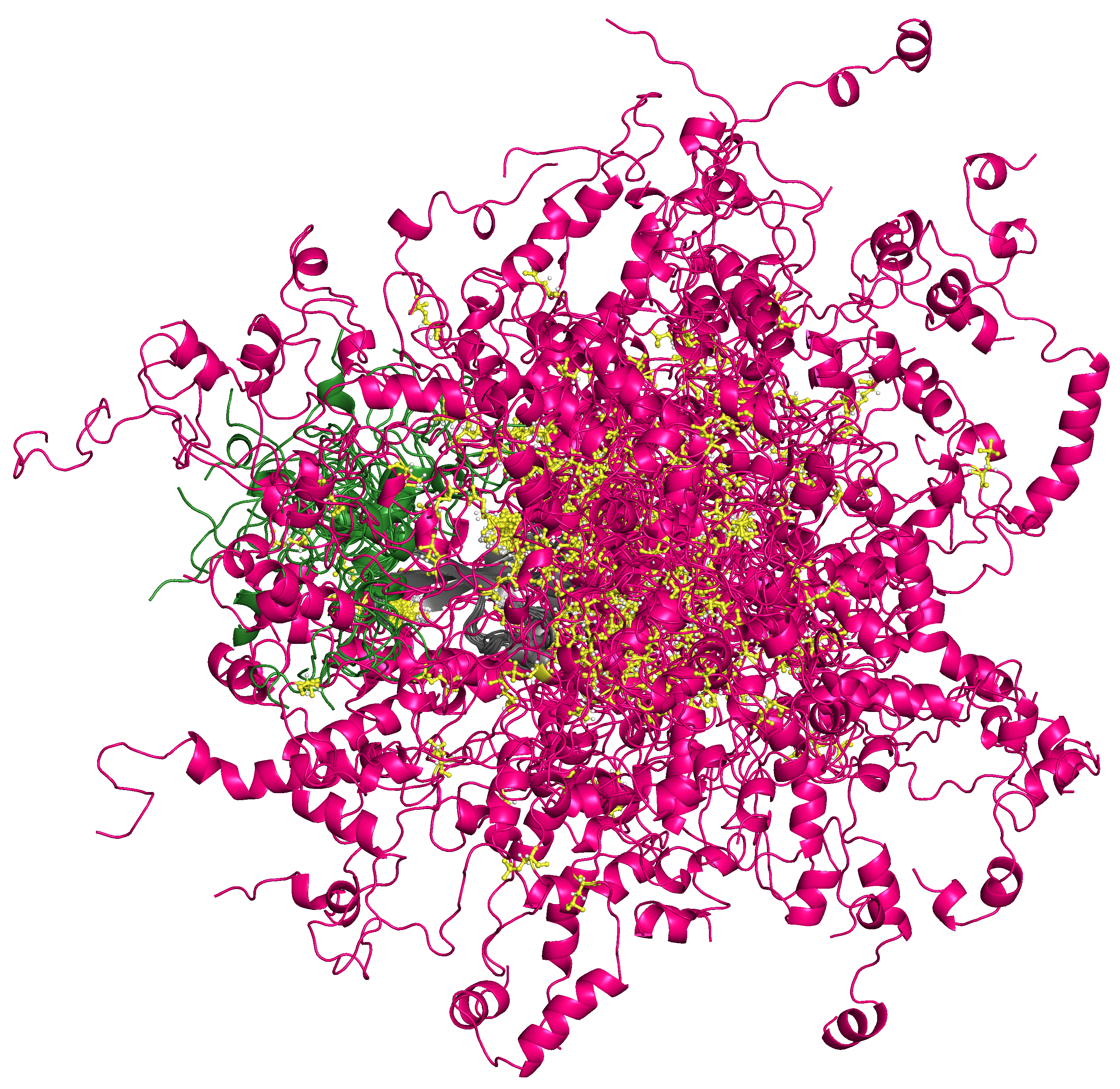

### 5p4EBP2_rama_plot.png

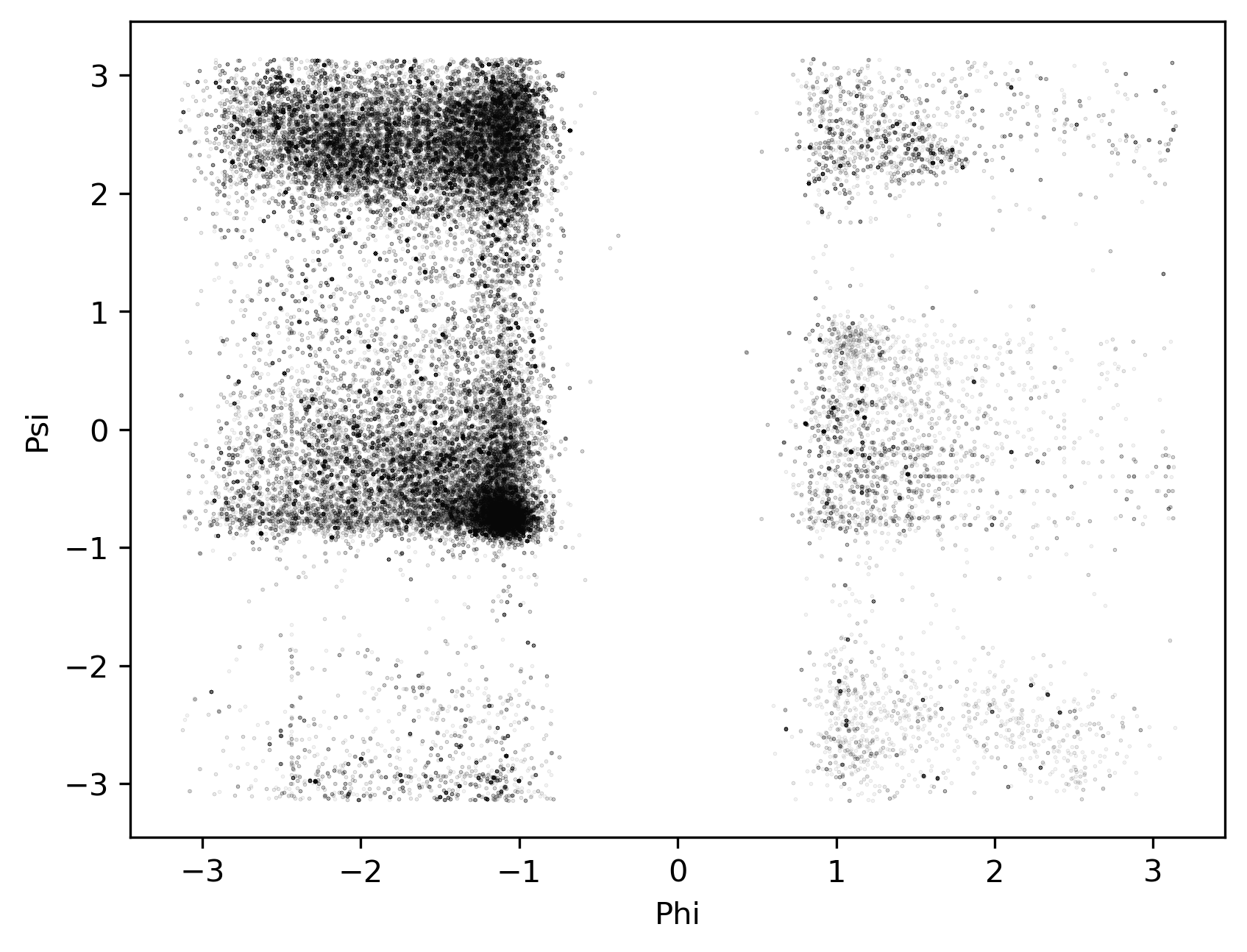

### 5p4EBP2_ramaSS.png

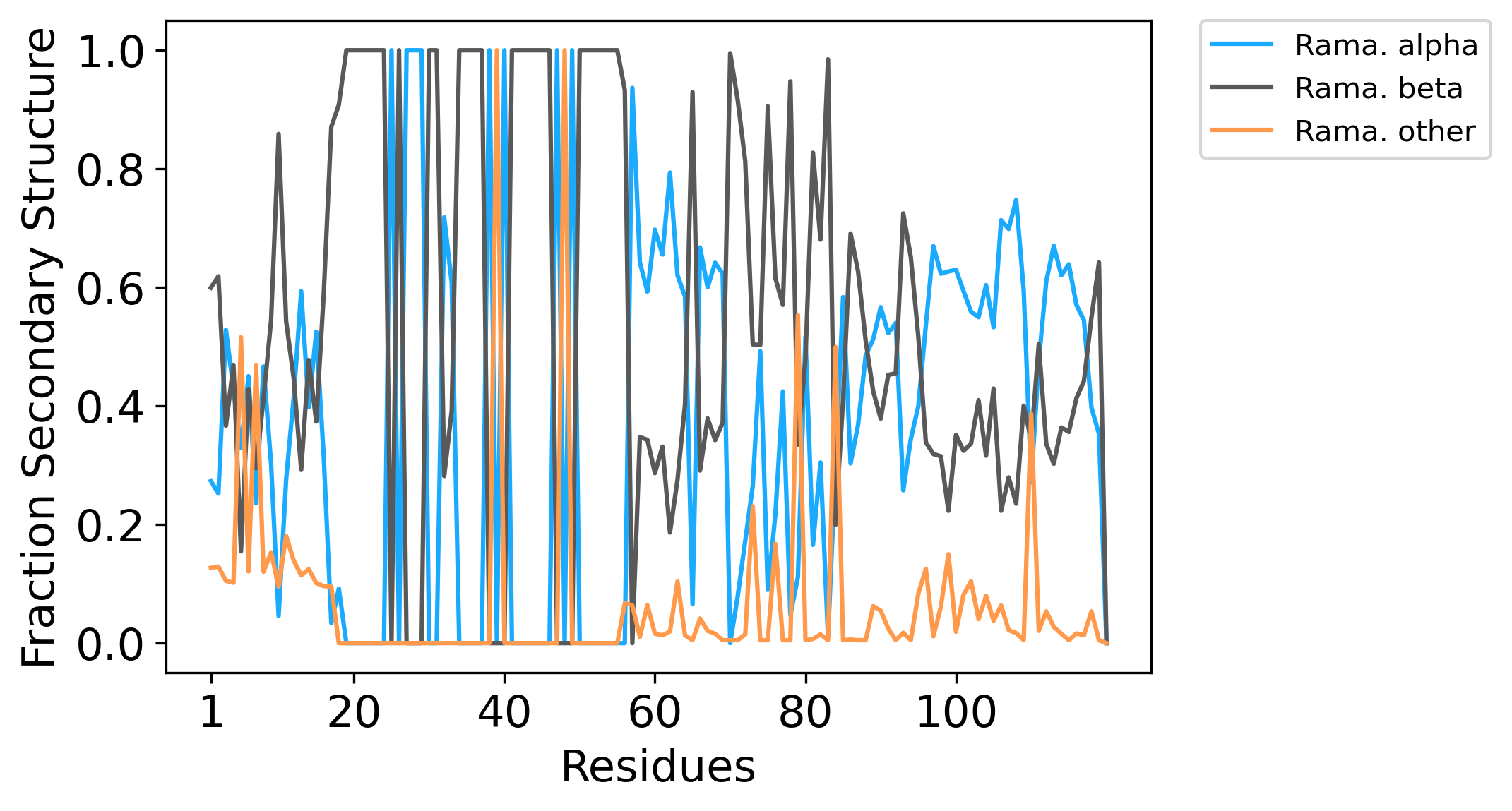

### 5p4EBP2_ramaSS_long.png

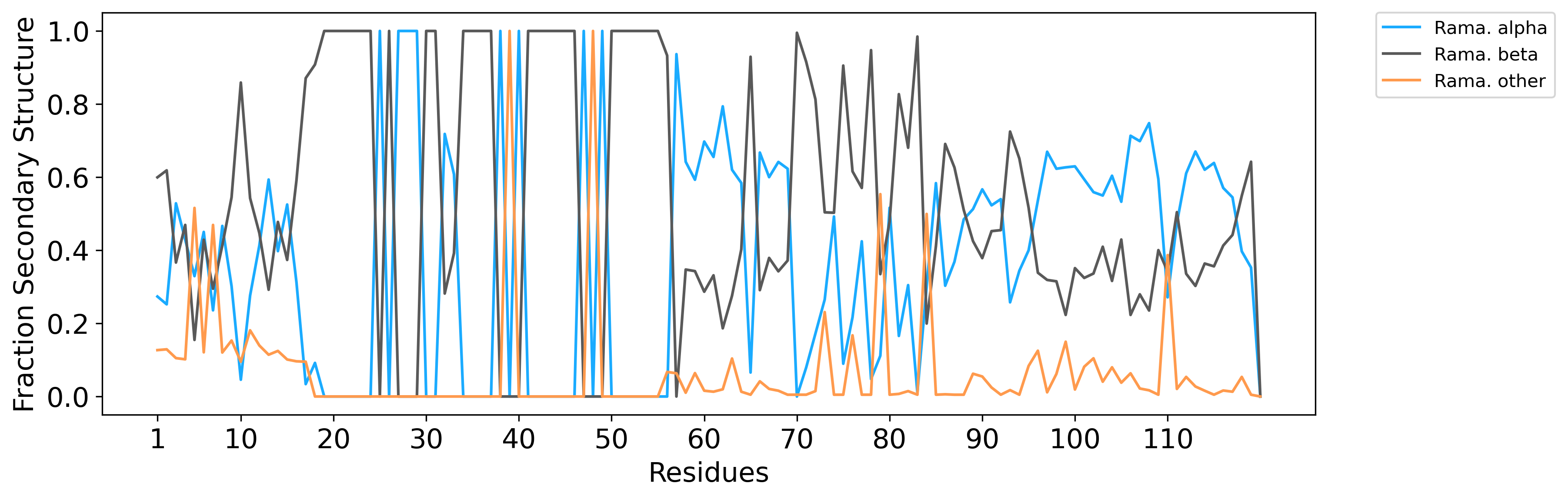

### A2ADR_AF2_dsspSS.png

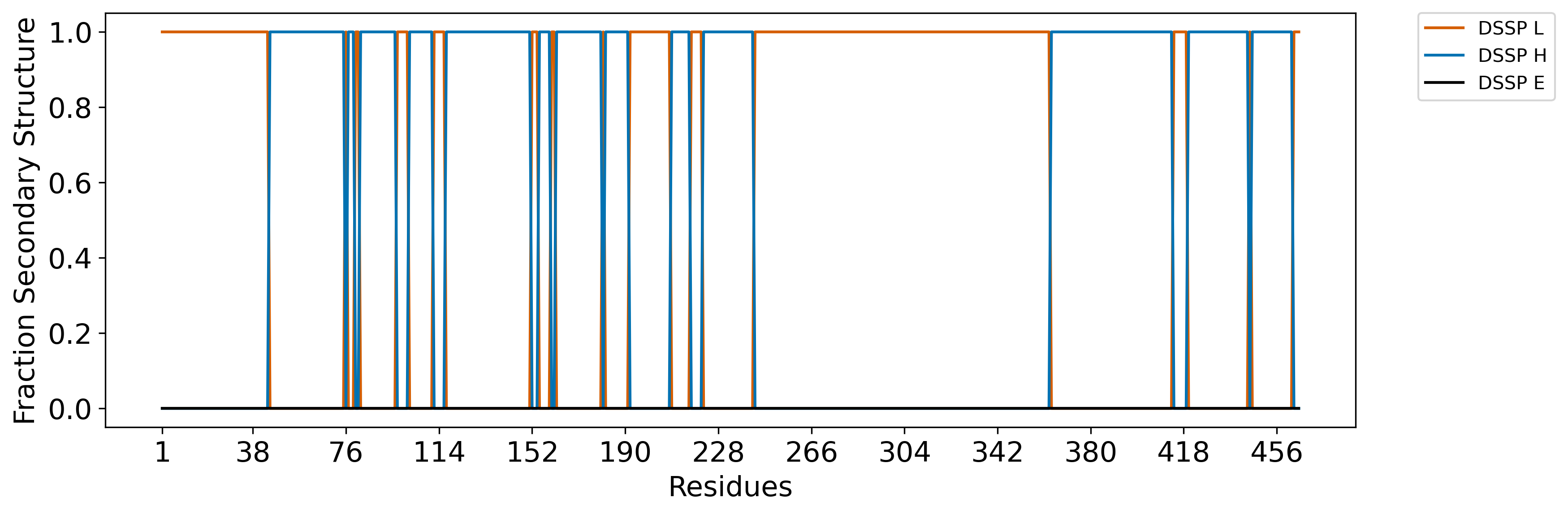

### A2ADR_AF2_ramaSS.png

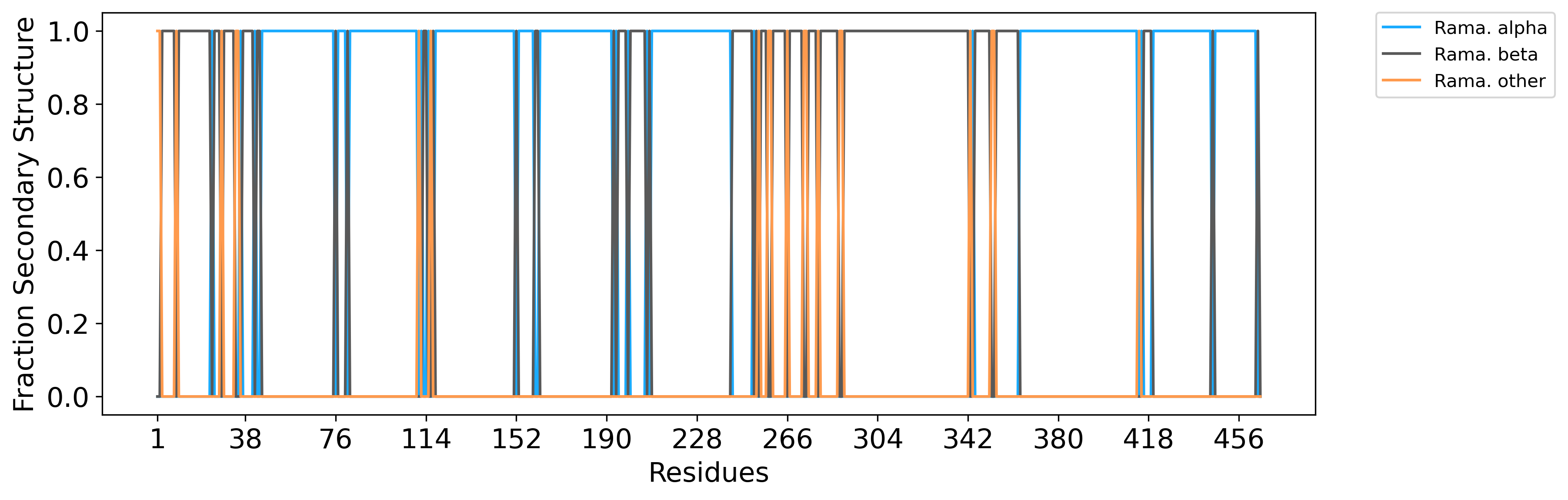

### A2ADR_CIDR_distances.png

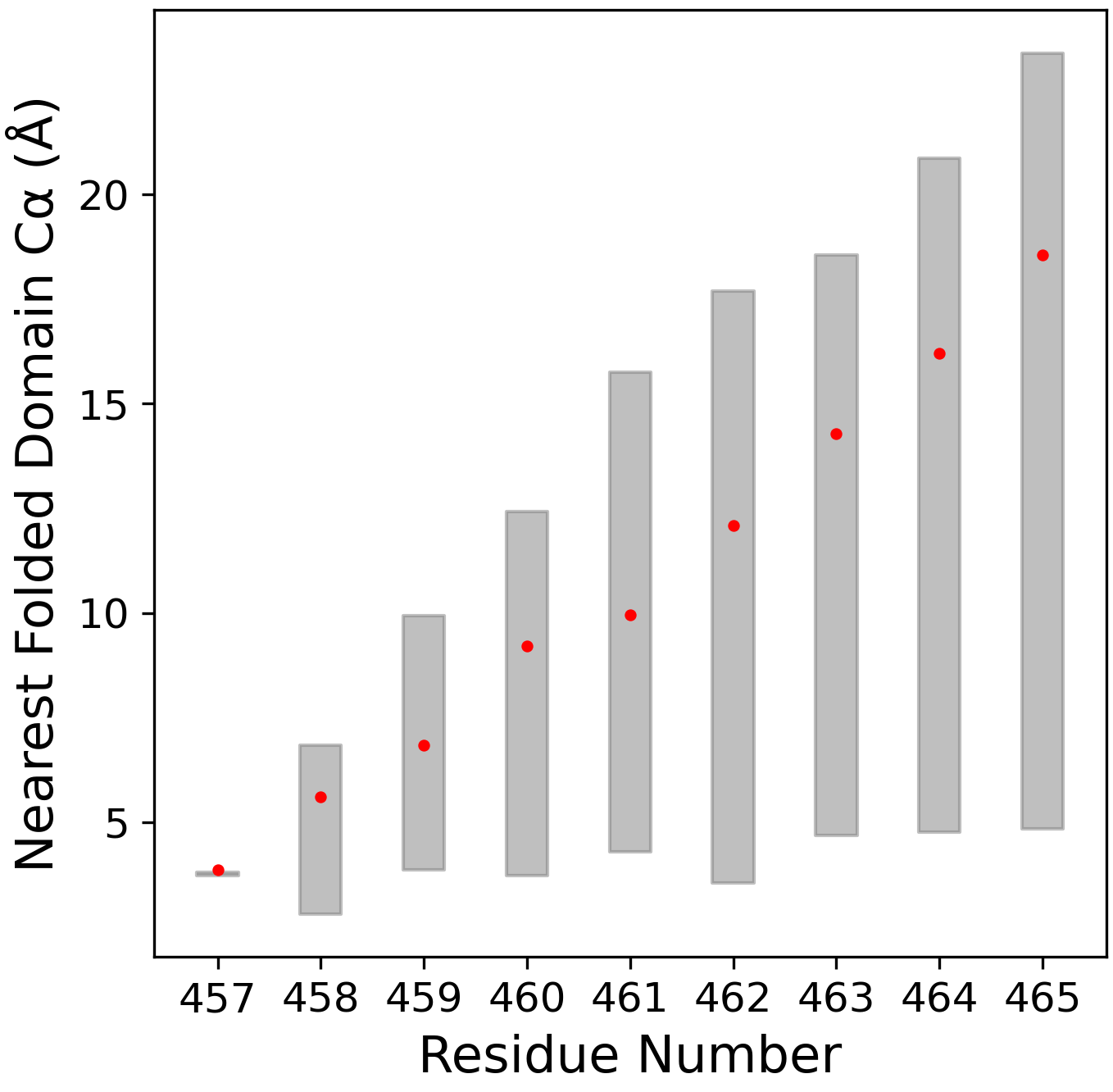

### A2ADR_fracSS.png

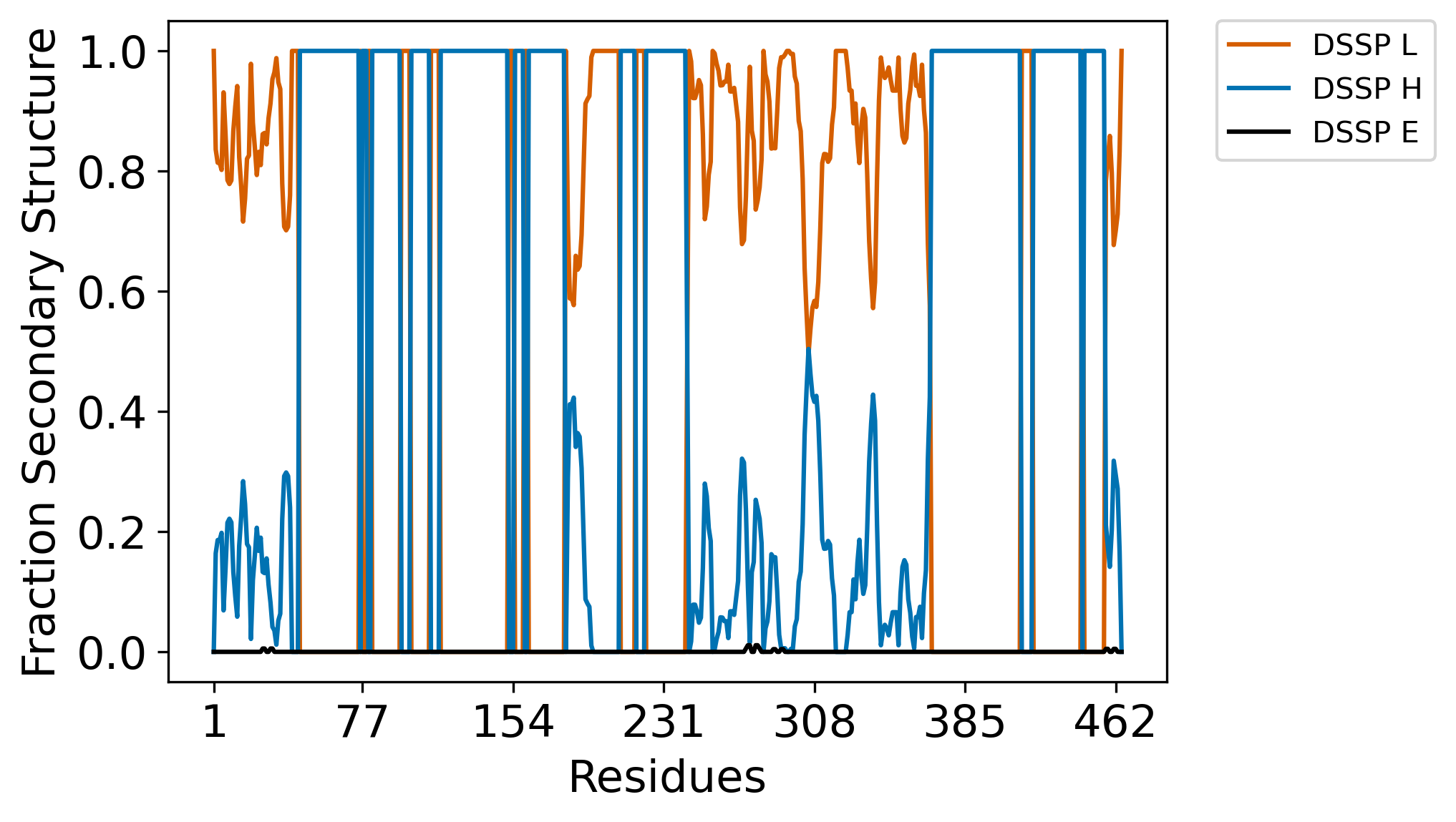

### A2ADR_fracSS_long.png

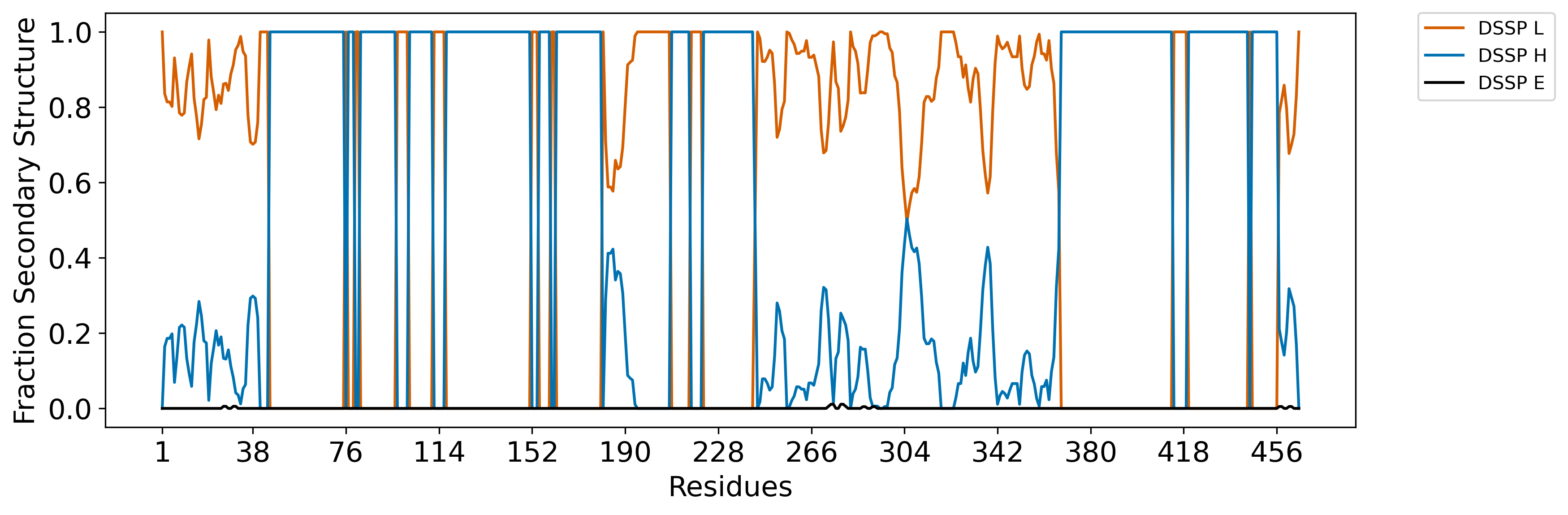

### A2ADR_LIDR1_distances.png

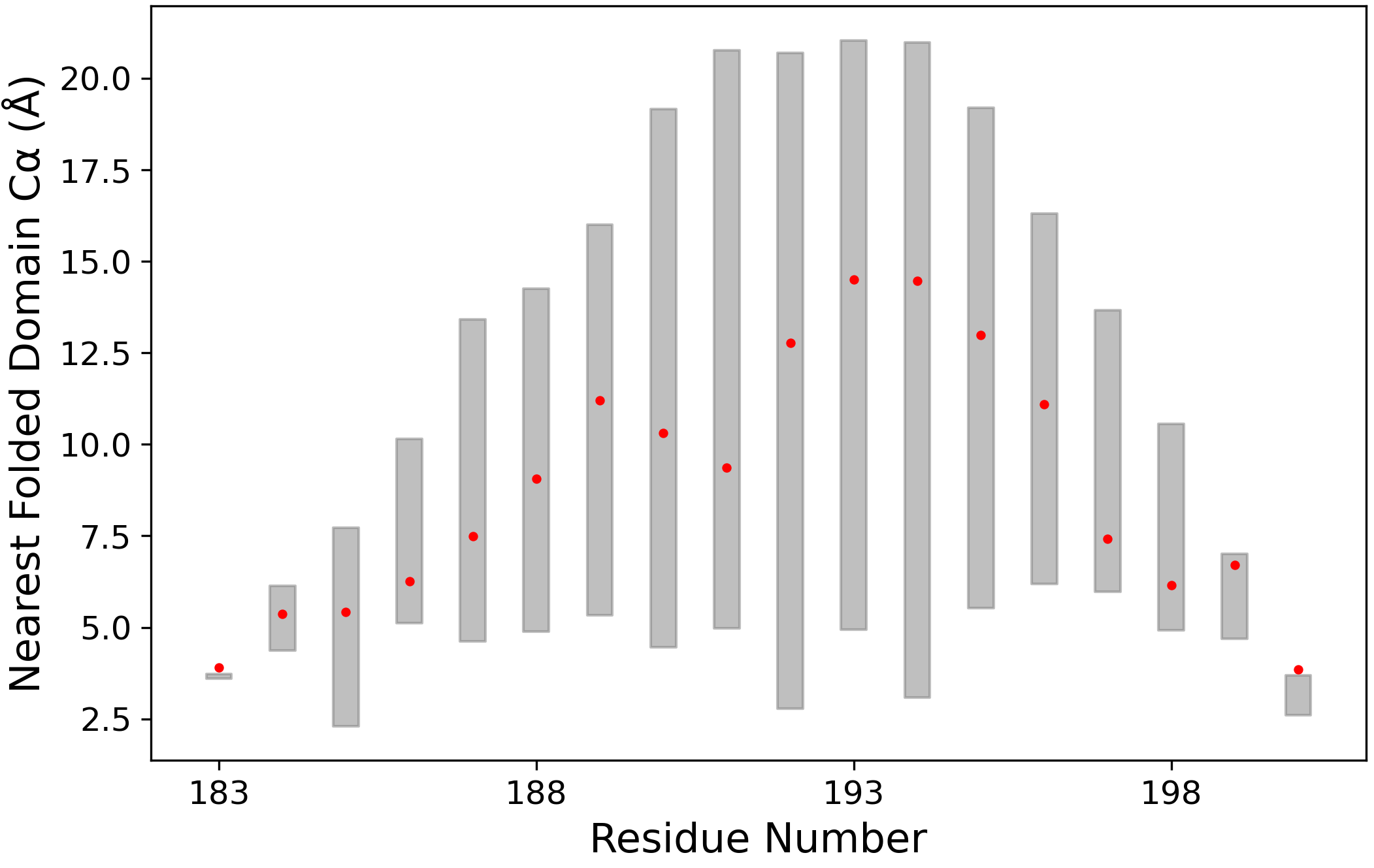

### A2ADR_LIDR2_distances.png

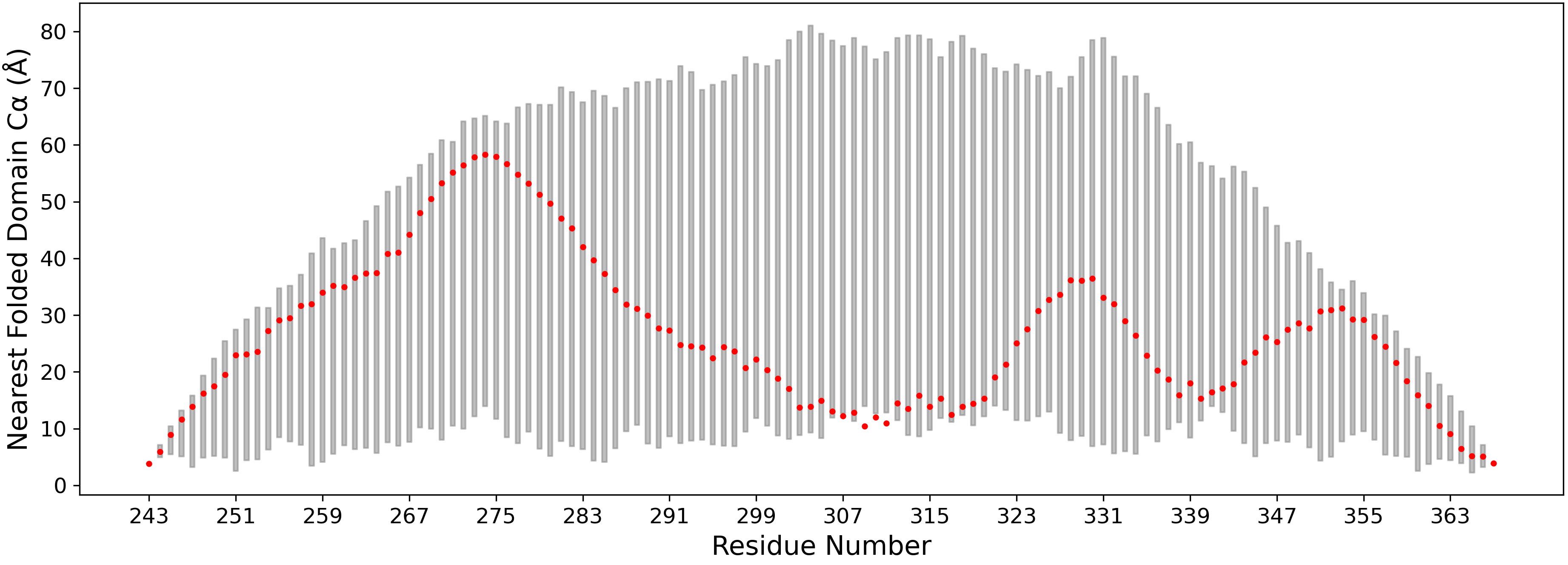

### A2ADR_n100.png

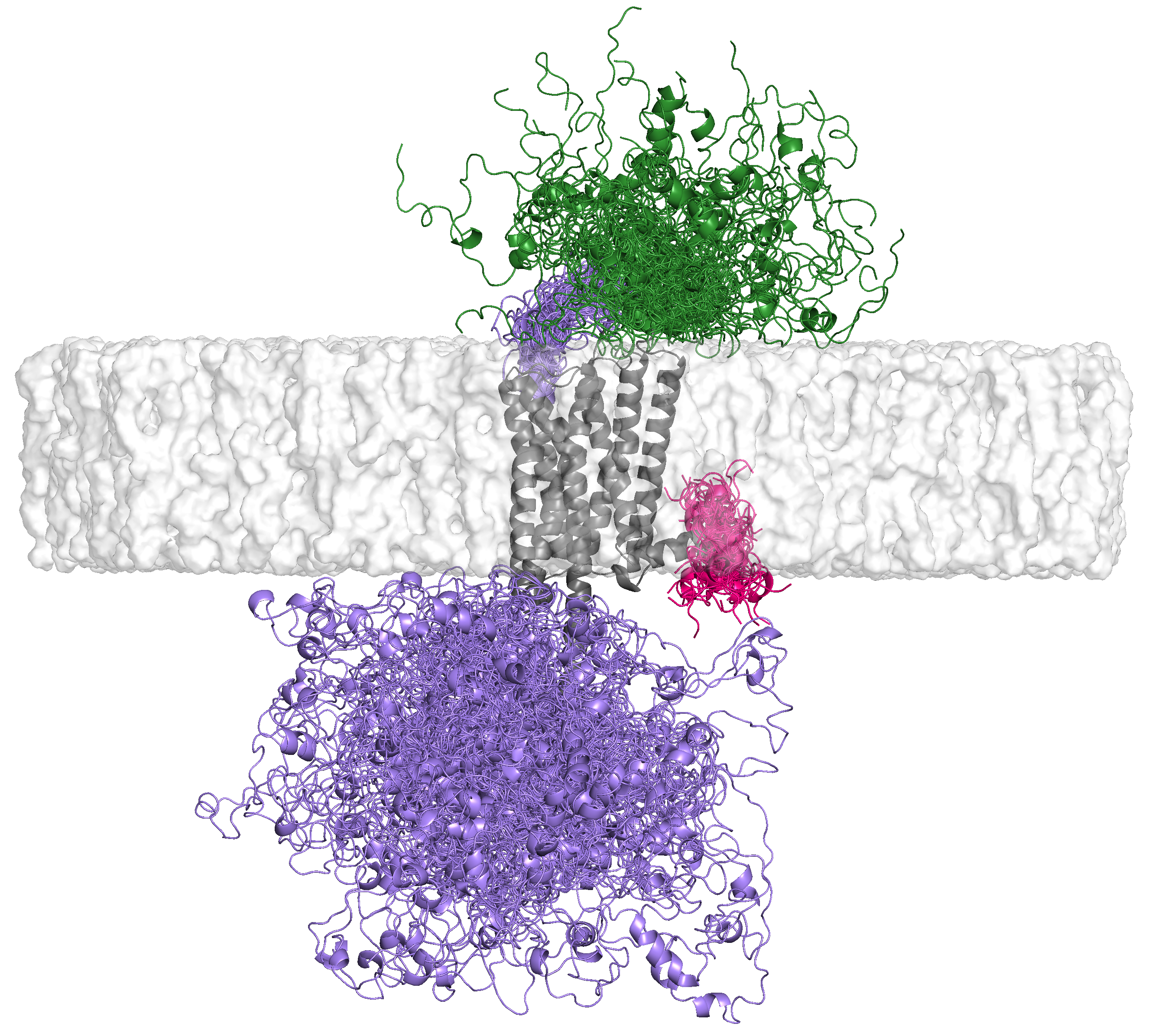

### A2ADR_NIDR_distances.png

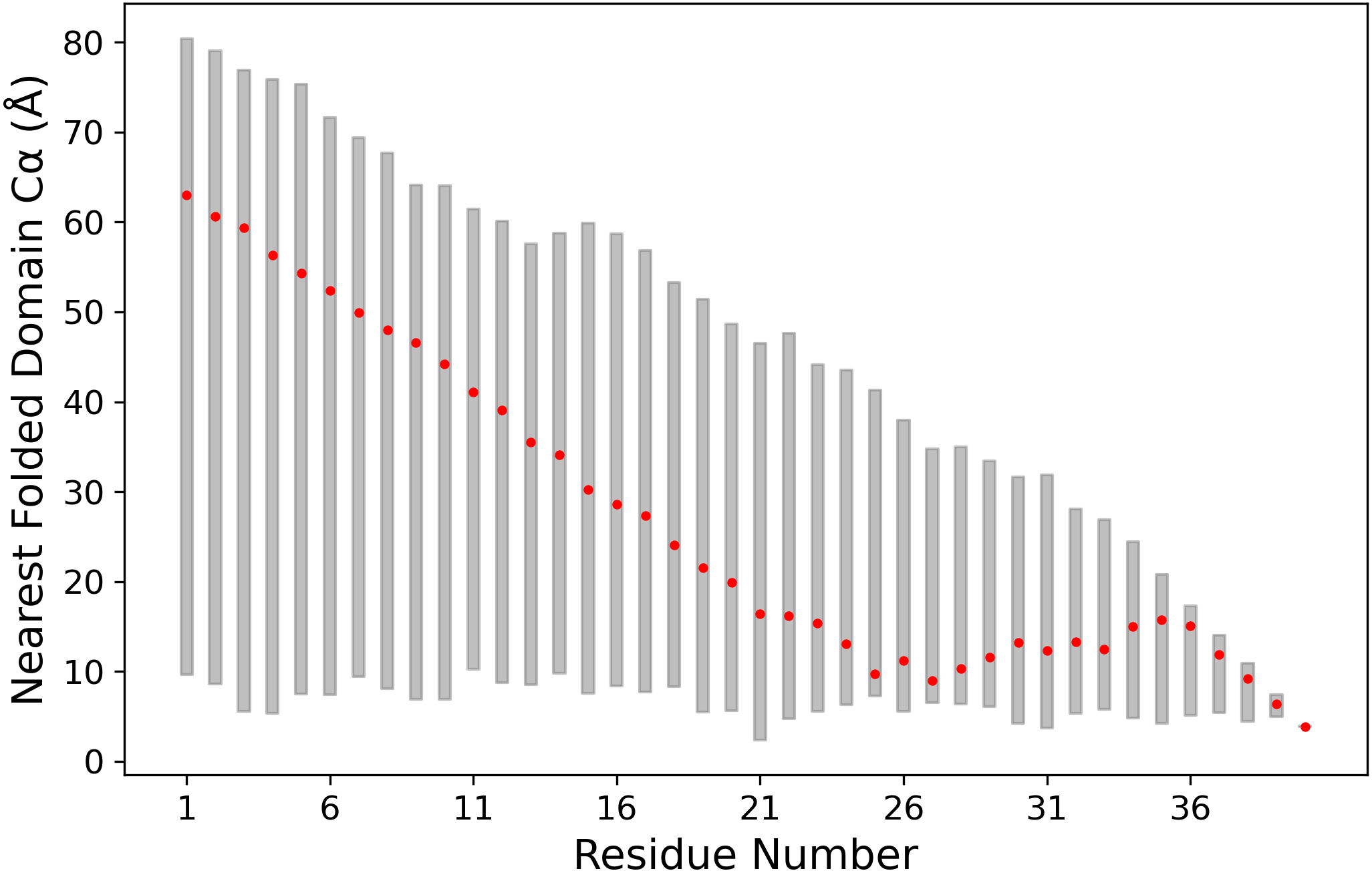

### A2ADR_rama_plot.png

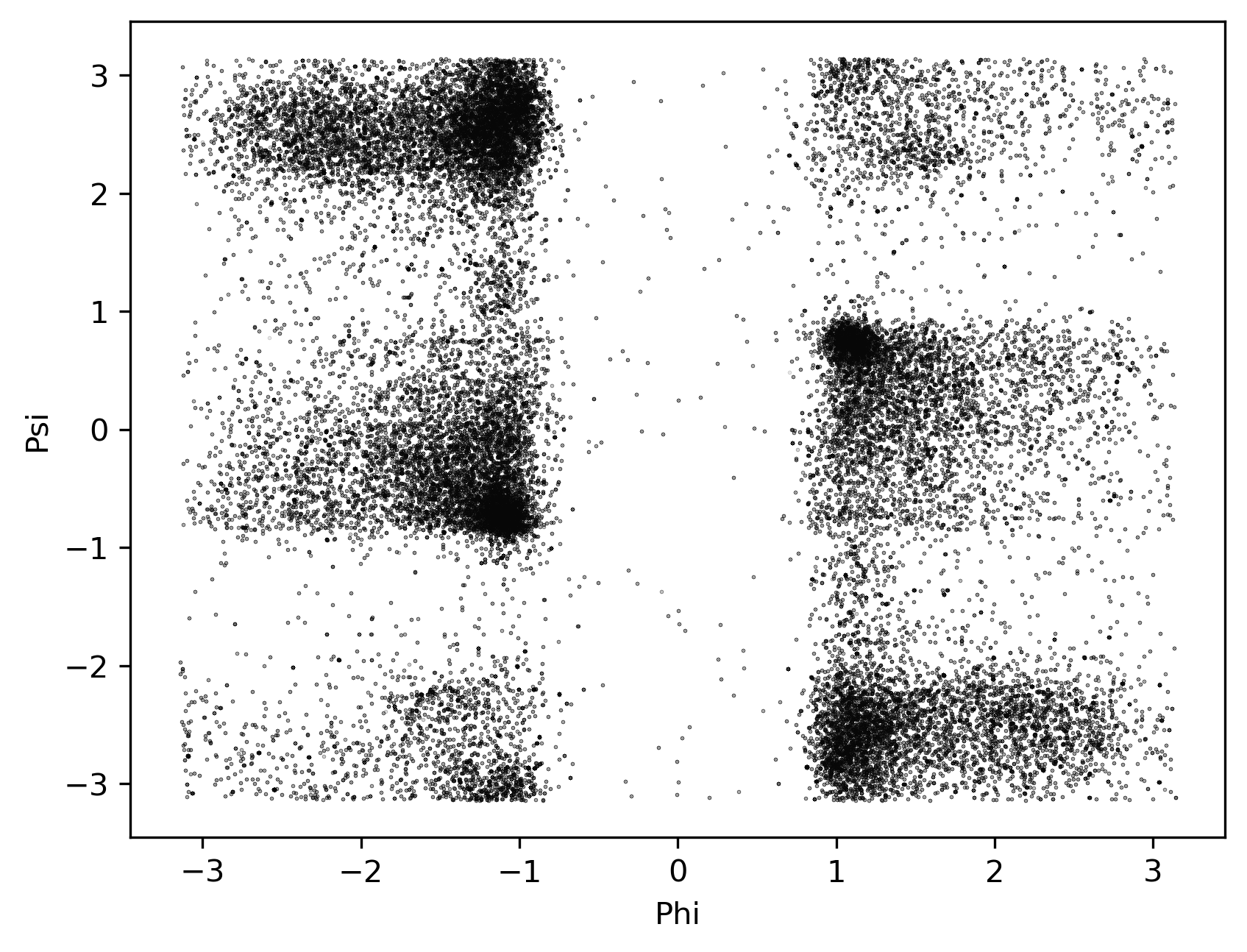

### A2ADR_ramaSS.png

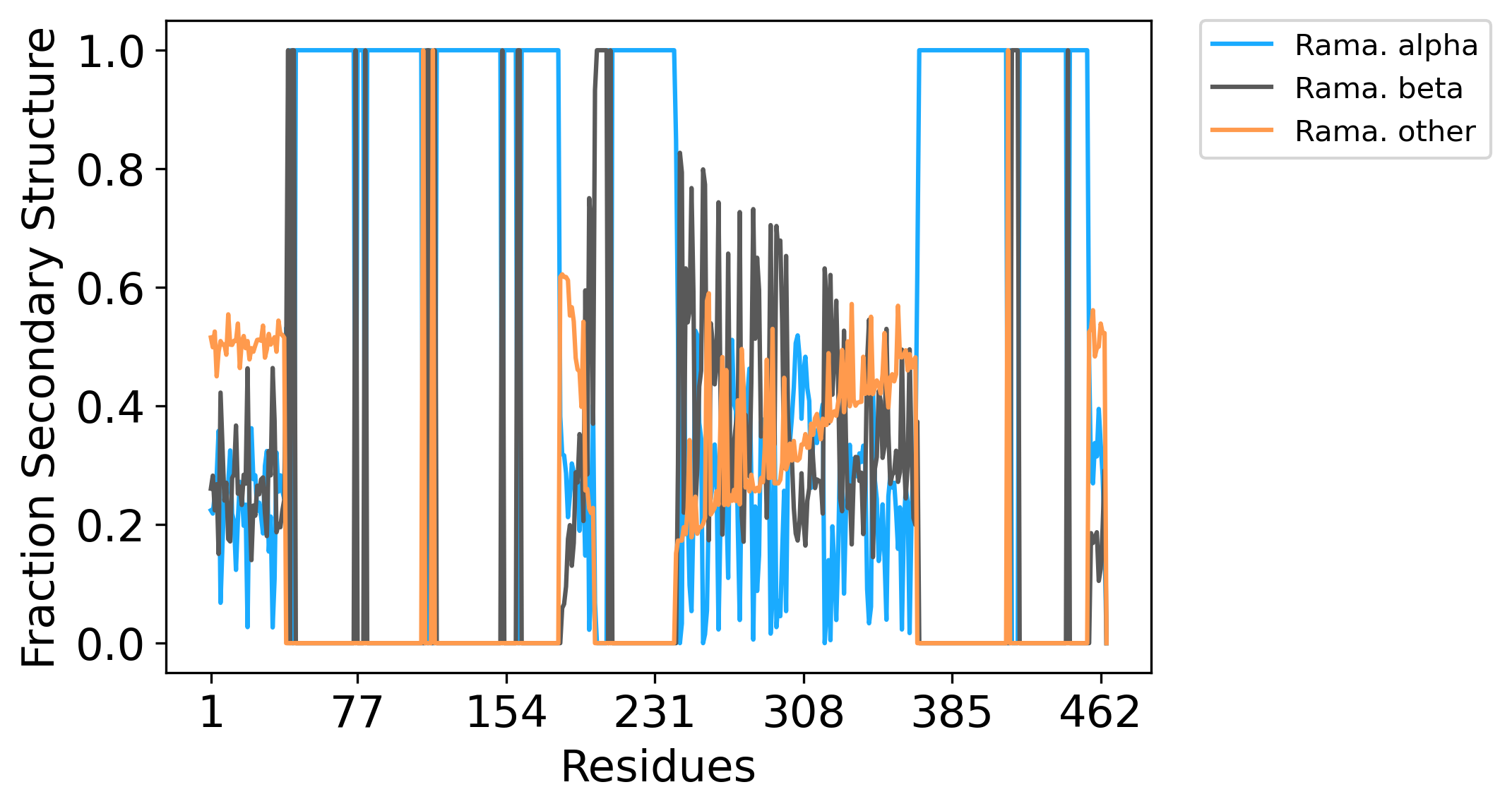

### A2ADR_ramaSS_long.png

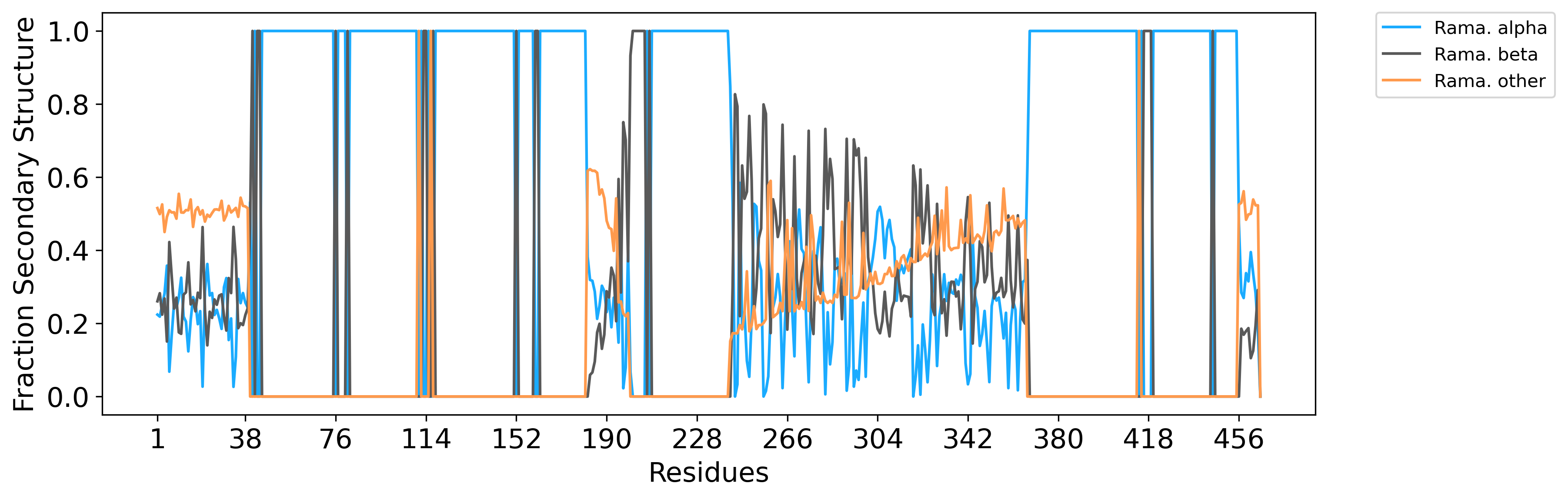

### A9_fracSS.png

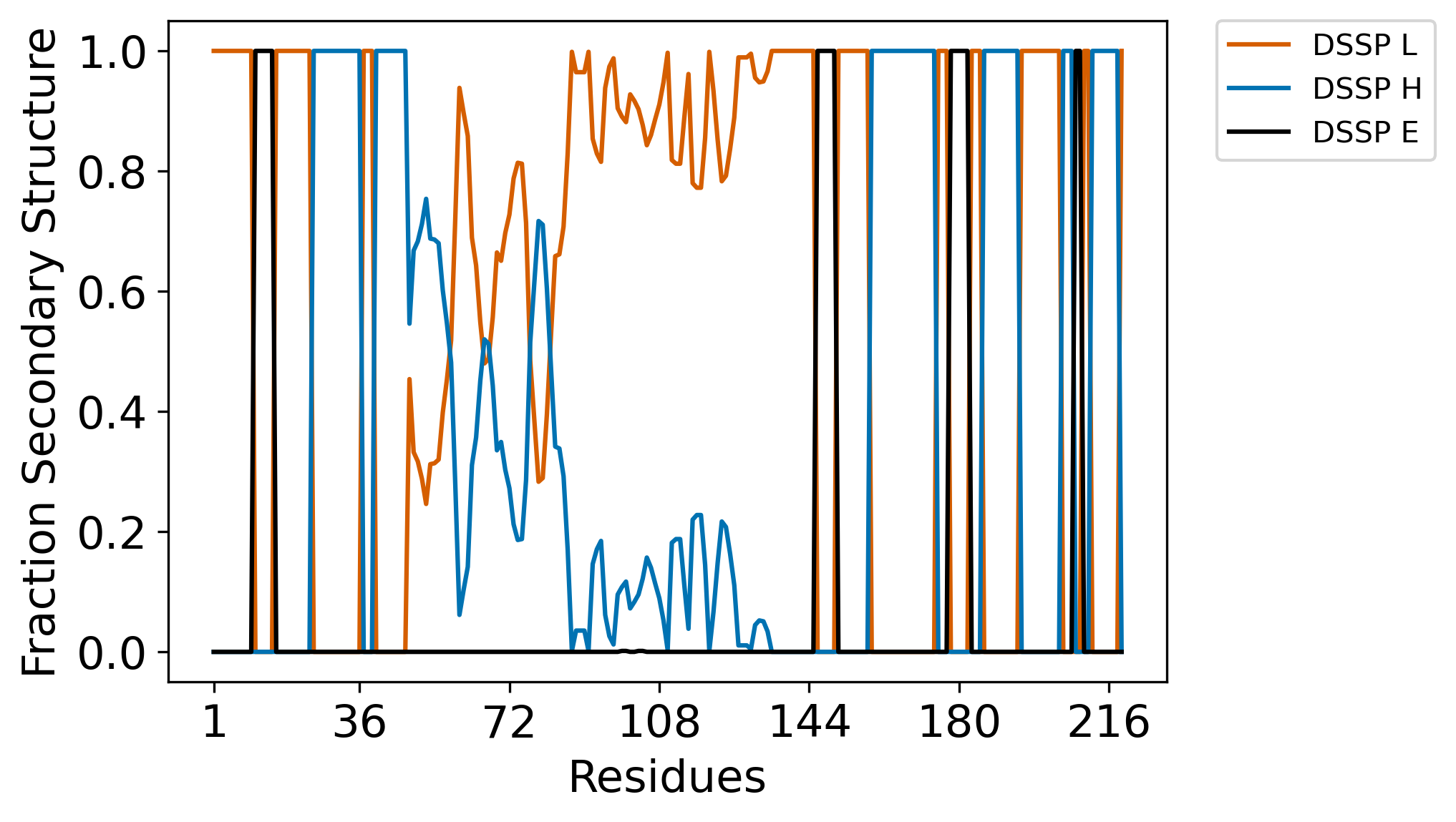

### A9_fracSS_long.png

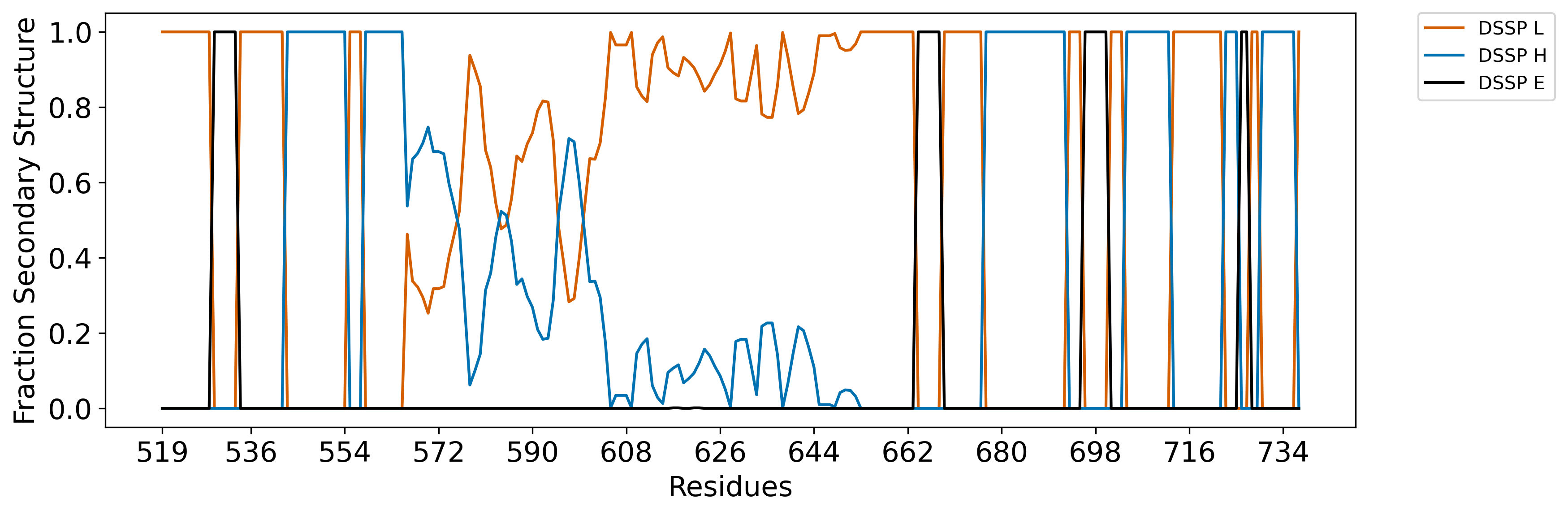

### A9_n1.png

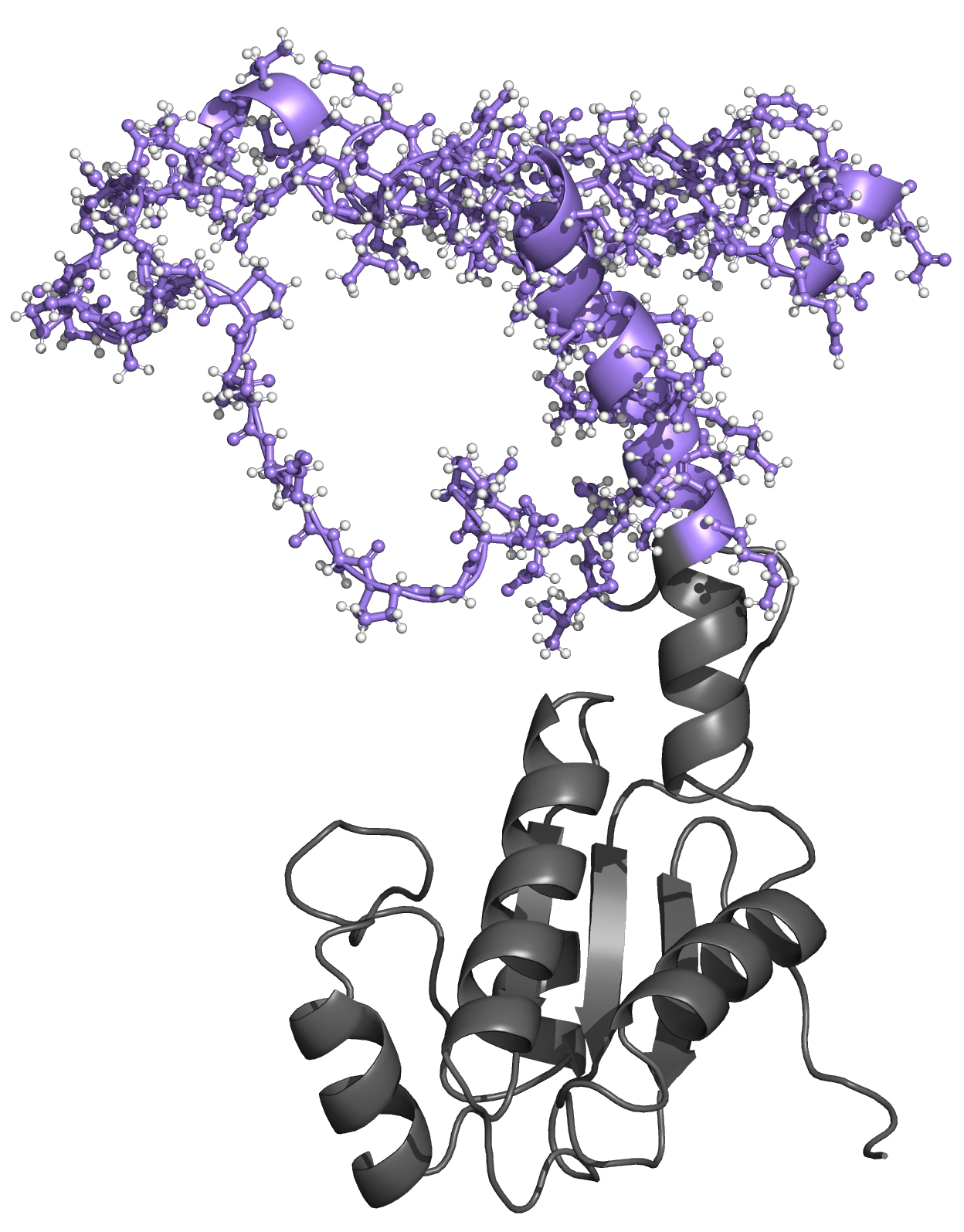

### A9_n100.png

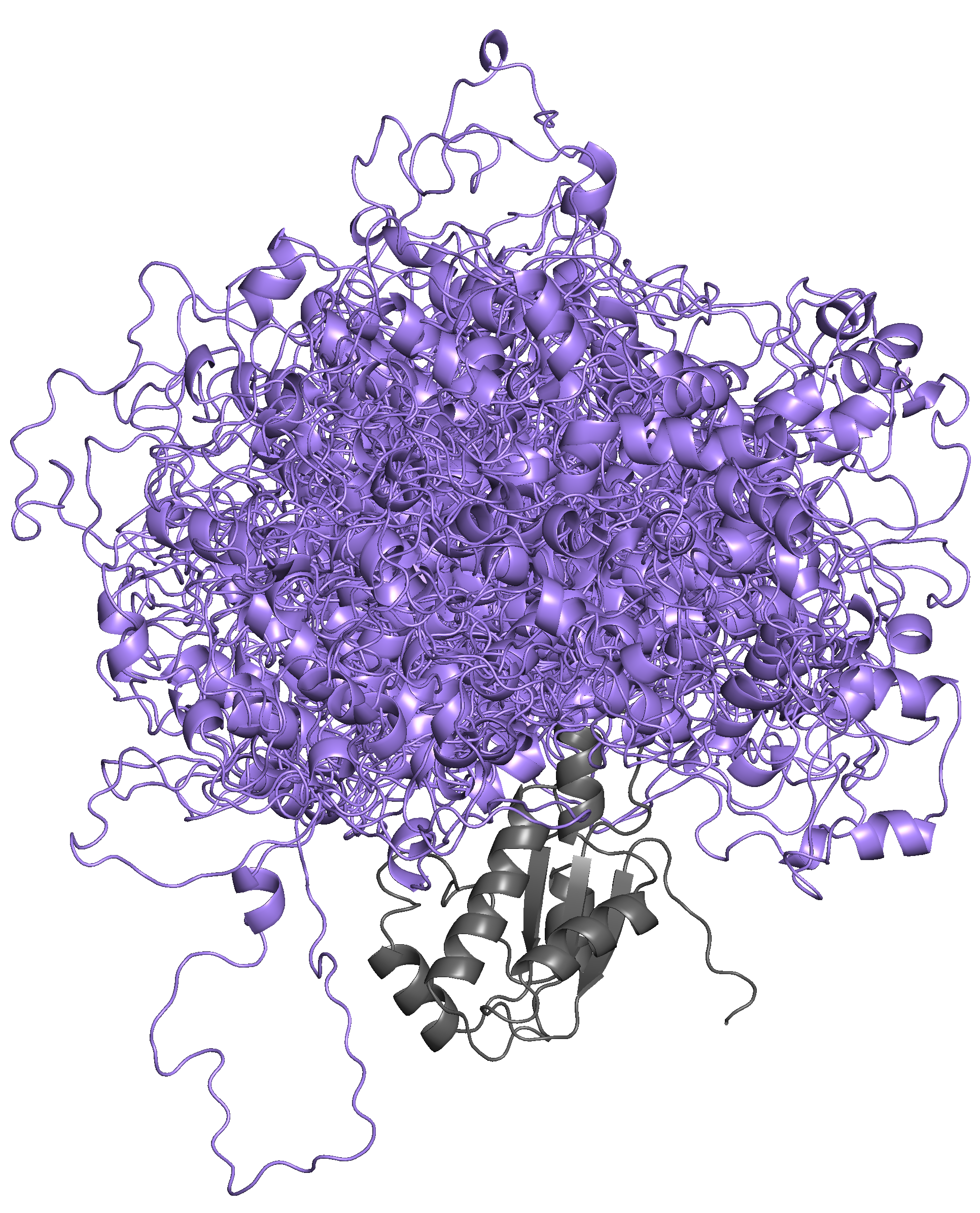

### A9_STAS_rama_plot.png

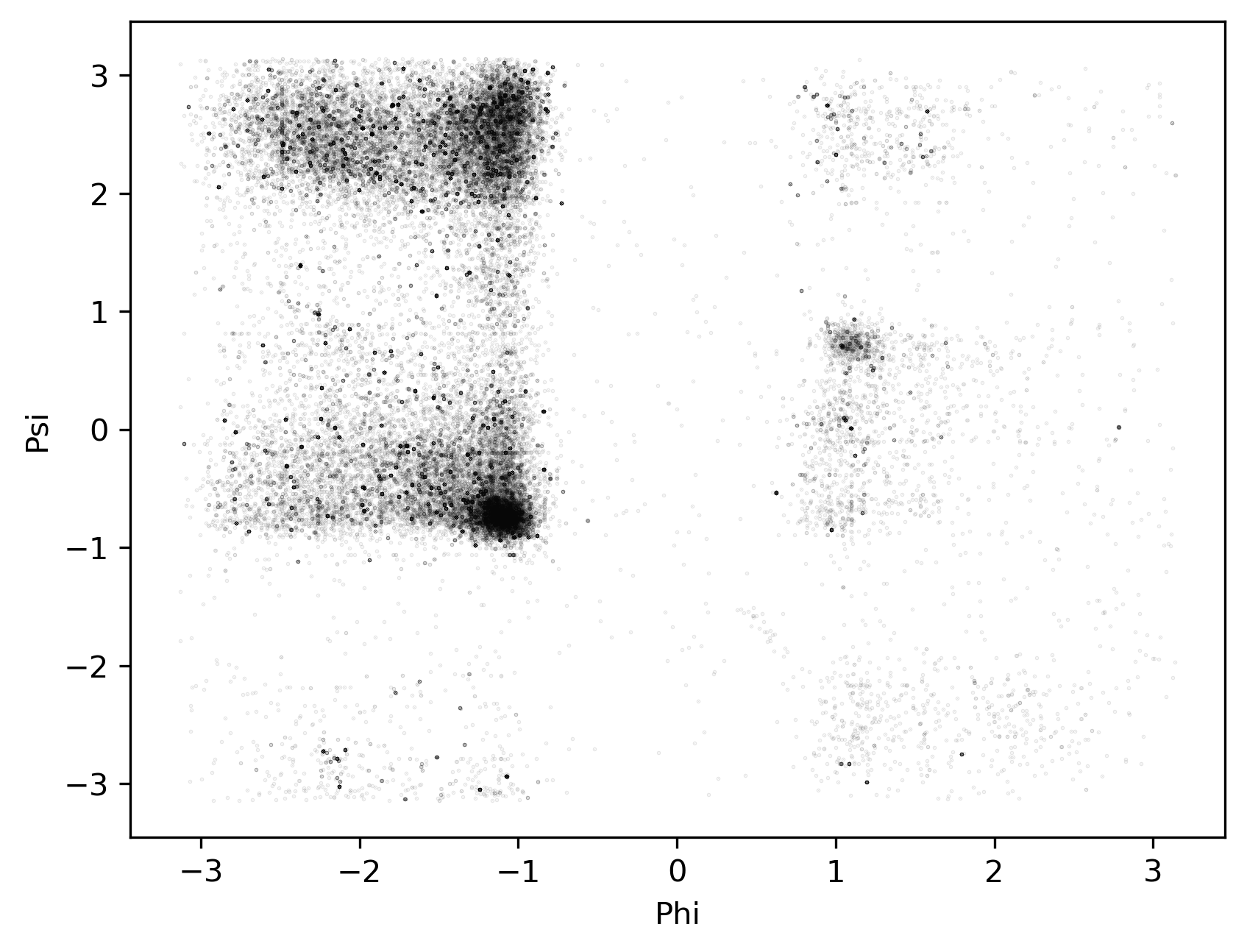

### A9_STAS_ramaSS.png

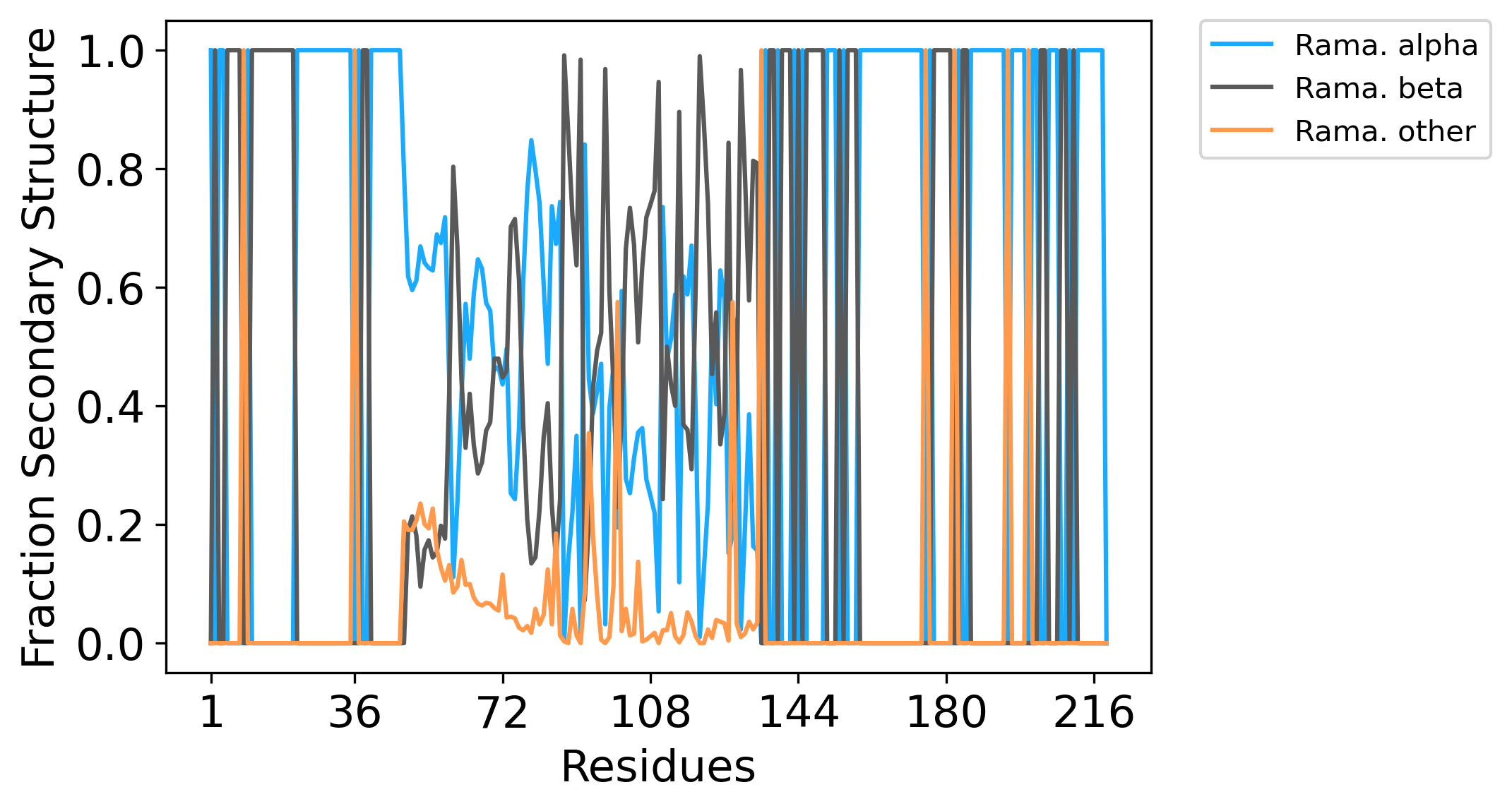

### A9_STAS_ramaSS_long.png

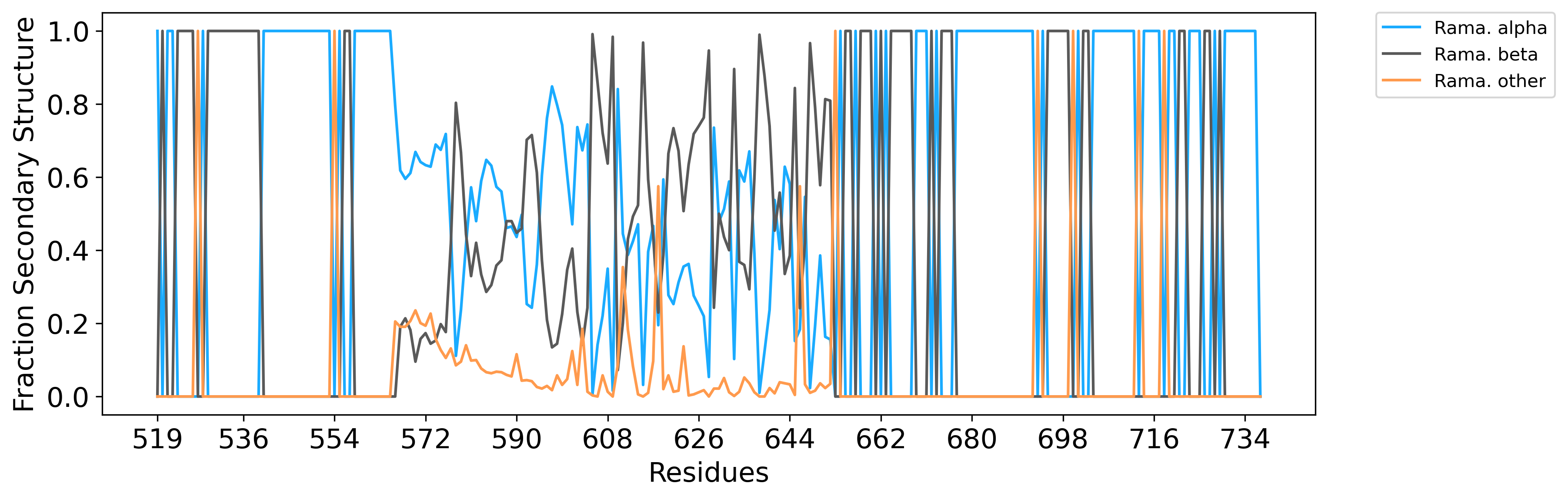

### A9_STAS_template.png

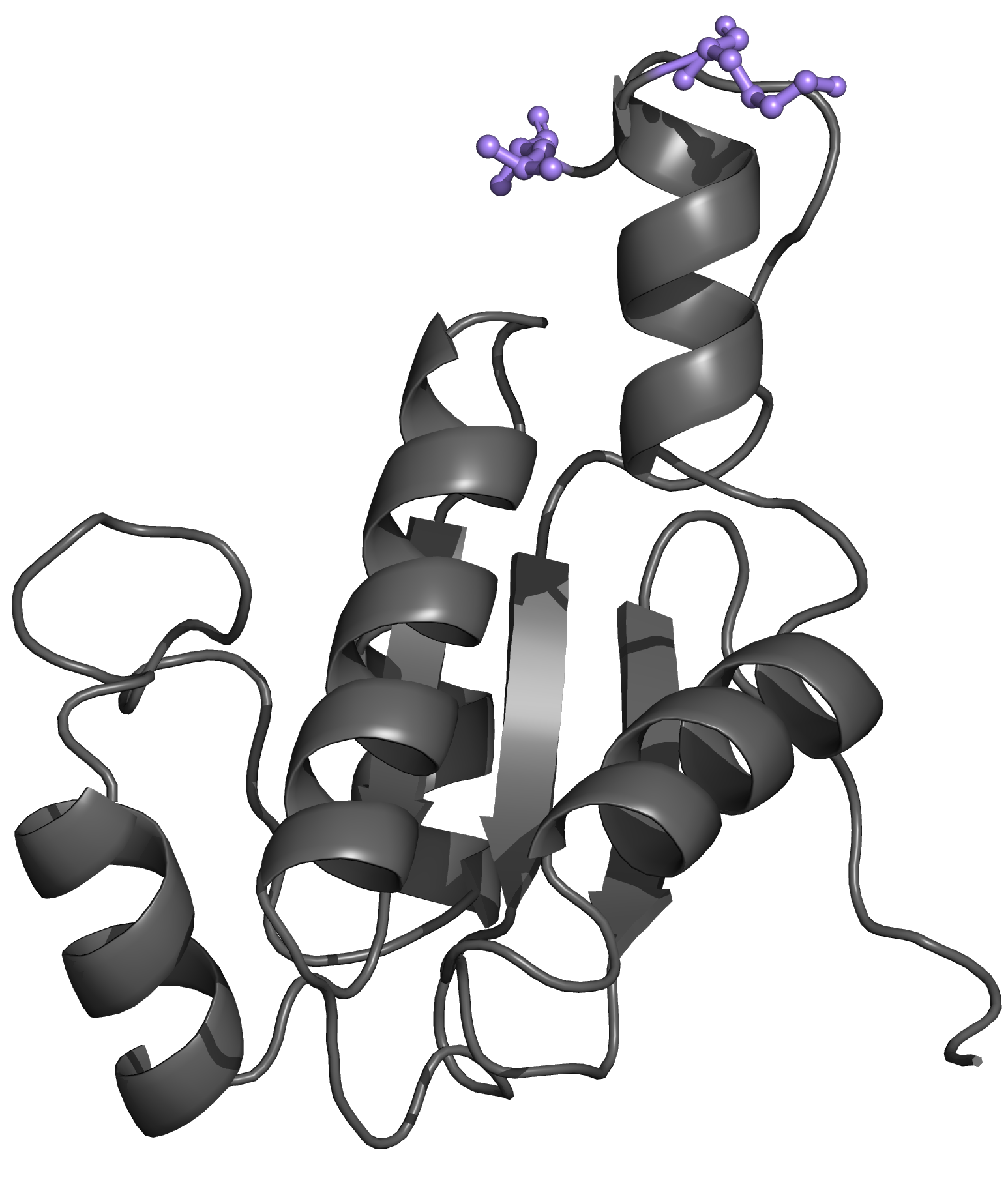

### AF-P04637-F1.png

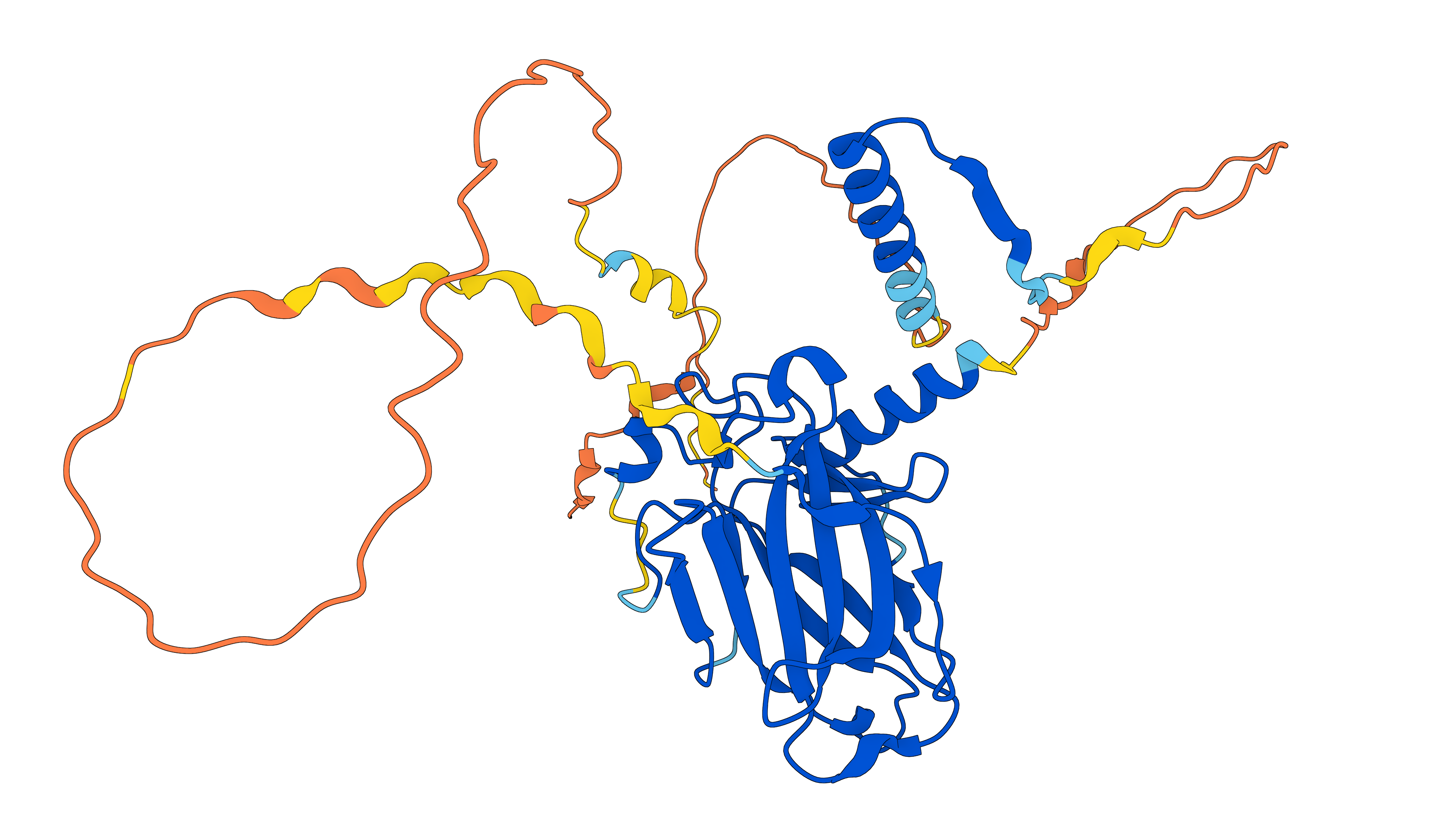

### AF-P08913-F1.png

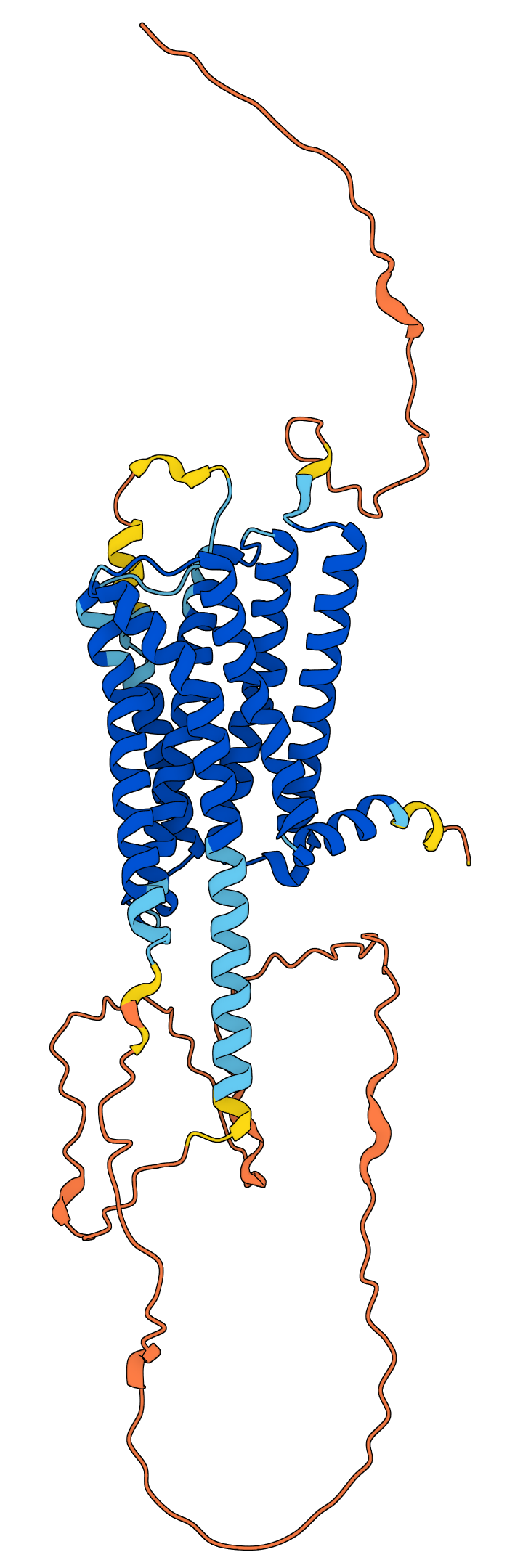

### BP2_fracSS.png

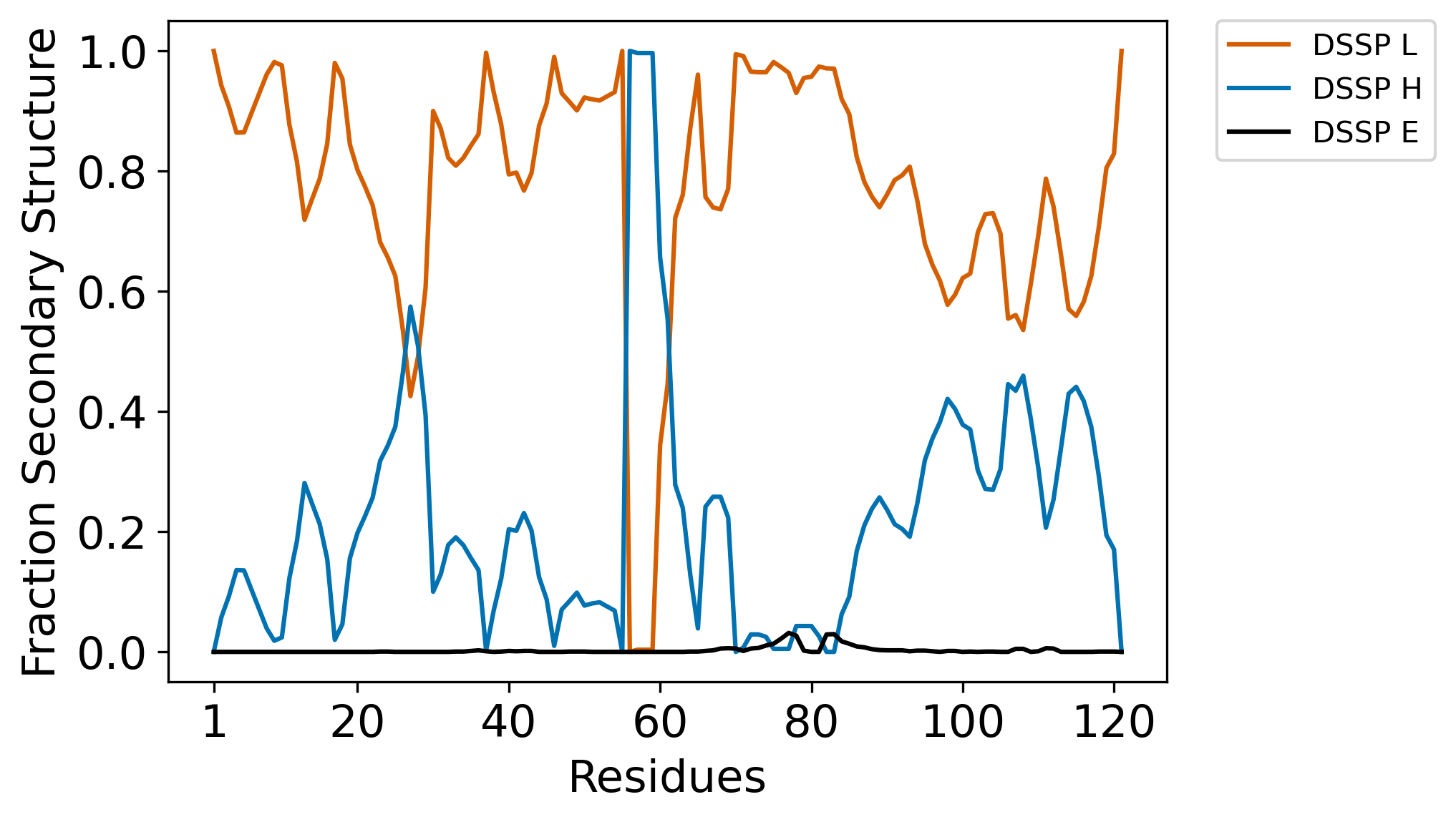

### BP2_fracSS_long.png

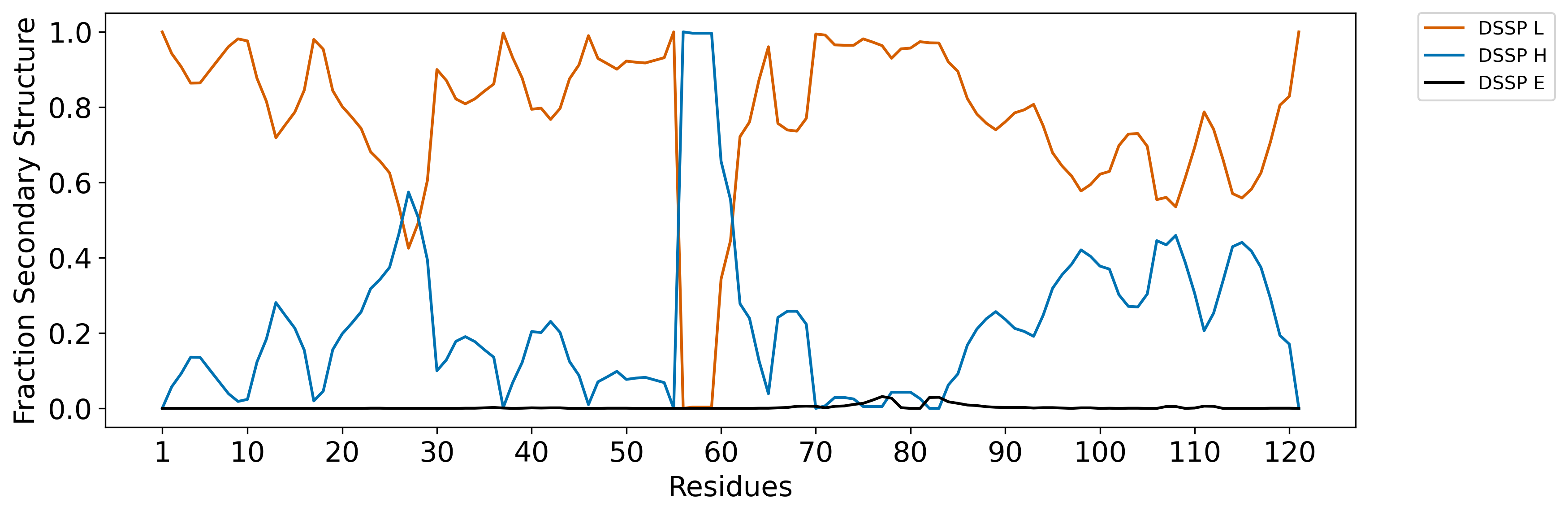

### BP2_rama_plot.png

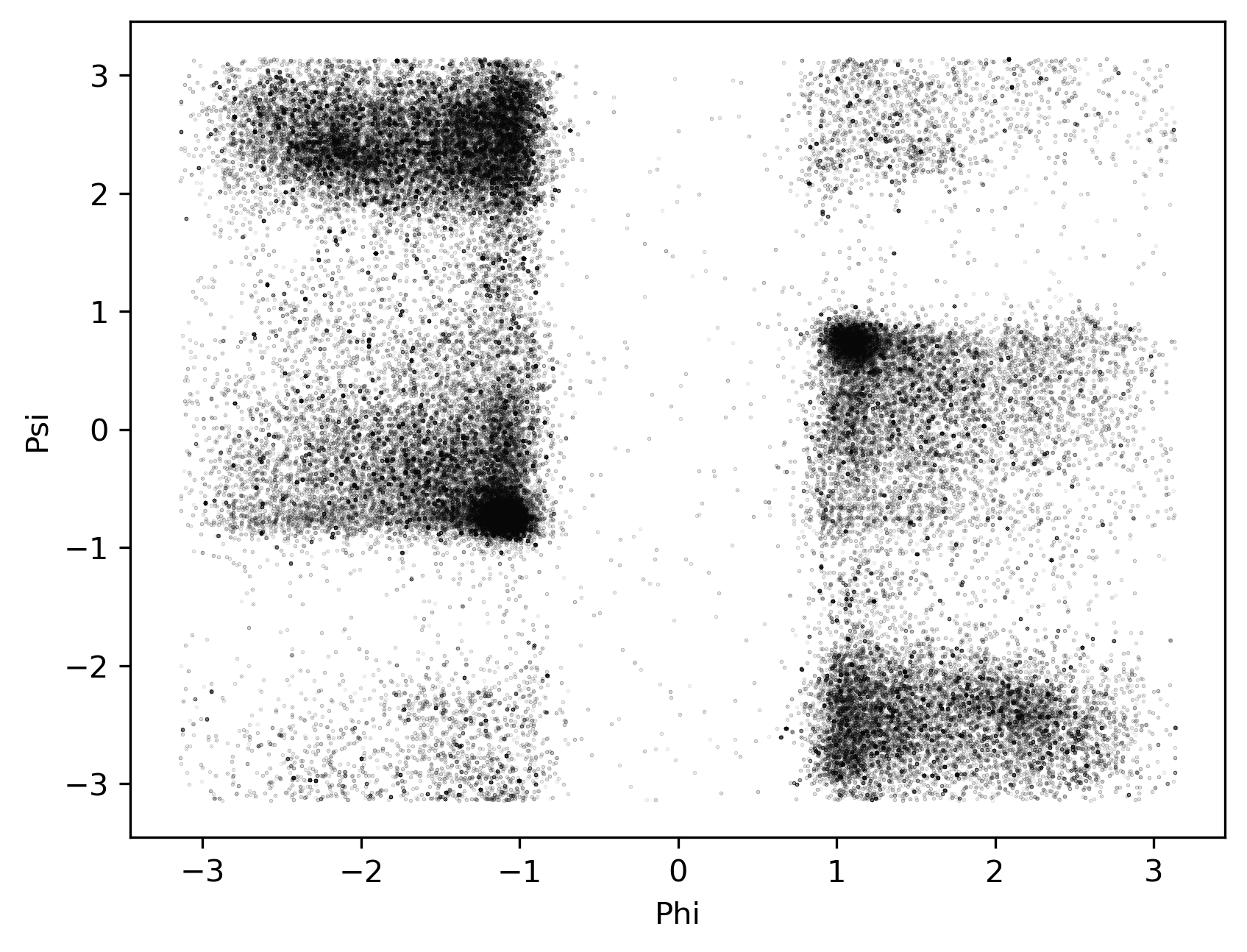

### BP2_ramaSS.png

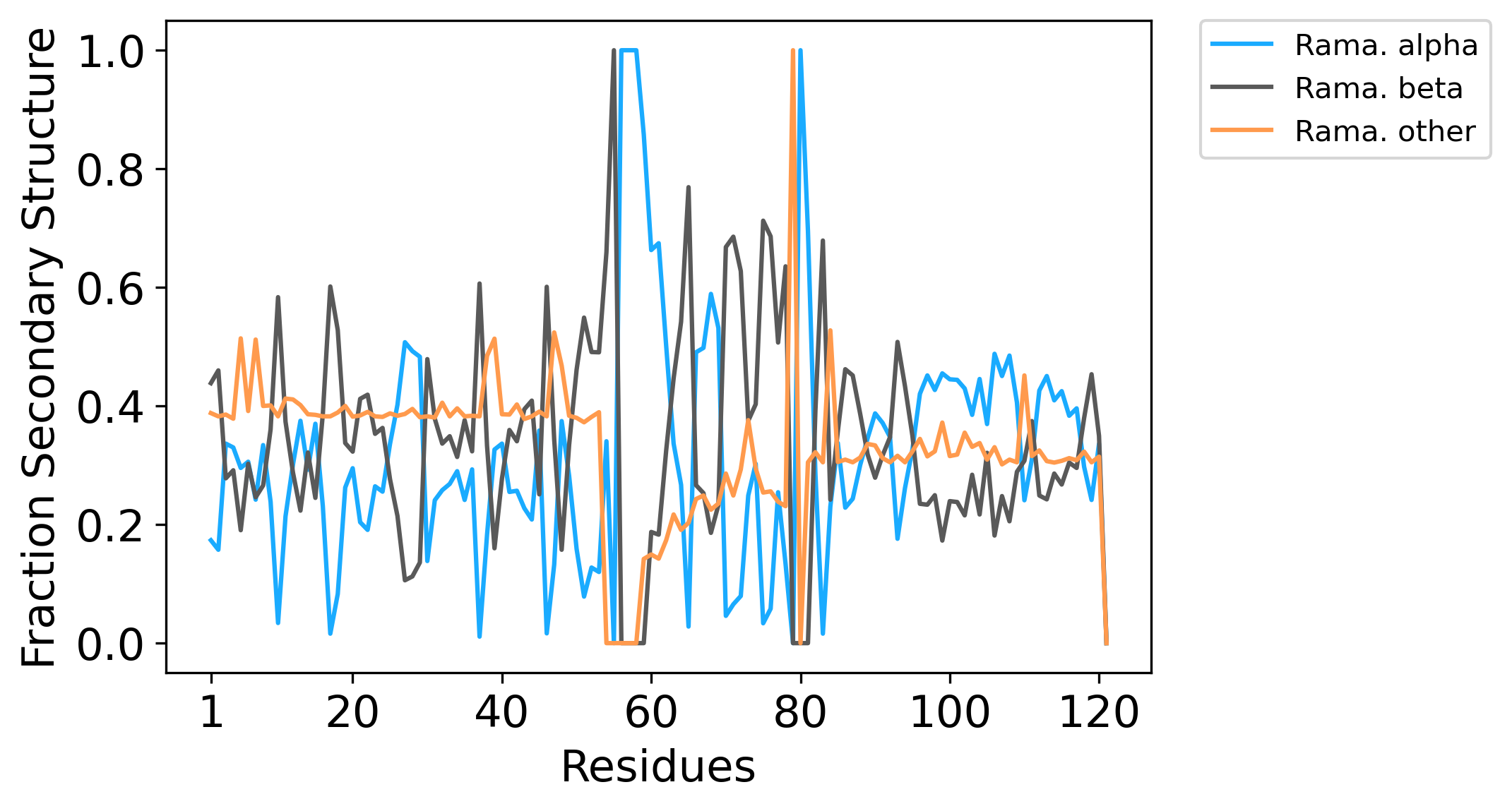

### BP2_ramaSS_long.png

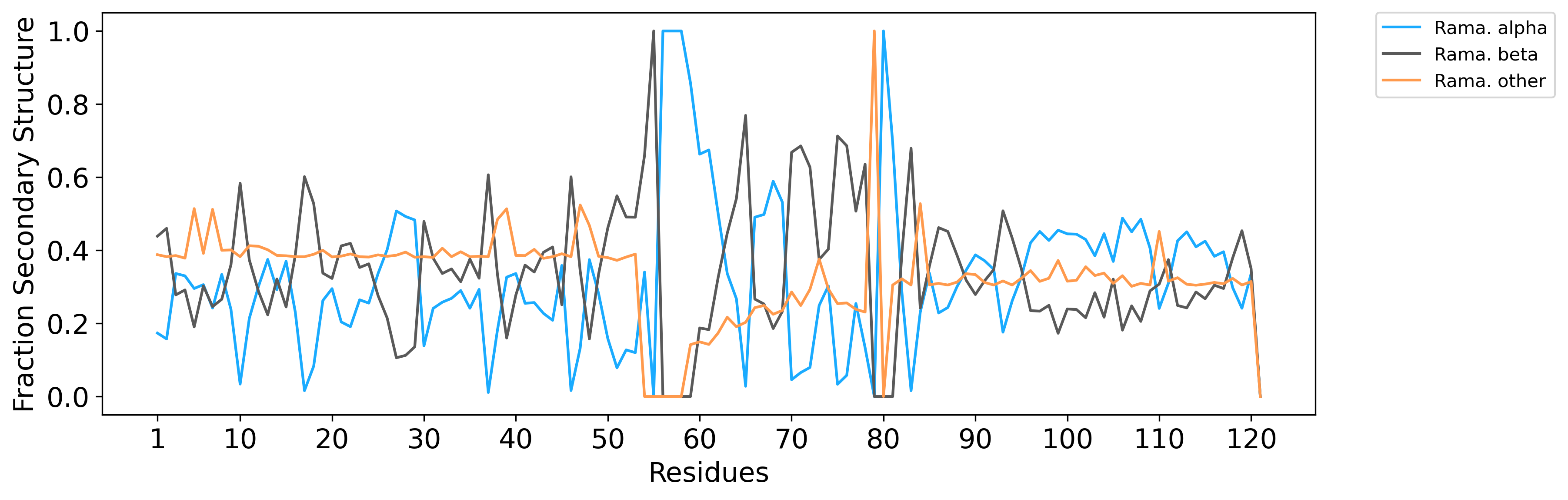

### BP24E_n100.png

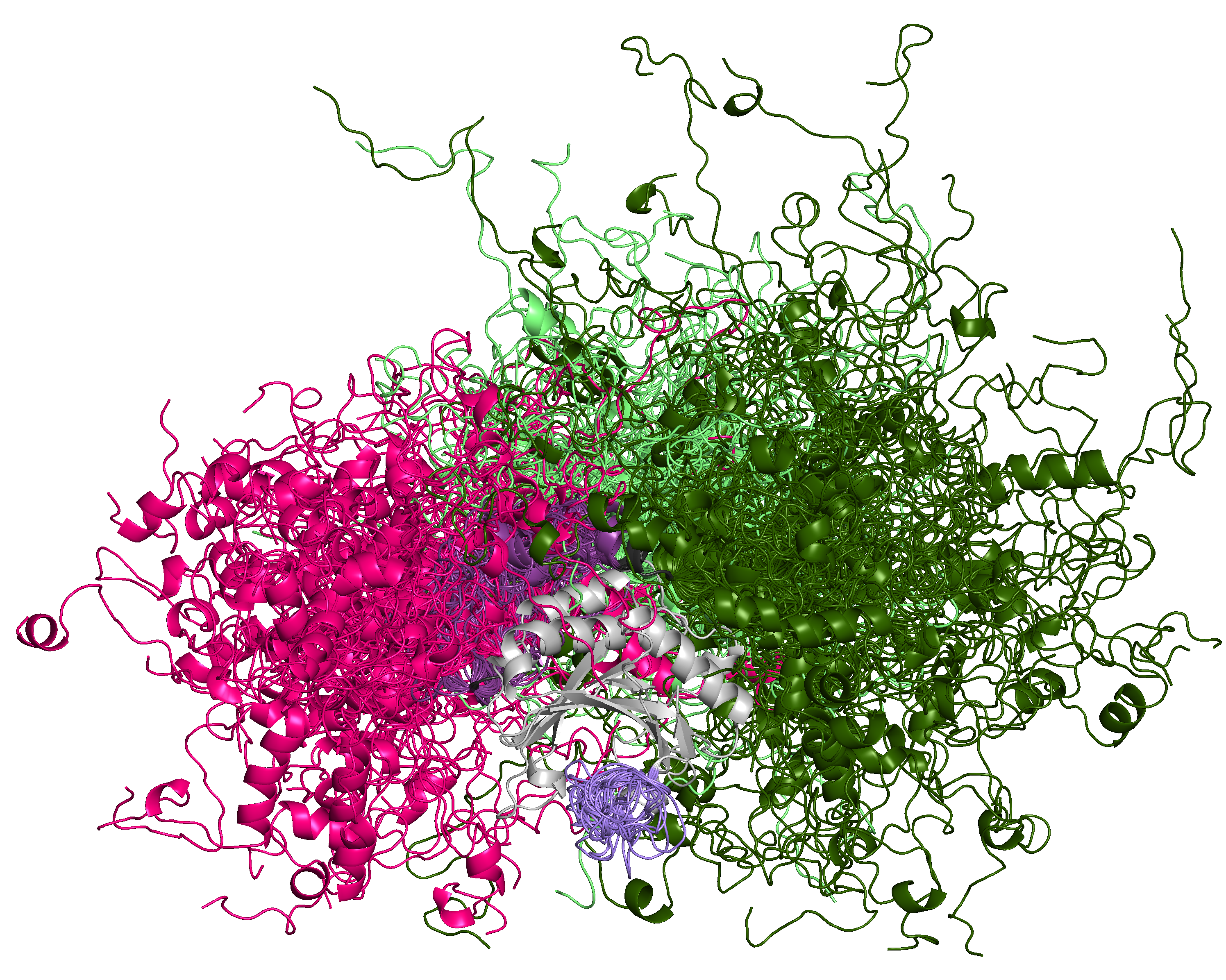

### BP24E_template.png

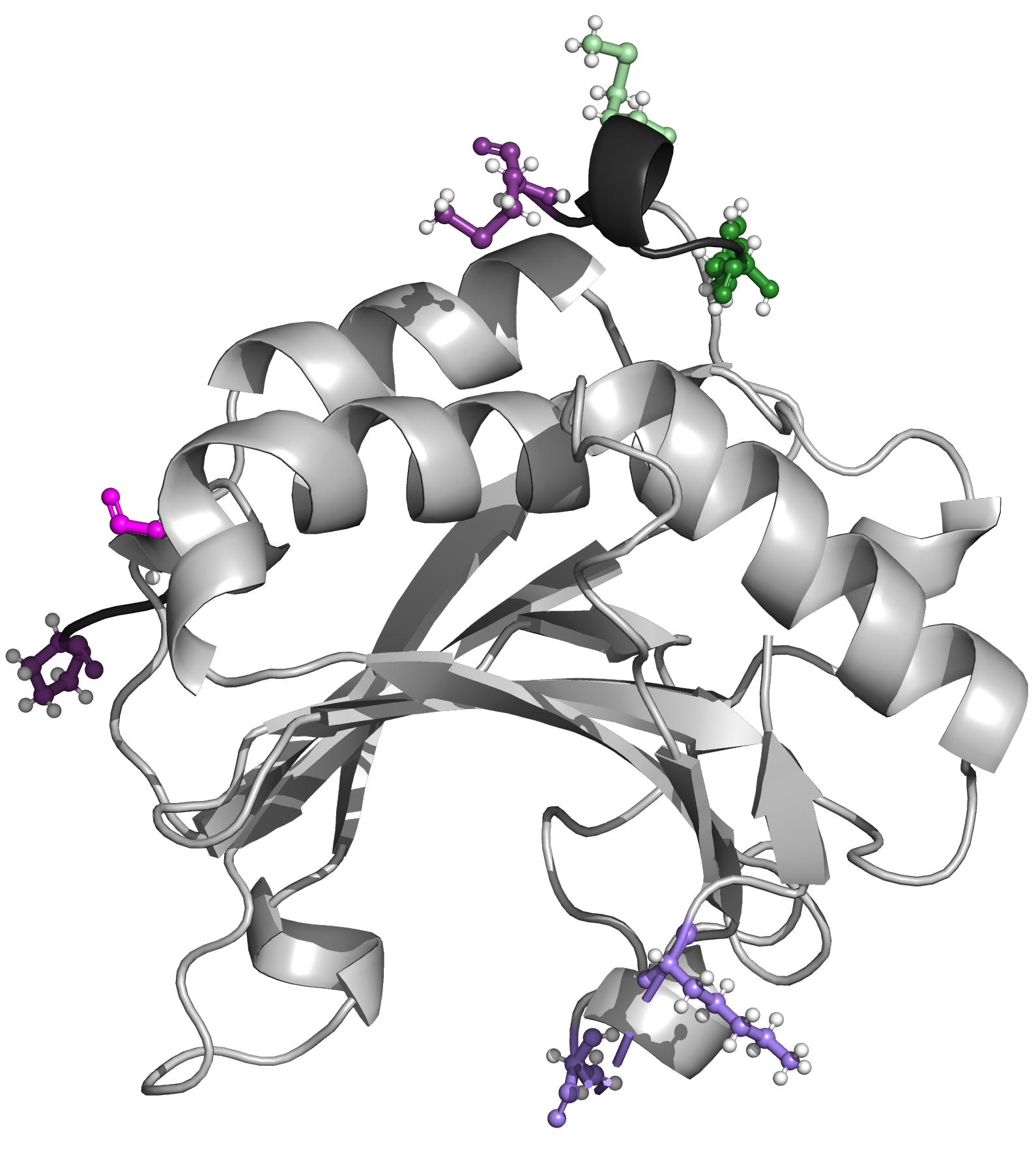

### idpcg_db_rama_plot.png

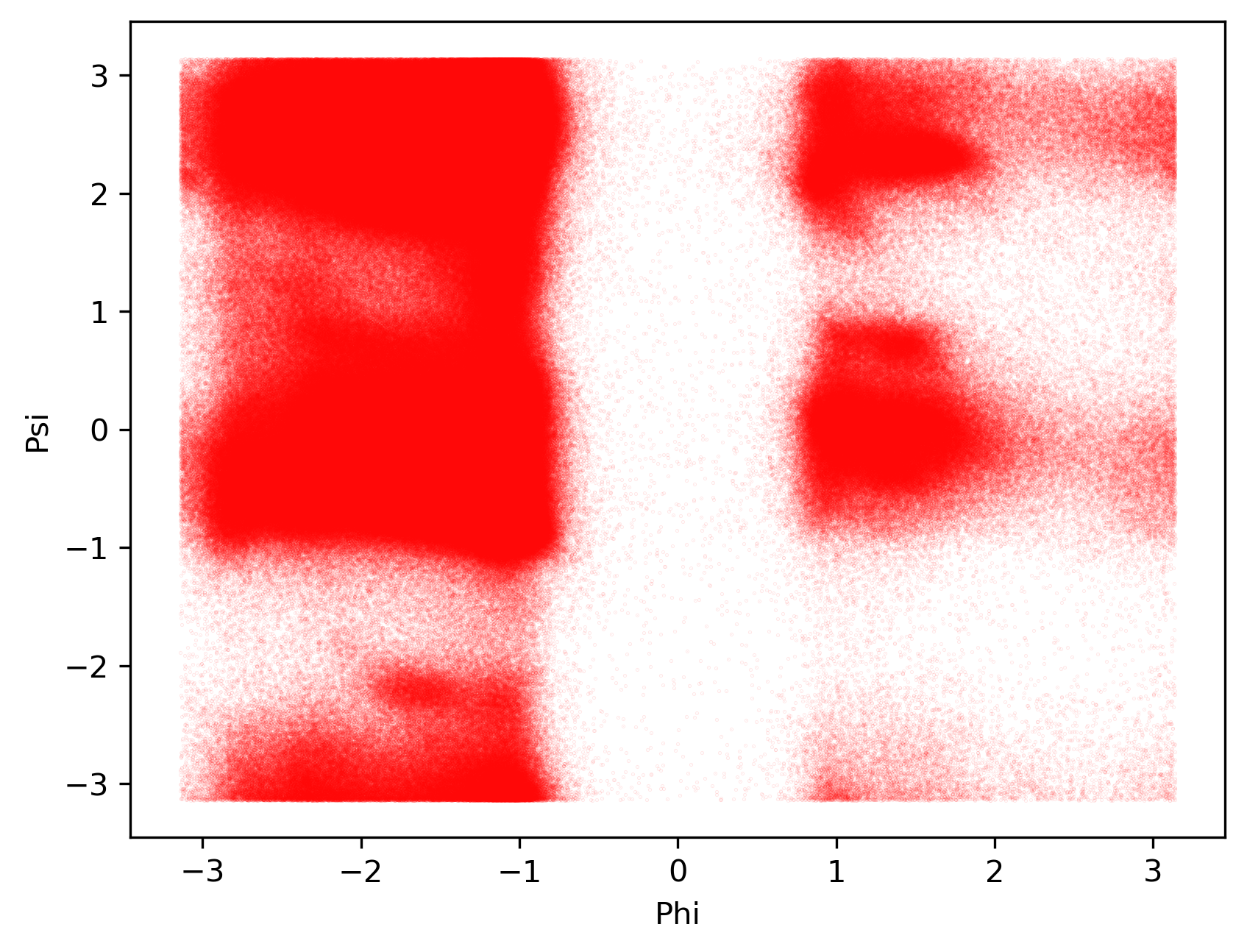

### p53_AF2_dsspSS.png

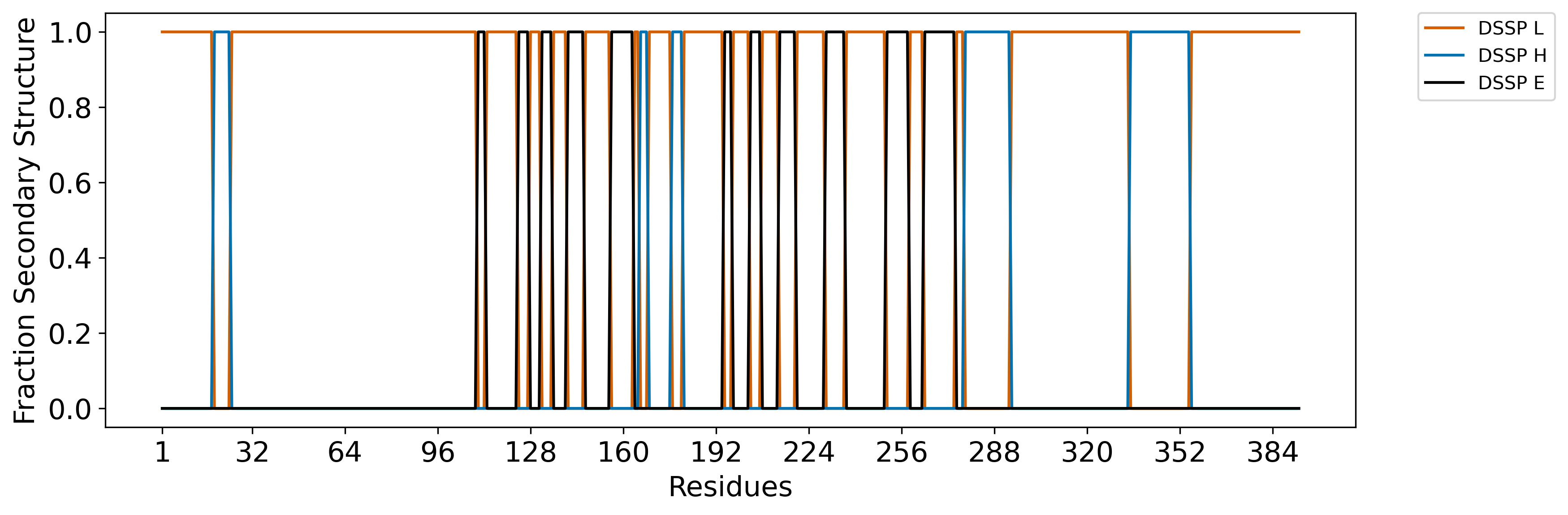

### p53_AF2_ramaSS.png

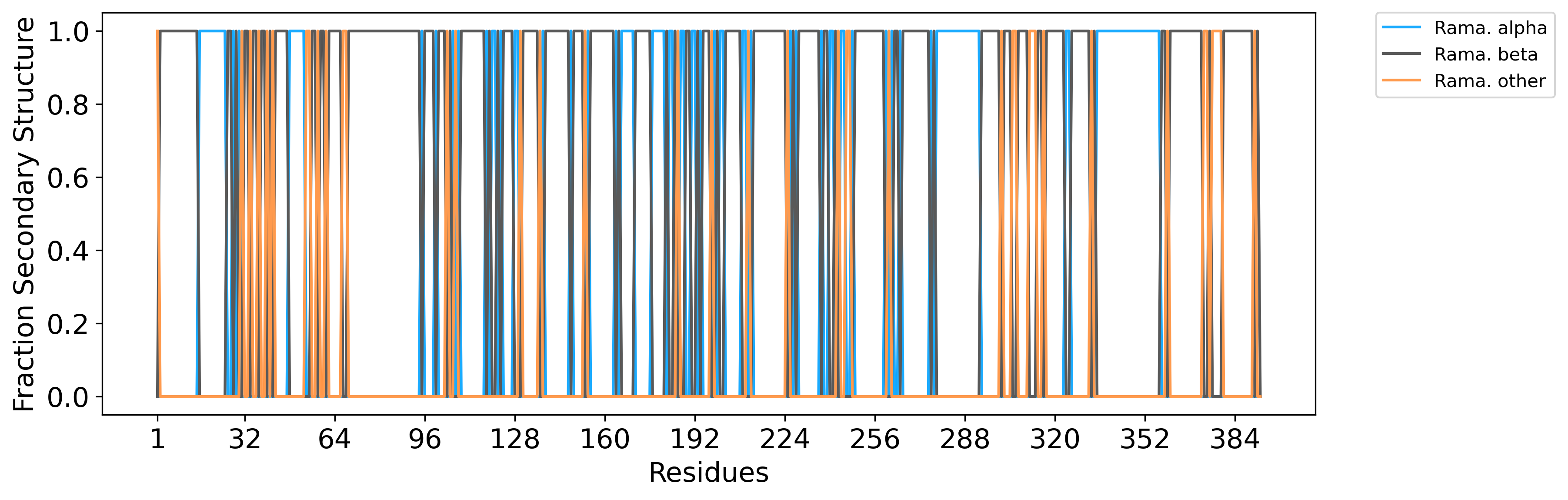

### p53_CIDR_distances.png

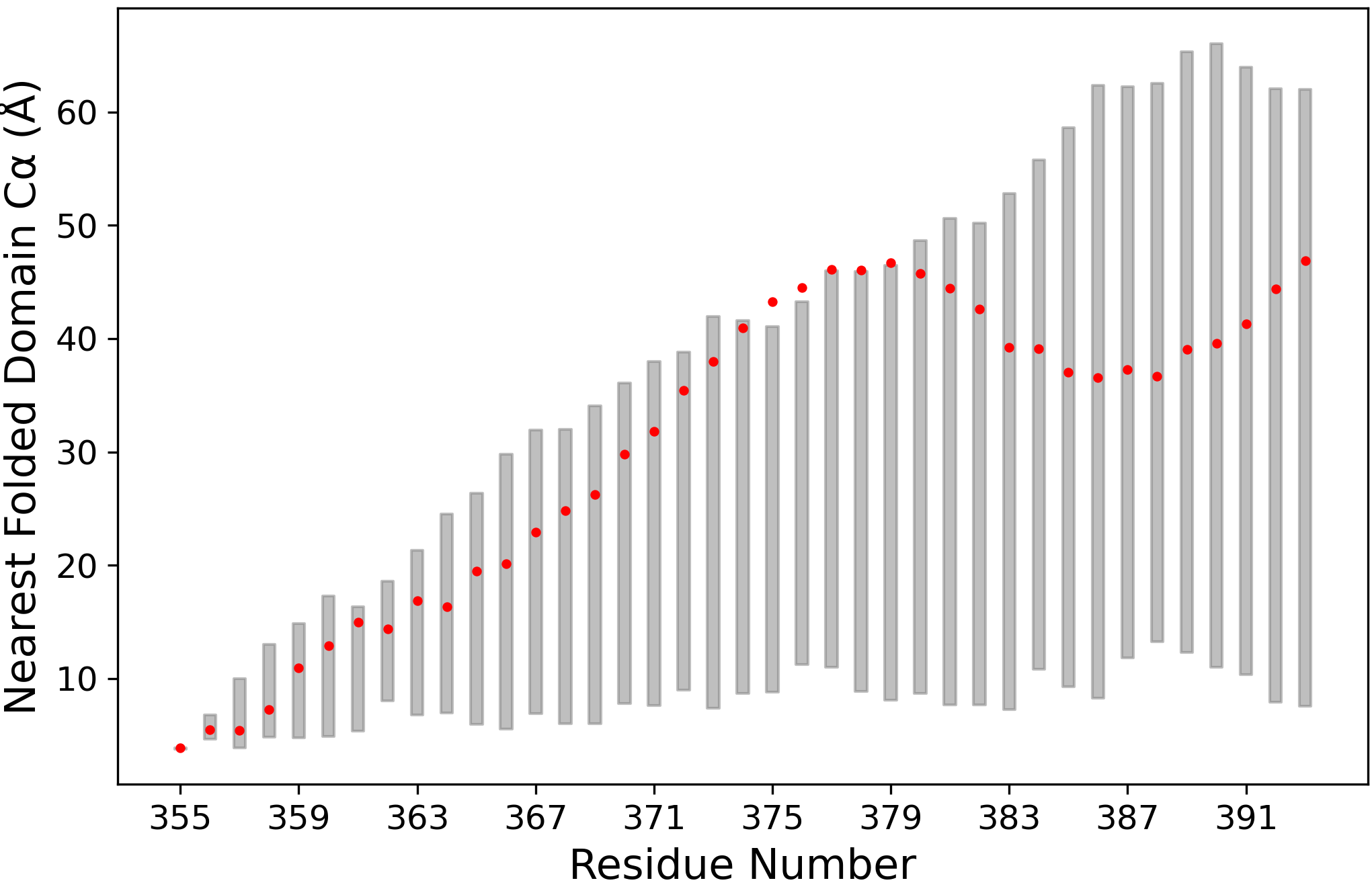

### p53_fracSS.png

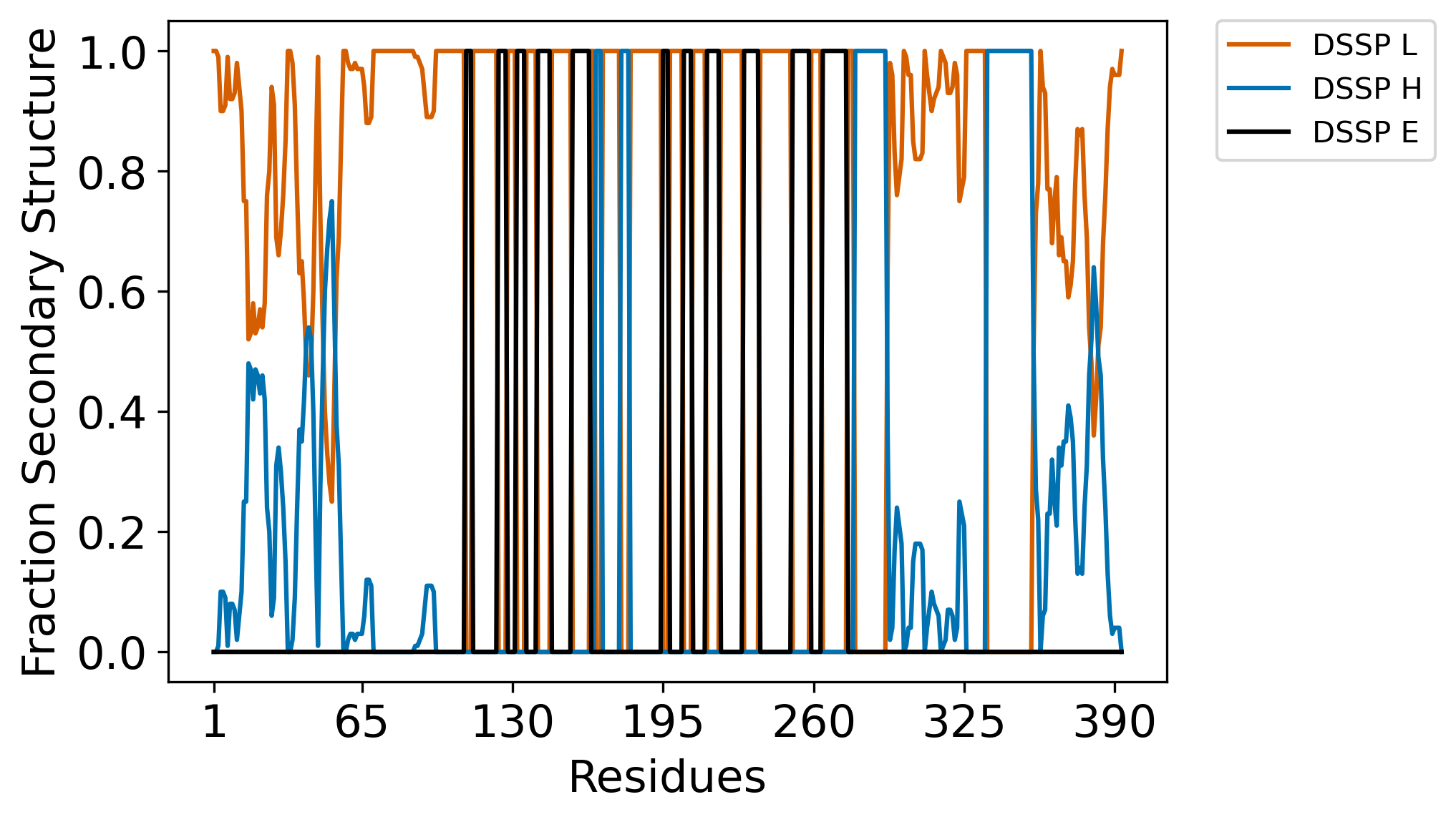

### p53_fracSS_long.png

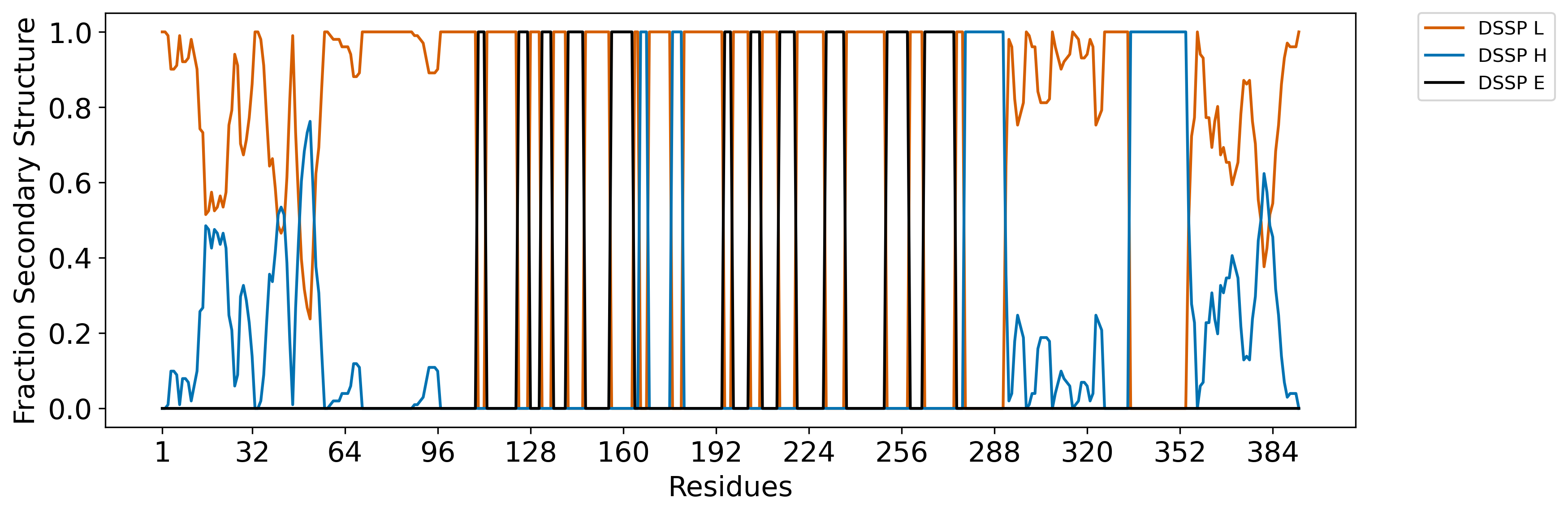

### p53_LIDR1_distances.png

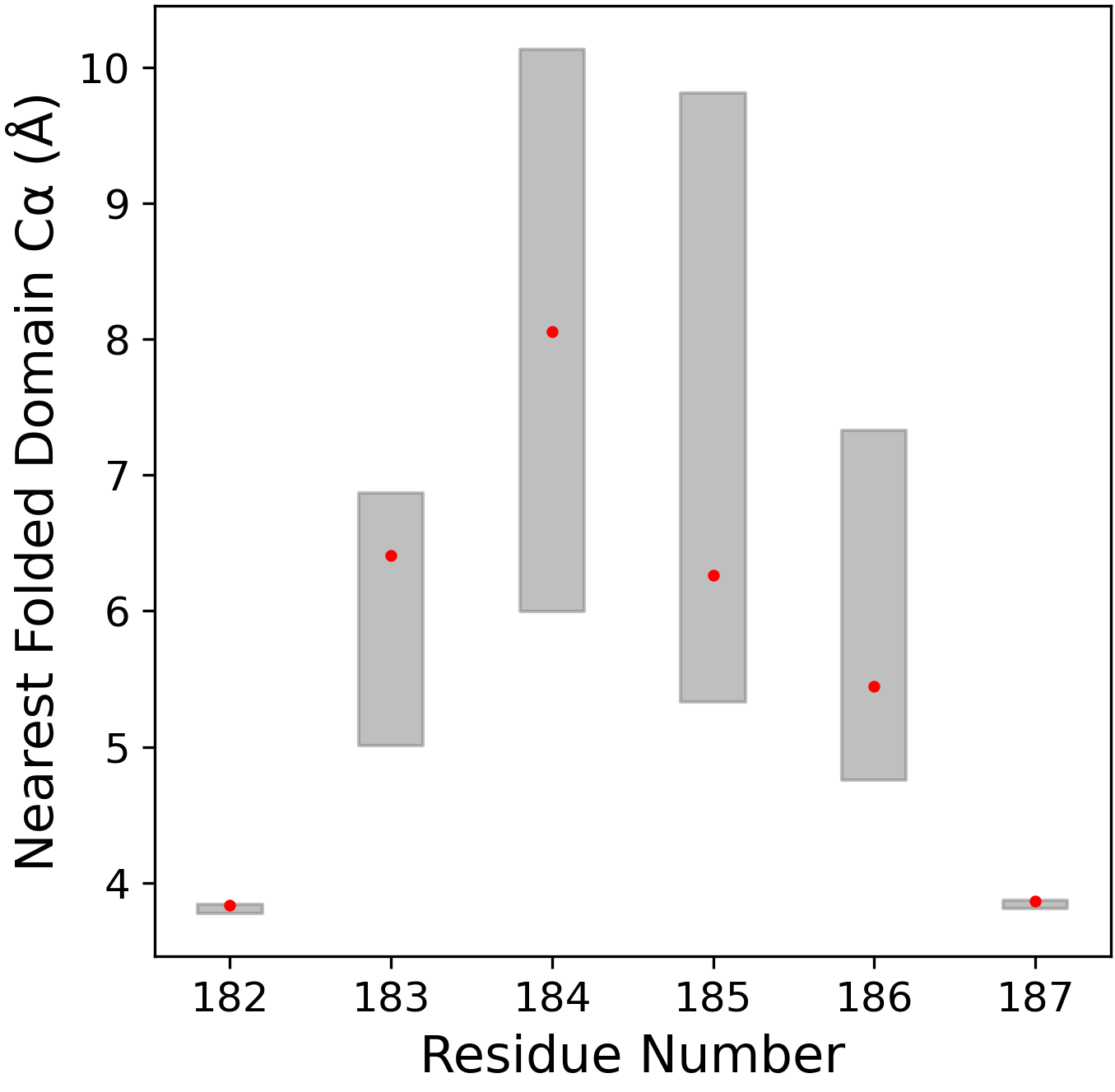

### p53_LIDR2_distances.png

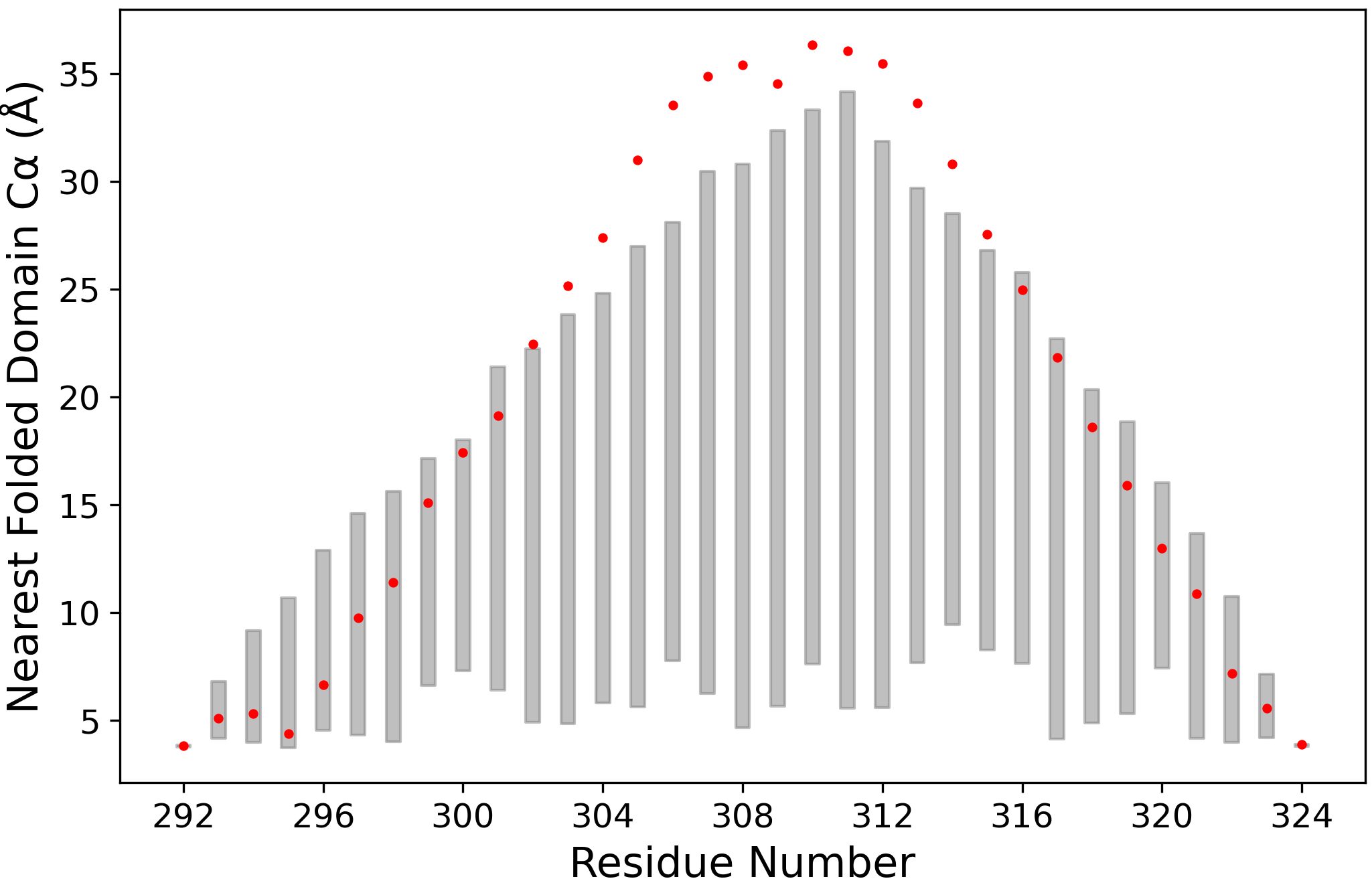

### p53_n1.png

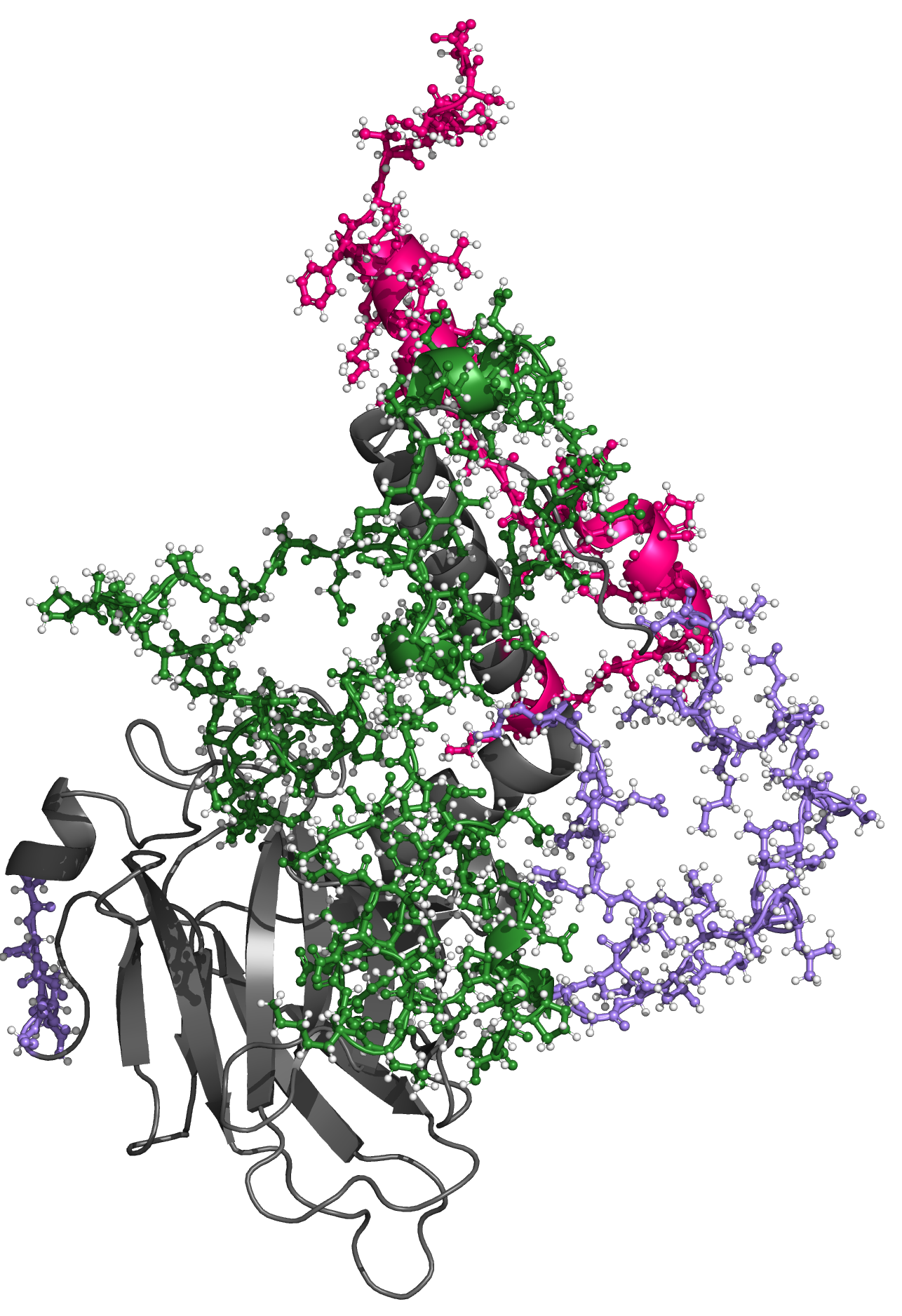

### p53_n100.png

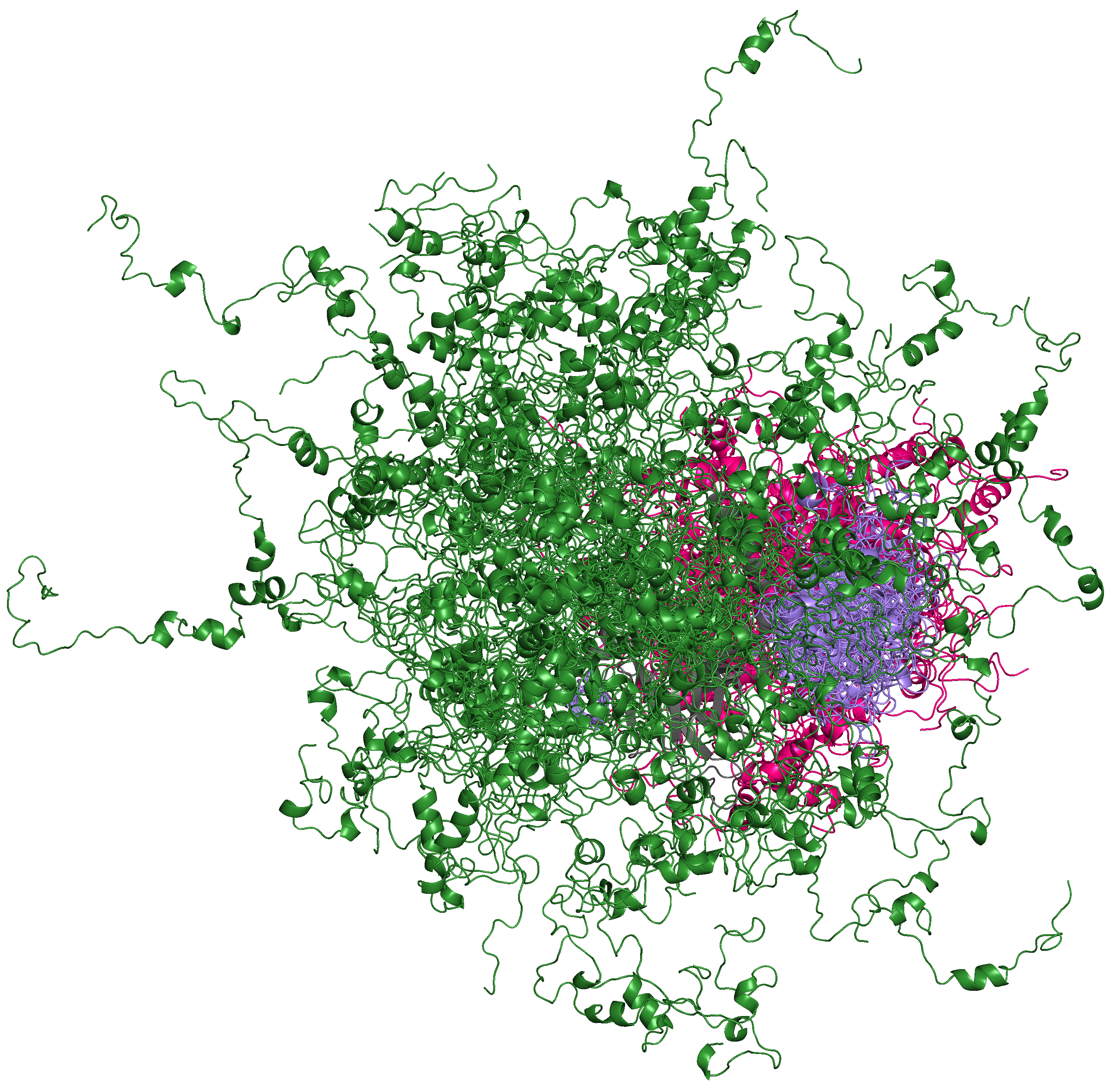

### p53_NIDR_distances.png

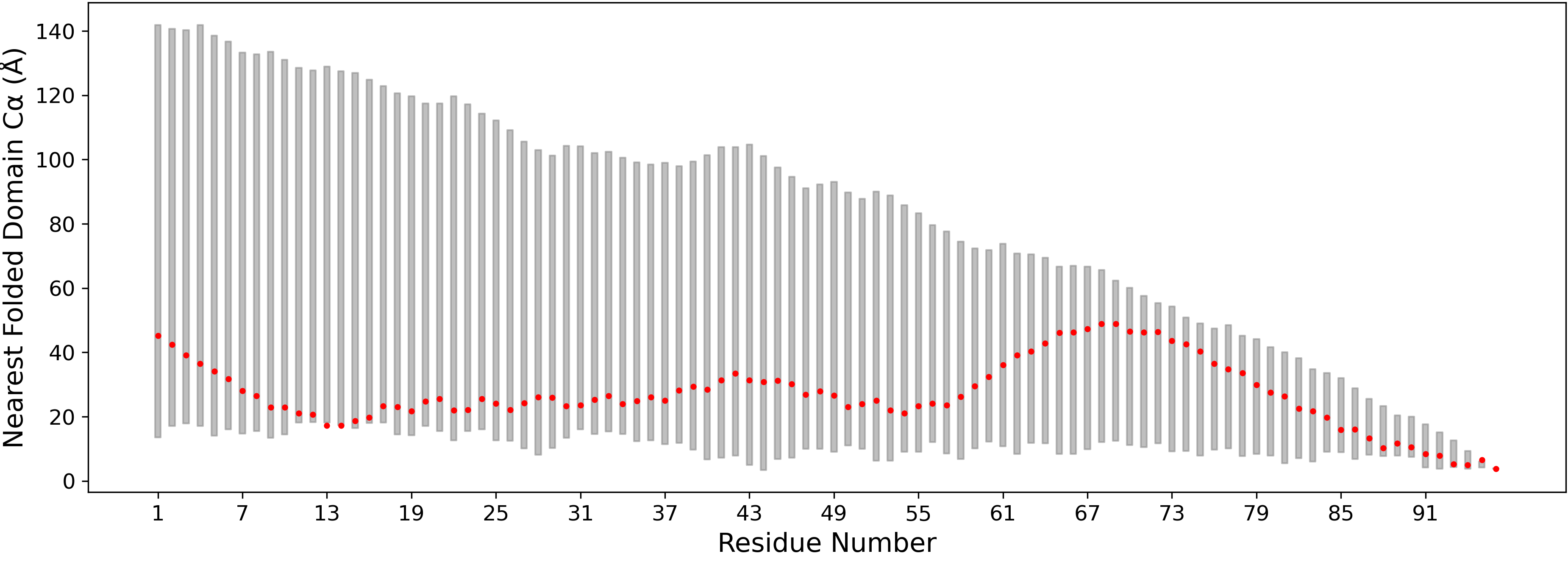

### p53_rama_plot.png

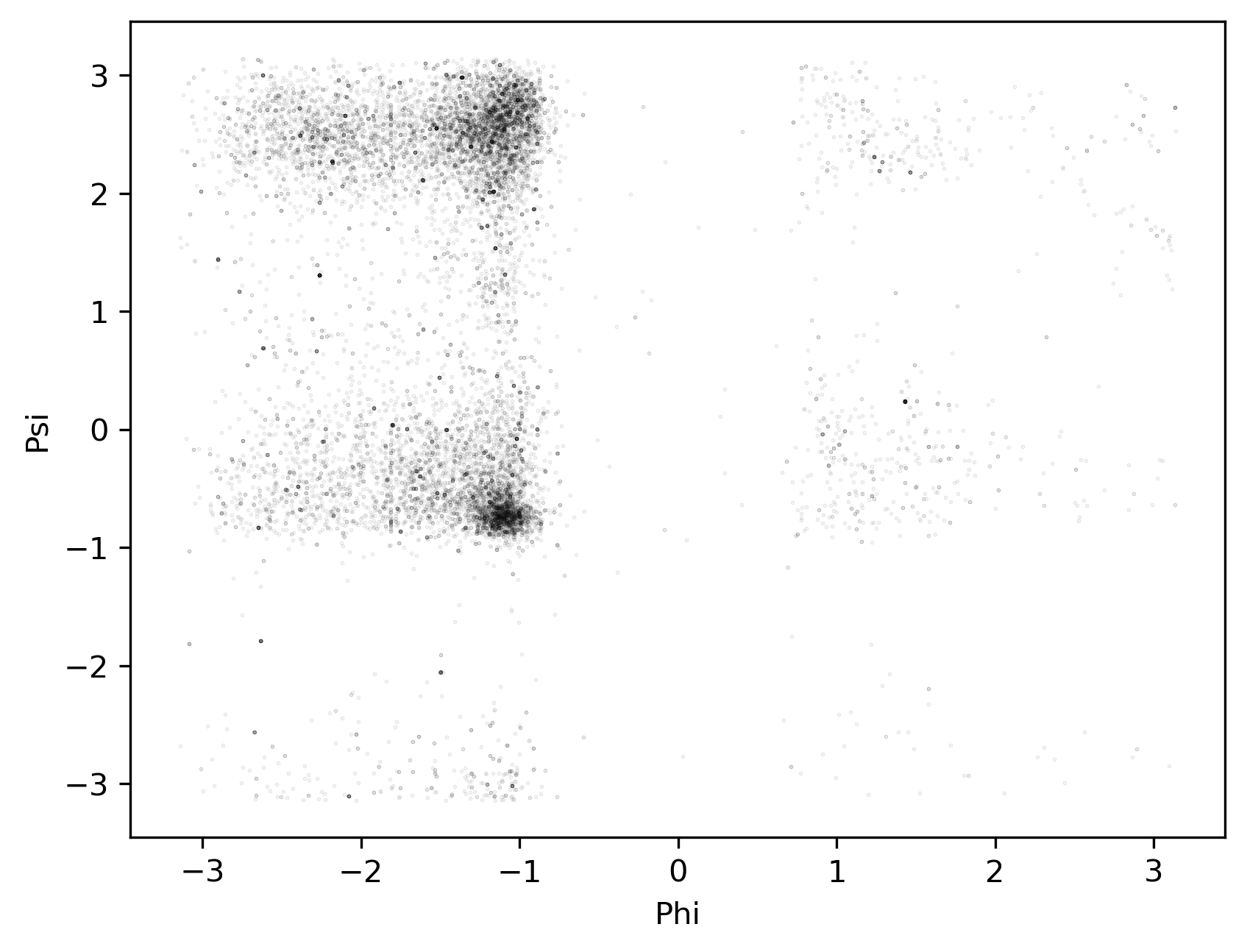

### p53_ramaSS.png

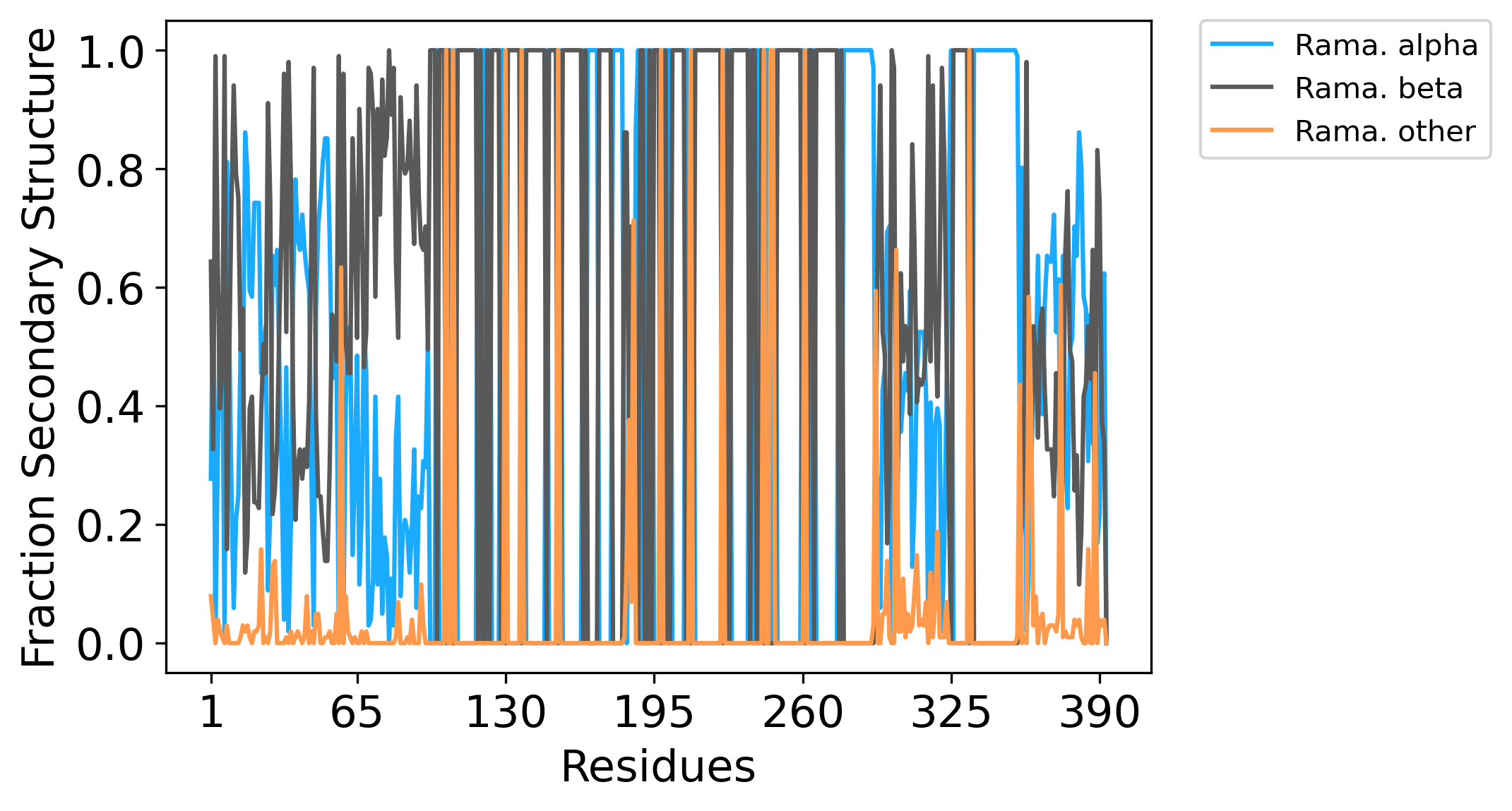

### p53_ramaSS_long.png

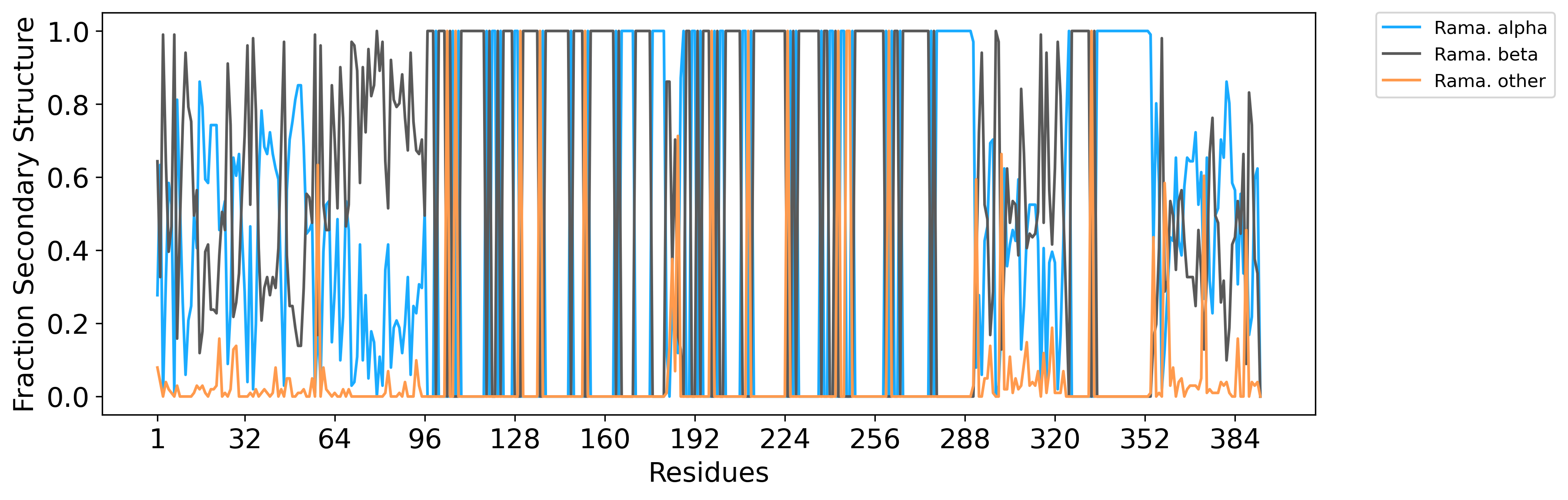
